# Oncogenic activation of Nrf2 by specific knockout of Nrf1α that acts as a dominant tumor repressor

**DOI:** 10.1101/403220

**Authors:** Lu Qiu, Meng Wang, Shaofan Hu, Xufang Ru, Yonggang Ren, Siwang Yu, Yiguo Zhang

**Affiliations:** The Laboratory of Cell Biochemistry and Topogenetic Regulation, College of Bioengineering and Faculty of Sciences, Chongqing University, No. 174 Shazheng Street, Shapingba District, Chongqing 400044, China; State Key Laboratory of Natural and Biomimetic Drugs; Department of Molecular and Cellular Pharmacology, Peking University School of Pharmaceutical Sciences, No 38 Xueyuan Rd., Haidian District, Beijing 100191, China

**Keywords:** Nrf1α, Nrf2, PTEN, Keap1, COX2, AP-1, tumor repressor, tumor promoter, regulatory networks

## Abstract

Liver-specific knockout of Nrf1 in mice leads to non-alcoholic steatohepatitis with dyslipidemia, and its deterioration results in spontaneous hepatoma, but the underlying mechanism remains elusive. A similar pathological model is herein reconstructed by using human Nrf1α-specific knockout cell lines. We demonstrated that a marked increase of the inflammation marker COX2 in *Nrf1α^−/−^* cells. Loss of Nrf1α leads to hyperactivation of Nrf2, which results from substantial decreases in both Keap1 and PTEN in *Nrf1α^−/−^* cells. Further investigation of xenograft mice showed that malignant growth of *Nrf1α^−/−^*-derived tumor is almost abolished by silencing Nrf2, while *Nrf1α^+/+^*-tumor is markedly repressed by inactive *Nrf2^−/−ΔTA^*, but unaffected by *a priori* constitutive activator of *caNrf2^ΔN^*. Mechanistic studies unraveled there exist opposing and unifying inter-regulatory cross-talks between Nrf1 and Nrf2. Collectively, Nrf1α manifests a dominant tumor-suppressive effect by confining Nrf2 oncogenicity, while Nrf2 can directly activate the transcriptional expression of *Nrf1* to form a negative feedback loop.

**HIGHLIGHTS:**

- Opposing and unifying inter-regulatory cross-talks between Nrf1α and Nrf2
- Malignant growth of *Nrf1α^−/−^*-derived tumor is prevented by silencing Nrf2
- Hyper-activation of Nrf2 by *Nrf1α^−/−^* results from decreased Keap1 and PTEN
- *Nrf1α^+/+^*-tumor is repressed by *Nrf2^−/−ΔTA^*, but unaltered by its active *caNrf2^ΔN^*

## INTRODUCTION

The steady-state lipid levels are crucial for maintaining cellular and organismal homeostasis, not only in term of energy metabolism, but also to prevent potential cytotoxity. Conversely, excessive nutrients (and metabolic stress) can culminate in a series of severe diseases such as diabetes, obesity and fatty liver. Notably, non-alcoholic fatty liver disease (NAFLD) affects 25% of global population, up to 80% of obese people having this disease (Corte et al., 2015; Younossi et al., 2018). NAFLD comprises a continuum series of pathological conditions varying in severity of liver injury and exacerbation. Among them, non-alcoholic steatohepatitis (NASH) is defined as a serious process with inflammation and hepatocyte damage, and also hence regarded as a major cause of liver fibrosis, cirrhosis, and even cancer, such as hepatocellular carcinoma (HCC), among those caused by unknown etiologies (Friedman et al., 2018; Michelotti et al., 2013; Rowell and Anstee, 2015; Wong et al., 2018). However, the axiomatic mechanisms underlying development of NASH and malignant transformation into hepatoma remain elusive.

The cumulative evidence obtained from distinct animal models resembling human NASH (Friedman et al., 2018) demonstrates that homeostatic and nutrient-stimulated lipid metabolisms are tightly regulated by multiple layers of diverse signaling to transcription factor networks to monitor precision expression of target genes (Jump et al., 2013; Karagianni and Talianidis, 2015). Among them, SREBP1c (sterol-regulatory element binding protein 1c) is established as a key marker and therapeutic target for hepatosteatosis, because transgenic over-expression of this factor leads to hepatosteatosis, but not hepatoma (Nakayama et al., 2007). Yet, similar hyperactivation of SREBP1c by knockout of *GP78*, an endoplasmic reticulum (ER) membrane-bound E3 ligase, occurs with age-related obesity, NASH and HCC (Zhang et al., 2015a). Conversely, hepatosteatosis is partially mitigated by the deficiency of SREBP1c (Yahagi et al., 2002), but sufficiently ameliorated by blockage of SREBP processing by deletion of *SCAP* (SREBP cleavage-activating protein) (Moon et al., 2012). These findings indicate an additive involvement of other factors beyond SREBPs in NASH-associated malignant pathology.

Interestingly, spontaneous NASH with massive hepatomegaly and hepatoma results from the hepatocyte-specific knockout of *PTEN* (phosphatase and tensin homolog, a well-known tumor repressor) in mice (Horie et al., 2004). Loss of PTEN leads to constitutive activation of the phosphatidylinositol 3-kinases (PI3K)-AKT-mTOR pathway so as to augment expression of metabolic genes regulated by SREBP1c and PPARγ in cancer proliferative cells (Hollander et al., 2011; Lee et al., 2018; Shimano and Sato, 2017). This process is accompanied by nuclear accumulation of Nrf2 (nuclear factor erythroid 2-like 2, also called NFE2L2) in *PTEN*-deficient cells (Mitsuishi et al., 2012; Sakamoto et al., 2009). Nrf2 and Nrf1 are two principal members of the cap’n’collar (CNC) basic-region leucine zipper (bZIP) family to transactivate antioxidant response element (ARE)-driven genes involved in detoxification, cytoprotection, metabolism and proliferation. Significantly, aberrant accumulation of Nrf2 and activation of target genes are significantly incremented by simultaneous deletion of *PTEN* (leading to a GSK3β-directed phosphodegron of Nrf2 targeting this CNC-bZIP protein to the β-TrCP-based E3 ubiquitin ligase Cullin 1-mediated proteasomal degradation) and Keap1 (acting as an adaptor targeting Nrf2 to the Cullin 3-mediated proteasomal degradation), resulting in a deterioration of *PTEN^−/−^*-leading cancer pathology (Best et al., 2018; Rojo et al., 2014; Taguchi et al., 2014). Conversely, malignant transformation of double *PTEN:Keap1* knockout mice is alleviated by additive deletion of Nrf2 (Taguchi et al., 2014), implying Nrf2 promotes carcinogenesis. This is consistent with further observations that activity of Nrf2 is required for oncogenic KRAS-driven tumorigenesis (DeNicola et al., 2011) and its activation by antidiabetic agents accelerates tumor metastasis in xenograft models (Wang et al., 2016). However, non-neoplastic lesions are caused by constitutive active Nrf2 (caNrf2) mutants lacking the Keap1-binding sites in transgenic mice (Schafer et al., 2010; Shanmugam et al., 2017), but their cytoprotection against carcinogenesis is enhanced. Further investigation of a dominant-negative dnNrf2 mutant (also suppresses other CNC-bZIP factors, such as Nrf1) has demonstrated that the basal ARE-driven gene expression, but not their inducible expression, is crucial for anti-tumor chemoprevention against the chemical-induced carcinogenesis (auf dem Keller et al., 2006). Yet, the underlying mechanism by which Nrf2 is determined to exert dual opposing roles in tumor suppression or promotion remains unknown to date.

Significantly, another phenotype of spontaneous NASH and hepatoma is manifested in conditional *Nrf1^−/−^* mice, displaying a bulk of lipid drops in the ER with dramatic morphological changes (Ohtsuji et al., 2008; Xu et al., 2005). Global knockout mice of *Nrf1^−/−^* die of severe oxidative stress-induced damages and fetal liver hypoplasia during development (Chan et al., 1998; Chen et al., 2003). By contrast, global *Nrf2^−/−^* knockout mice are viable and fertile, without any obvious pathological phenotypes occurring during normal growth and development (Itoh et al., 1997). Such facts indicate that Nrf1 is not compensated by Nrf2, although both are widely co-expressed in various tissues and have similar overlapping roles in coordinately regulating ARE-driven genes. Further insights also reveal that Nrf1 exerts unique essential functions, which are distinctive from Nrf2, in maintaining cellular redox, lipid and protein homeostasis, as well as organ integrity, through regulation of distinct target genes (Bugno et al., 2015; Zhang and Xiang, 2016). This is reinforced by further investigation of other organ-specific *Nrf1* deficiency or over-activation in mice, which exhibit distinct pathological phenotypes, such as type 2 diabetes, neurodegenerative and cardiovascular disease (Hirotsu et al., 2014; Kobayashi et al., 2011; Lee et al., 2011; Zheng et al., 2015). In addition to functionality of Nrf1 as an indispensable CNC-bZIP transcription factor, it is also identified to act as a directly ER membrane-bound sensor to govern cholesterol homeostasis through the consensus recognition motifs (Widenmaier et al., 2017; Zhang et al., 2014b) and lipid distribution in distinct tissues (Bartelt et al., 2018; Hou et al., 2018). However, it is regrettable to unclearly define which isoforms of Nrf1 are required to execute its unique physio-pathological functions, because almost all isoforms of the factor are disrupted to varying extents in the above-described experimental models.

Upon translation of Nrf1, its N-terminal ER-targeting signal anchor enables the nascent full-length protein Nrf1α to be topologically integrated within and around the membranes, while other domains of the CNC-bZIP protein are partitioned on the luminal or cytoplasmic sides (Zhang et al., 2007b; Zhang et al., 2014b). Subsequently, some luminal-resident domains of Nrf1α are dynamically repositioned across membranes through a p97-driven retrotranslocation pathway (Radhakrishnan et al., 2014; Sha and Goldberg, 2014; Zhang et al., 2015b). In these topovectorial processes of Nrf1α, it is subjected to specific post-translational modifications (e.g. glycosylation, deglycosylation, ubiquitination), and also selective juxtamembrane proteolytic processing of the CNC-bZIP factor so as to yield multiple isoforms with different and opposing activities, during maturation into an activator (Koizumi et al., 2016; Xiang et al., 2018a; Xiang et al., 2018b). In addition, distinct variants of Nrf1, including its long TCF11, short Nrf1β/LCR-F1 and small dominant-negative Nrf1γ/δ, are also generated by alternative translation from various lengths of alternatively-spliced mRNA transcripts (Zhang et al., 2014a). Yet, each Nrf1 isoform-specific physiological function virtually remains obscure.

Notably, specific gene-editing knockout of Nrf1α leads to a significant increase in the malignant proliferation of *Nrf1α^−/−^-*derived hepatoma and the tumor metastasis to the liver in xenograft model mice (Ren et al., 2016). This work reveals that Nrf1α may act as a tumor suppressor, but the underlying mechanism remains unclear. Herein, our present work further unravels that Nrf1 and Nrf2 have mutual opposing and unified inter-regulatory cross-talks towards downstream genes. For instance, aberrant hyperactivation of Nrf2 leads to a constitutive increase of its target cycloxygenase-2 (COX2) in *Nrf1α^−/−^* cells. Such hyper-activation of Nrf2 by knockout of Nrf1α is accompanied by substantial decreases in both Keap1 and PTEN. The malignant growth of *Nrf1α^−/−^*-derived tumor is significantly prevented by knockdown of Nrf2, while *Nrf1α^+/+^*-bearing tumor is also markedly suppressed by knockout of Nrf2, but unaffected by *a priori* constitutive activator of Nrf2 (i.e. caNrf2^ΔN^). Such distinct phenotypes of animal xenograft tumors are determined by differential expression of different subsets of genes regulated by Nrf1α or Nrf2 alone or both. These collective findings demonstrated that Nrf1α manifests as a dominant tumor-suppressor to confine Nrf2 oncogenicity. Conversely, though Nrf2 acts as a tumor promoter, it directly mediates the transcriptional expression of *Nrf1* so as to form a negative feedback loop.

## RESULTS

### The human *Nrf1α^−/−^* -and *Nrf2^−/−ΔTA^*-driven cell models are established

Since the phenotypes of liver-specific *Nrf1^−/−^* mice resemble human pathogenesis of hepatic steatosis, NASH and HCC (Ohtsuji et al., 2008; Tsujita et al., 2014; Xu et al., 2005), this is thus inferred available for exploring the underlying mechanisms whereby NASH is transformed for the malignant progression towards hepatoma (Figure 1A). However, it is unknown whether human Nrf1α exerts similar effects to those obtained from the aforementioned mouse models. For this end, a similar pathological model was here recapitulated by genome-editing knockout of *Nrf1α* from human HepG2 cells, aiming to elucidate the mechanism by which a non-resolving NASH-based inflammation is exacerbated. To achieve the genomic locus-specific knockout of *Nrf1α*, we created a pair of TALEN-directed constructs to yield a specific deletion of Nrf1α-derived isoforms from the single *Nfe2l1* gene, but with shorter variants *Nrf1β* to *Nrf1δ* unaffected (Figures 1B and S1A). In the parallel experiments, another pair of CRISPR/cas9-mediated constructs were engineered to delete the Nrf2-specific codons (covering its 42-175 amino acids within essential Keap1-binding and most of its transactivation domains) from the *Nef2l2* gene to yield an inactive mutant *Nrf2^−/−ΔTA^* (Figures 1D, 1E and S1B). Consequently, two monoclonal hepatoma cell lines of *Nrf1α^−/−^* and *Nrf2^−/−ΔTA^* were, respectively, established and confirmed to be true by sequencing their genomic DNAs, and Western blotting with specific antibodies (Figure 1, B to E). Further real-time qPCRs with specific primers that recognized distinct nucleotide fragments showed that knockout of *Nrf1α* substantially abolished expression of total *Nrf1* mRNAs in *Nrf1α^−/−^* cells (Figure 1B, *right panel*). Similar results were also obtained from other clones of *Nrf1α^−/−^* cell lines (Ren et al., 2016). Notably, *Nrf2^−/−ΔTA^* cells gave rise to an inactive mutant lacking nt124-526 of Nrf2, but the expression of its DNA-binding domains (DBD)-containing mRNA transcripts was unaltered (Figure 1D, *right panel*). Therefore, the resulting inactive mutant *Nrf2^ΔTA^* polypeptides may still, theoretically, bind Nrf2-target genes, and also circumvent competitive occupancy with other complementary factors, upon loss of the prototypic Nrf2.

**Figure 1.**
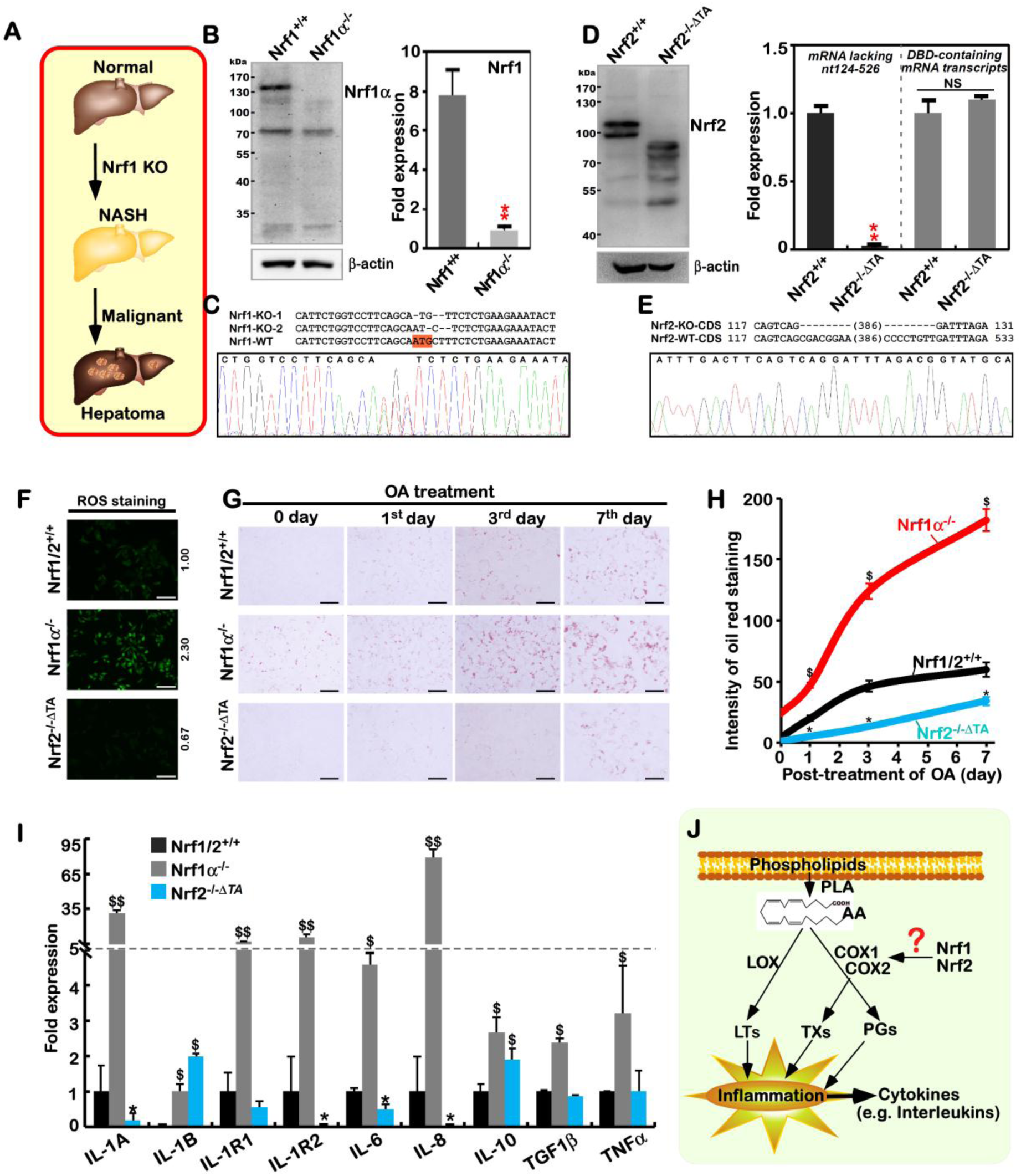
Establishment of Nrf1α-specific knockout cell models with the NASH phenotype. (A) Schematic diagrams for the liver-specific *Nrf1^−/−^* knockout mice that develop spontaneous non-alcoholic steatohepatitis (NASH) and deteriorate into hepatoma eventually. (B) Both Western blotting (WB, *left*) and real-time quantitative PCR (qPCR, *right*) were employed to identify the protein and mRNA levels of Nrf1 in a monoclonal *Nrf1α^−/−^* knockout cell line. The data are shown as mean ± SEM (n = 3×3, \**p* < 0.01). (C) Sequencing peaks of the genomic DNA fragments across the *Nrf1α-*specific knockout site, as indicated by alignment with wild type (WT) standard sequence. (D) Expression of inactive Nrf2 mutant protein and mRNA levels in a monoclonal *Nrf2^−/−^Δ^TA^* cell line was identified by WB (*left*) and qPCR with distinct primer pairs (*right*). The data are shown as mean ± SEM (n = 3×3, \**p* < 0.01; NS = no statistical difference). (E) Sequencing peaks of the genomic DNA fragments across the *Nrf2-*specific knockout site, as indicated by alignment with normal WT sequence. (F) ROS staining of *Nrf1/2^+/+^, Nrf1α^−/−^* and *Nrf2^−/−^Δ^TA^* cells. They had been treated with 5 μM of DCFH-DA (2’,7’-Dichlorodihydro-fluorescein diacetate) for 30 min, before being photographed under a fluorescence microscope. Scale bar = 100 μm. (G) Lipid staining of *Nrf1/2^+/+^, Nrf1α^−/−^* and *Nrf2^−/−^Δ^TA^* cells, that were or were not treated with 200 μM oleic acid (OA), before being stained with the oil red O agent, and photographed under a microscope. Scale bar = 25 μm. (H) Statistical analysis of the above lipid-stained (*G*) intensity, that was quantified and shown graphically. The data are represented as mean ± SEM (n = 3), with significant increases ($) or decreases (*), *p*< 0.01, compared with wild-type values. (I) The expression of inflammation-related genes in *Nrf1/2^+/+^, Nrf1α^−/−^* and *Nrf2^−/−^Δ^TA^* cells. The data obtained from transcriptome FPKM are shown as mean ± SEM (n=3; $, *p*< 0.01; $$, *p*< 0.001 and \**p*< 0.01, by comparison with wild-type). (J) Diagrammatic representation of a proposed model for Nrf1 and Nrf2 to regulate COX1 and COX2 essential for arachidonic acid metabolism relevant inflammatory response.

A plausible explanation of NASH pathogenesis is preferred to the classic two-hit hypothesis, in which the first hit is hepatosteatosis (caused by the accumulation of cholesterol and lipids), and the second hit is inflammation (induced by oxidative stress) (Friedman et al., 2018; Shimano and Sato, 2017). Such being the case, we next examine whether *Nrf1α^−/−^* cells act accordingly. As anticipated, it is illustrated by measuring intracellular hydrogen peroxide as a representative of reactive oxygen species (ROS), that endogenous oxidative stress was strikingly induced in *Nrf1α^−/−^* cells, but slightly relieved by inactive *Nrf2^−/−ΔTA^*, when compared with wild-type *Nrf1/2^+/+^* progenitor cells (Figure 1F). Subsequently, a significant accumulation of lipids was seen after staining of *Nrf1α^−/−^* cells, by comparison with *Nrf2^−/−ΔTA^* and *Nrf1/2^+/+^* cells (Figure 1G). With increasing time of oleic acid (OA) treatment to 7 days, the lipid accumulation was significantly incremented in *Nrf1α^−/−^* cells to a maximum ∼182-fold estimated. While compared with ∼60-fold accumulation of lipids in *Nrf1/2^+/+^* cells, the overload was substantially alleviated by inactive *Nrf2^−/−ΔTA^* to ∼34-fold (Figure 1, G & H).

In addition to lipid metabolic disorders resulting from loss of Nrf1’s function to regulate *LPIN1, PGC-1β* and other metabolic genes (Hirotsu et al., 2012; Tsujita et al., 2014), NASH has a not-yet-identified characteristic of refractory inflammation. Accordingly, we determined the transcriptional expression of key genes encoding cytokines and relevant receptors involved in putative inflammatory responses. As expected, expression of all nine examined genes encoding IL-1A, IL-1B, IL-1R1, IL-1R2, IL-6, IL-8, IL-10, TGF1α, and TGF1β was significantly elevated in *Nrf1α^−/−^* cells (Figure 1I). By contrast with *Nrf1/2^+/+^* cells, mRNA expression of IL-1A, IL-1R2, IL-6 and IL-8 was markedly down-regulated by inactive *Nrf2^−/−ΔTA^*, while both IL-1B and IL-10 expression was still marginally up-regulated, but with no changes in other genes (Figure 1I). Collectively, these data demonstrate that the NASH-prone phenotypes are recapitulated by employing human *Nrf1α^−/−^*-driven cells, in which Nrf2 may also be critical for this pathogenesis.

### The inflammation marker COX2 is up-regulated, while COX1 is down-regulated, in *Nrf1α^−/−^* cells

Development of inflammation (e.g. NASH) and malignant transformation into carcinogenesis have a clear relevance to lipid peroxidation, and particularly degradation metabolites of arachidonic acid (AA), such as prostaglandins (PGs), thromboxanes (TXs) and leukotrienes (LTs) (Castellone et al., 2005; Chowdhry et al., 2010; Gupta and Dubois, 2001). In the AA metabolism network, cyclooxygenase 1 (COX1) and COX2 are the rate-limiting enzymes that convert AA into PGs, of which COX2 is considered as a key inflammation marker (Wang and Dubois, 2010) and also identified as a direct target of Nrf2 (Itoh et al., 2004; Sherratt et al., 2003). Since Nrf2 (and Nrf1) can be recruited to directly bind the ARE-containing promoters of *COX2* and *COX1* before transactivating both genes, it is thereby hypothesized that hyper-expression of the inflammation-related genes in *Nrf1α^−/−^* cells is attributable to overstimulation of PG and TX products from the catalyzation by COX2 and COX1 (Figure 1J). To address this, we here examined whether (and how) key rate-limiting enzymes in AA metabolism are influenced by loss of *Nrf1α* or *Nrf2* functions.

As anticipated, a real-time qPCR analysis revealed that mRNA levels of *COX1* were almost completely abolished in *Nrf1α^−/−^* cells, but obviously increased in *Nrf2^−/−ΔTA^* cells by comparison to those obtained from wild-type *Nrf1/2^+/+^* cells (Figure 2A). Contrarily, expression of *COX2* was substantially augmented in *Nrf1α^−/−^* cells, but almost unaffected by inactive *Nrf2^−/−ΔTA^* when compared to the value in *Nrf1/2^+/+^* cells. Furthermore, expression of *ALOX5* and *FLAP* in *Nrf1α^−/−^* cells was up-regulated at much higher levels than those measured in *Nrf2^−/−ΔTA^* cells at considerable levels (Figure 2A). Next, whether such differences in expression of the AA metabolism genes are attributable to differential and even opposing regulation by Nrf1 and Nrf2 was further examined. Consistently, almost no protein expression of COX1 was detected in *Nrf1α^−/−^* cells, while the abundances of COX2 and ALOX5 proteins were significantly increased, when compared with *Nrf1/2^+/+^* cells (Figure 2B). However, COX1 was highly expressed in *Nrf2^−/−ΔTA^* cells at a greater level than that obtained from wild-type cells (Figure 2C). Conversely, COX2 was substantially diminished or abolished by the inactive mutant *Nrf2^−/−ΔTA^*, whereas ALOX5 was almost unaffected (Figure 2C).

**Figure 2.**
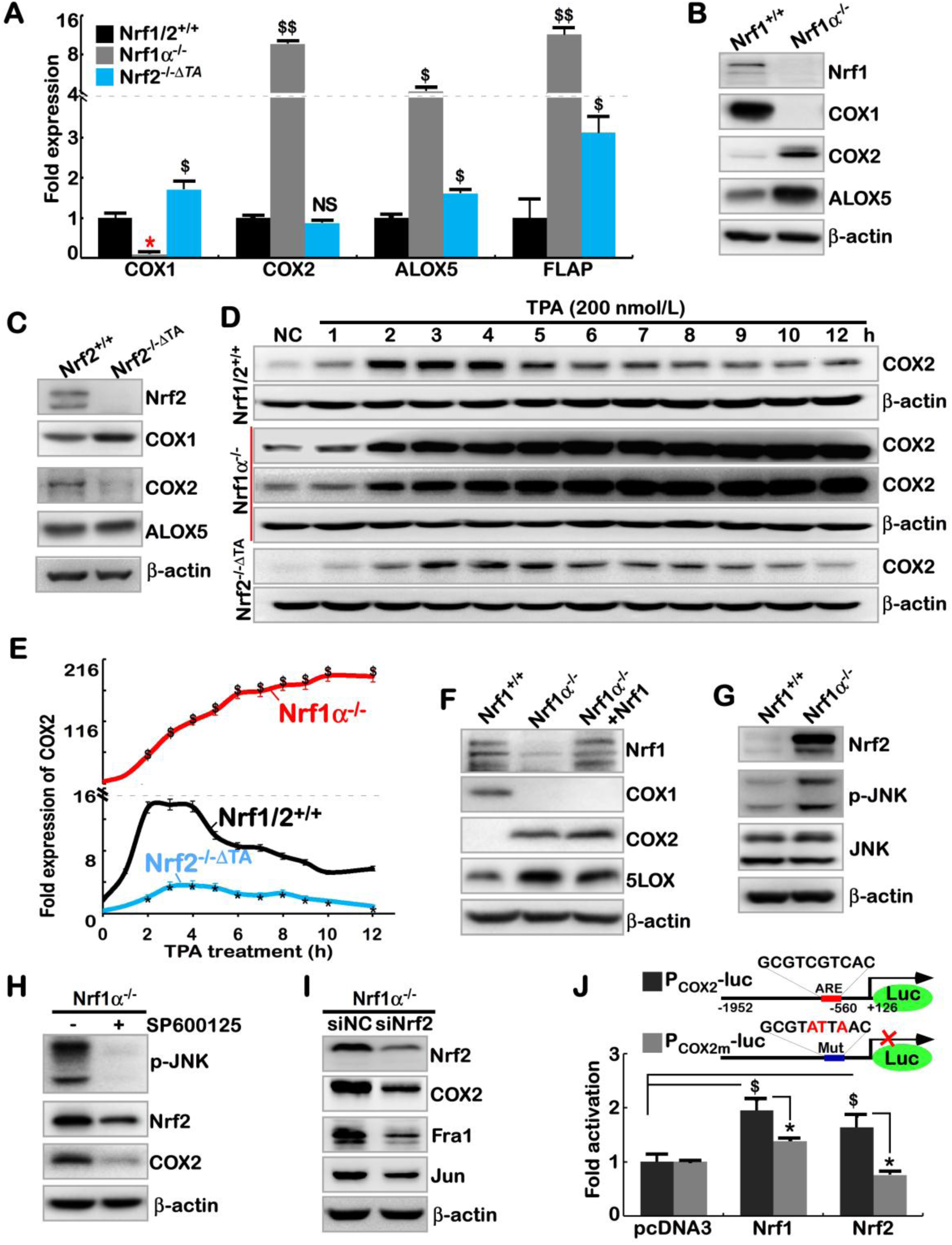
Differential or opposing roles of Nrf1α and Nrf2 in regulating *COX2* and *COX1* genes. (A) The mRNA levels of *COX1, COX2, ALOX5*, and *FLAP* was determined by real-time qPCR in *Nrf1/2^+/+^, Nrf1α^−/−^* and *Nrf2^−/−ΔTA^* cells. The data are shown as mean ± SEM (n = 3×3, \**p*< 0.01; $, *p*< 0.01; $$, *p*< 0.001). (B) The protein levels of COX1, COX2, ALOX5, Nrf1, and β-actin (as a loading control) in *Nrf1α^−/−^* and *Nrf1/2^+/+^* cells were visualized by Western blotting. (C) Western blotting of *Nrf1α^−/−^* and *Nrf2^−/−ΔTA^* cells to determine protein levels of COX1, COX2, ALOX5, Nrf1, and β-actin. (D) Time-course analysis of COX2 in *Nrf1/2^+/+^, Nrf1α^−/−^* and *Nrf2^−/−ΔTA^* cells, that had been treated for 1 h to 12 h with 100 nM TPA (12-*O*-tetradecanoyl phorbol-13-acetate), before being examined by Western blotting. (E) The intensity of the above anti-COX2 immunoblots (*D*) was quantified by normalizing the untreated value, which is shown graphically. The data are shown as mean ± SEM (n = 3, \**p*< 0.01; $, *p*< 0.01). (F) After the restoration of Nrf1 into *Nrf1α^−/−^* cells by packaging with the Lentivirus, changed abundances of Nrf1, COX1, COX2, and ALOX5 among *Nrf1/2^+/+^, Nrf1α^−/−^* and *Nrf1α^−/−^*+Nrf1-restored cell lines were examined by Western blotting. (G) Differences in Nrf2, p-JNK and JNK expression between *Nrf1/2^+/+^* and *Nrf1α^−/−^* cells were unraveled by Western blotting. (H) The changes in p-JNK, Nrf2 and COX2 were examined, following 24 h treatment of *Nrf1α^−/−^* cells with 20 μM of SP600125. (I) Alterations in Nrf2, COX2, Fra1, and Jun by siRNA interference with Nrf2 in *Nrf1α^−/−^* cells were determined by Western blotting. (J) The human *COX2* promoter-driven reporter *P_COX2_-luc* and its mutant (*upper*) were constructed before the luciferase assay. *Nrf1/2^+/+^* cells were co-transfected with either *P_COX2_-luc* or mutant, together with an internal control reporter *pRL-TK*, plus an expression construct for Nrf1 or Nrf2 or empty pcDNA3 plasmid and allowed for 24-h recovery before the *P_COX2_-luc* activity was calculated (*lower*). The data are shown as mean ± SEM (n = 3×3, * p< 0.01; $, p< 0.01 compared to the pcDNA3 values).

Since COX1 is constitutive essential for normal physiological homeostasis, while COX2 is an inducibly expressed enzyme to be stimulated by inflammatory stress (O’Banion, 1999; Tanabe and Tohnai, 2002), the changing trends of COX2 induction by 12-O-Tetradecanoylphorbol-13-acetate (TPA) stimulation are evaluated. As a result, stimulation of *Nrf1/2^+/+^* cells by TPA caused an obvious induction of COX2 expression to a ∼14-fold maximum at 2 h after treatment; this value was being maintained to 4 h, before being gradually decreased to a ∼5-fold level by 10-h treatment (Figure 2D, 2E). In *Nrf1α^−/−^* cells, the constitute up-expression of COX2 was set to 18-fold as its starting point, subsequent incremental abundances of this enzymatic protein were further induced to a maximum of ∼190-fold by 10-h TPA treatment and maintained before the experiment was terminated (Figure 2E,*red curve*). Relatively, a weak response of COX2 to TPA stimulation of *Nrf2^−/−ΔTA^* cells was also observed from 2 h to 8 h, with an inducible peak at 4 h after treatment (Figure 2E, *blue curve*). Further assays of the luciferase reporter *P_COX2_-Luc* (in which the 2078-bp promoter *COX2* gene was constructed) revealed that transcriptional expression of the reporter gene was induced at 4 h after TPA treatment of *Nrf1α^−/−^* cells, and the TPA-stimulated increases were continuously maintained until 24 h (Figure S2A, S2B). However, no obvious changes in the *P_COX2_-Luc* activity were detected in TPA-treated *Nrf2^−/−ΔTA^* or *Nrf1/2^+/+^* cells. These collective findings imply a striking disparity in Nrf1α-and Nrf2-mediated induction of COX2 by TPA.

### Hyper-expression of COX2 results from increased Nrf2 and JNK-mediated AP-1 in *Nrf1α^−/−^* cells

Intriguingly, the abundance of COX2, as a well-known direct target of Nrf2, was not decreased, but rather marginally increased by ectopic expression of Nrf1 that had been restored into *Nrf1α^−/−^* cells (Figure 2F), in which expression of COS-1 was also rescued, albeit both genes encompass the ARE sites recognized by Nrf1 and Nrf2 (Itoh et al., 2004; Sherratt et al., 2003). These seemingly paradoxical results, along with the above-described data from *Nrf1α^−/−^* cells, suggest that Nrf1α may have an ambivalent relationship with Nrf2 in regulating *COX1* and *COX2* genes. Rather, this confusing but exciting finding arouses our *de facto* curiosity to explore which possible pathways enable Nrf1 to indirectly regulate COX2 (Figure S3A), albeit this enzyme has been shown to be monitored by CREB, NF-κB, STAT1, FOXM1, ETS1, ELF3 and JNK-regulated AP1 (Ghosh et al., 2007; Grall et al., 2005; Kang et al., 2006; Sharma-Walia et al., 2010; Xu and Shu, 2013; Zhang et al., 2007a). Consequently, the real-time qPCR analysis revealed that mRNA levels of only *RELB,* but not other members of the NF-κB family (that regulates cellular responses to inflammation), were significantly up-regulated in *Nrf1α^−/−^* cells (Figure S3B). This may be coincident with the notion that ablation of an IκB (inhibitor of NF-κB) kinase IKKγ in liver parenchymal cells causes spontaneous NASH and HCC (Luedde et al., 2007). However, Figure S3C showed that the abundance of over-expressed COS2 in *Nrf1α^−/−^* cells was unaltered by the caffeic acid phenethyl ester (CAPE, a potent specific inhibitor of NF-κB (Natarajan et al., 1996)), and was also not significantly diminished by JSH-23, a broad spectrum inhibitor of NF-κB (Shin et al., 2004). Thus, it is inferable there implicates an NF-κB-independent pathway to up-regulate expression of COX2 in *Nrf1α^−/−^* cells. In addition, it may be not necessary for modest inducible expression of *ETS1* (one of the E26 transformation-specific transcription factors), because this was accompanied by substantial down-regulation of another family member *ELF*3 (Figure S3B).

Further treatments of *Nrf1α^−/−^* cells with either of the two CREB inhibitors H-89 and BAPTA-AM (Sharma-Walia et al., 2010; Tymianski et al., 1994) demonstrated that the elevated expression of COX2 was unaffected (Figure S3D). However, it is, to our surprise (Figure S3E), found that the forced abundance of COX2 in *Nrf1α^−/−^* cells was sufficiently abolished by a JNK-specific inhibitor SP600125 (Bennett et al., 2001). Further investigations revealed no changes in both mRNA and protein levels of JNK (Figures 2G and S3), but the phosphorylated JNK abundance was significantly increased in *Nrf1α^−/−^* cells when compared to those obtained from *Nrf1/2^+/+^* cells (Figure 2G). Therefore, it is initially postulated that *Nrf1α^−/−^* cells gave rise to the forced expression of COX2 possibly by activation of JNK signaling. Next, in-depth insights into the signaling of JNK towards downstream target genes unraveled that expression of only *c-Jun*, but not other examined genes, was significantly elevated in *Nrf1α^−/−^* cells (Figures S4B). Further assays of luciferase reporter genes *P_COX2_-Luc* and *P_TRE_-Luc* (in which TRE indicates TPA-responsive element inserted within the promoter region) verified that AP-1 (a functional heterodimer of Jun and Fos) is favorably required for the transactivation of COX2 in *Nrf1α^−/−^* cells (Figure S4C). By defining distinct AP-1 components (e.g. Jun, Fos, Fra1) at mRNA and protein levels, it is validated that AP-1 was activated in *Nrf1α^−/−^* cells, but not in *Nrf2^−/−ΔTA^* cells (Figure S4, D to F). In addition, Figure S4G illustrated that hyper-expression of COX2 in *Nrf1α^−/−^* cells was not suppressed by AP-1 inhibitor SR11302 (Fanjul et al., 1994). Further knockdown of Jun or Fra1 led to a decrease of COX2, but this was no proportional to the silencing of Jun or Fra1 at lower levels (Figure S4H). All together, AP-1 activation by JNK signaling is involved in, but not essential for making a significant contribution to the reinforced expression of COX2 in *Nrf1α^−/−^* cells.

Fortunately, the evidence that expression of Nrf2 and its nuclear translocation are attenuated by JNK inhibitor SP600125 (Ahn et al., 2017; Bak et al., 2016) implicates there exists a direct link between JNK and Nrf2. Consistently, the abundance of Nrf2 protein was surprisingly augmented in *Nrf1α^−/−^* cells, which was accompanied by an increase in the phosphorylated JNK (Figure 2G). Similarly, great increases in expression of both COX2 and Nrf2 were caused by knockout of Nrf1α in HL7702^Nrf1α*−/−*^ (established on the base of the non-cancerous HL7702 hepatocyte line (Figure S5A,B). Further treatments of *Nrf1α^−/−^* cells with SP600125 or transfection with Nrf2-targeting siRNA, unraveled that reduction of Nrf2 or JNK appeared to be proportional to the decreased abundance of COX2 (Figure 2H, 2I and S4I). Collectively, these data indicate that the hyper-expression of COX2 in *Nrf1α^−/−^* cells is directly caused by increased Nrf2 protein, and the latter CNC-bZIP is also monitored by its upstream JNK signaling. This is supported by the *P_COX2_- Luc* reporter assays showing that Nrf2 mediated transactivation of the *COX2* gene driven by its ARE enhancer, because the activation was attenuated by its ARE mutant (i.e. P_COX2m_-*-Luc*) (Figure 2J). The transactivation of *P_COX2_-Luc* reporter by Nrf1 (Figure 2J), together with above-described data, indicates that Nrf1α also has one hand to exert a minor positive effect on COX2 by directly binding to its ARE enhancer, but this effect appears to be counteracted by another hand of Nrf1α to elicit a dominant-negative role by indirect inhibitory pathways.

Some AP-1 abundances were obviously suppressed by silencing Nrf2 in *Nrf1α^−/−^* cells (Figure 2I), and strikingly prevented by inactive *Nrf2^−/−ΔTA^*, by comparison with equivalent controls (Figure S4F). Together with the above-described results, these imply that AP-1 is dominantly repressed by Nrf1α, but positively regulated by Nrf2. Rather, no available evidence has been presented here to support the notion that AP-1 activates transcription of Nrf2 (Tao et al., 2014). In mouse embryonic fibroblasts (MEFs), COX2 is co-regulated by Nrf1 and Nrf2, because its abundance was significantly abolished by global knockout of Nrf1 or Nrf2 (Figure S5C). Here, it should also be noted that global knockout of all mouse Nrf1 or Nrf2 DNA-binding domain-containing fragments was achieved by both gene-targeting manipulations (Ohtsuji et al., 2008; Tsujita et al., 2014). This is totally distinctive from specific gene-editing to delete the designed portions of human Nrf1α or Nrf2 (Figure 1). Importantly, knockout of Keap1 in MEFs (Figure S5D) and human HepG2 (Figure S5E) caused a remarkable increase in expression of Nrf1, Nrf2, COX2, and HO-1 to different extents detected. Overall, the precision regulation of COX2 by Nrf1 and Nrf2, along with Keap1, in distinct manners, is preferable to depend on distinctive cell types in different species.

### Nrf1α and Nrf2 transactivate ARE-driven miR-22 signaling to PTEN, but not to COX1

On the contrary to COX2, the isoenzyme COX1 was highly expressed in *Nrf2^−/−ΔTA^* cells (Figure 2C), but its expression was almost completely abolished in *Nrf1α^−/−^* cells (Figure 2B) and also not rescued by restoration of ectopic Nrf1 into *Nrf1α^−/−^* cells (Figure 2F), albeit Nrf2 was up-regulated (Figure 2G). Thereby, it is inferable that no matter how Nrf1α and Nrf2 have opposing or overlapping roles in regulating COX1 expression, Nrf2 exerts a dominant inhibitory effect on COX1, but this effect is fully contrary to regulating COX2. Thus, we speculate that the putative inhibition of COX1 by Nrf2 (and possibly Nrf1α) may be achieved through an indirect miRNA-regulatory pathway, except for directly ARE-binding to this gene. Fortunately, a candidate miR-22 was selected by predicting possibly miRNA-binding sites within the *COX1* 3′-UTR region (see http://www.targetscan.org/vert_72/). As expected, the real-time qPCR analysis unraveled that miR-22 expression was significantly increased in *Nrf1α^−/−^* cells but decreased in *Nrf2^−/−ΔTA^* cells (Figure 3A). Forced expression of Nrf1 or Nrf2 also caused an obvious increase in miR-22 expression in wild-type *Nrf1/2^+/+^* cells (Figure 3B). Further analysis of the *miR-22*-coding gene revealed there exists a consensus ARE site within its promoter (Figure 3C, *upper panel*). The promoter-driven luciferase reporter (i.e. miR22-ARE-Luc) was created, so as to assay for its transcription activity. The results showed that the *miR22-ARE-Luc* reporter gene was significantly transactivated by Nrf1 and Nrf2 (Figure 3C), and the transactivation was diminished by the mutant *miR22-AREm-luc*. Together, these imply direct and indirect transactivation of miR-22 possibly by Nrf1α and Nrf2.

**Figure 3.**
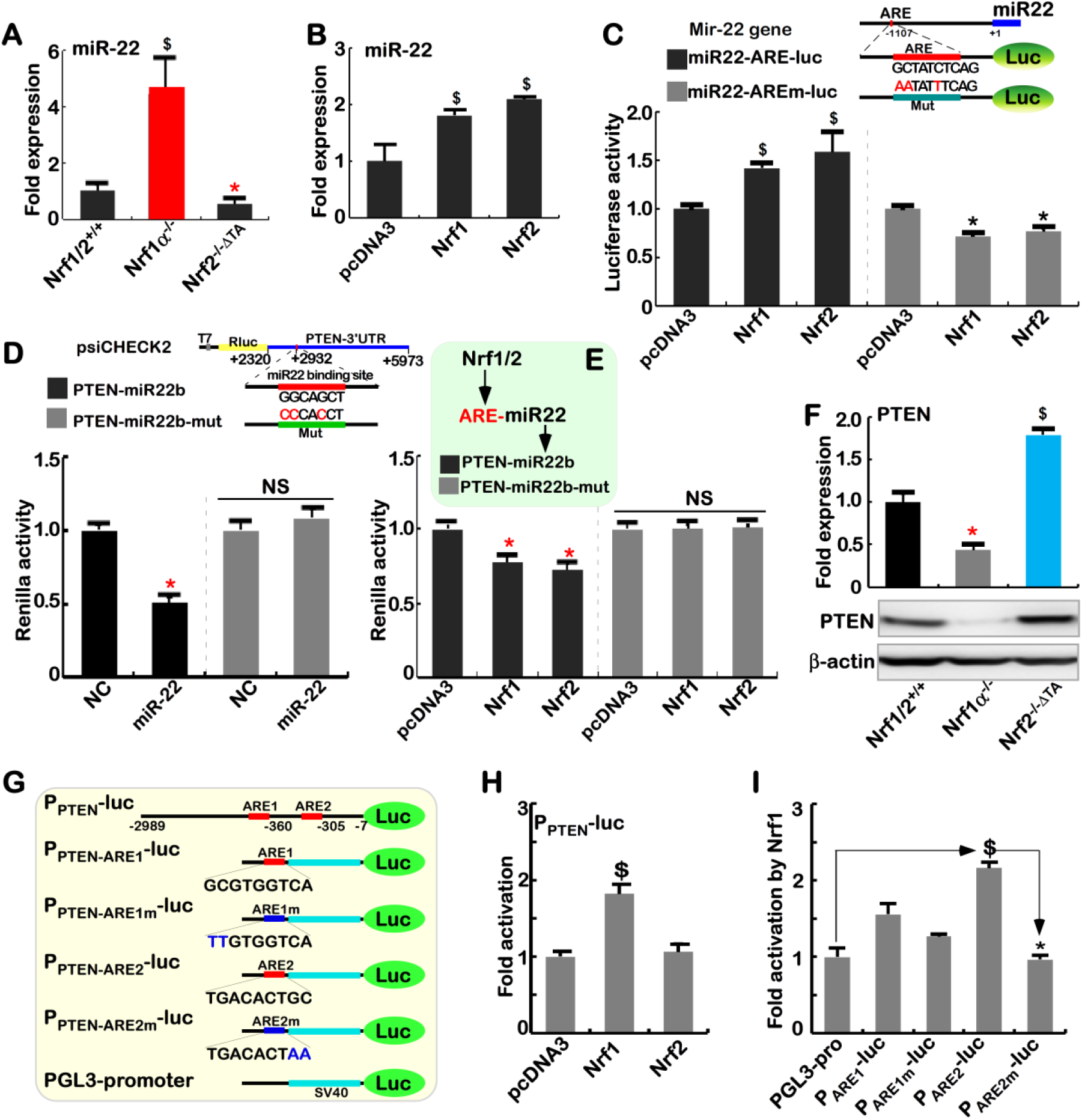
Different regulation of PTEN by Nrf1α and Nrf2 occurs through miR-22. (A) The content of miR-22 in *Nrf1/2^+/+^, Nrf1α^−/−^* and *Nrf2^−/−ΔTA^* cells was determined by qPCR with miR-22 specific primers. The data are shown as mean ± SEM (n = 3×3; \**p*< 0.01; $, *p*< 0.01 compared to wild-type values). (B) The miR-22 expression was altered by transfection of an expression construct for Nrf1 or Nrf2, or an empty pcDNA3 control, into *Nrf1/2^+/+^* cells. The qPCR data are shown as mean ± SEM (n = 3×3; $, *p*< 0.01). (C) The *miR22-ARE-luc* reporter driven by an ARE enhancer in the mir-22 gene promoter, and its mutant reporter *miR22- AREm-luc* were constructed. Either of reporter genes, together with *pRL-TK,* plus each of pcDNA3, Nrf1, or Nrf2 expression constructs were co-transfected into *Nrf1/2^+/+^* cells and then allowed for 24-h recovery before the luciferase activity measured. The data are shown as mean ± SEM (n = 3×3; \**p*< 0.01; $, *p*< 0.01) (D) There exists a miR-22 binding site in the PTEN’s 3’UTR region (that was constructed in the dual fluorescent psiCHECK2 vector to yield the *PTEN-miR22b* reporter). Either of *PTEN-miR22b* and *PTEN-miR22b-mut* was co-transfected with miR-22 expression plasmid or a negative control (NC) into *Nrf1/2^+/+^* cells, and then allowed for 24-h recovery, before the fluorescent activity was determined. The data are shown as mean ± SEM (n = 3×3; \**p*< 0.01; NS= no statistical difference). (E) Either *PTEN-miR22b* or *PTEN-miR22b-mut* was co-transfected with each of pcDNA3, Nrf1, or Nrf2 expression constructs *Nrf1/2^+/+^* cells and allowed for 24-h recovery, before the fluorescent activity was measured. The data are shown as mean ± SEM (n = 3×3; \**p*< 0.01). (F) The mRNA (*upper*) and protein (*lower*) levels of PTEN in *Nrf1/2^+/+^, Nrf1α^−/−^* and *Nrf2^−/−ΔTA^* cells were determined by qPCR and Western blotting, respectively. The data are shown as mean ± SEM (n = 3×3; \**p*< 0.01; $, *p*< 0.01). (G) Schematic representation of the PTEN promoter-containing *P_PTEN_-luc*, its distinct ARE-driven reporters (*P_ARE1_-luc* and *P_ARE2_-luc*) and indicated mutants, which were constructed into the PGL3-Promoter (*PGL3-Pro*) vector. (H) The *P_PTEN_-luc* and *pRL-TK*, along with an expression construct for Nrf1 or Nrf2, or pcDNA3 were co-transfected into *Nrf1/2^+/+^* cells and then allowed for 24-h recovery before being measured. The luciferase activity data are shown as mean ± SEM (n = 3×3; $, p< 0.01). (I) *Nrf1/2^+/+^* cells were co-transfected with an indicated luciferase reporter, together with *pRL-TK* and Nrf1 expression construct or pcDNA3 for 24 h before being determined. The data are shown as mean ± SEM (n = 3×3; $, p< 0.01; *p< 0.01).

Since negative regulation of PTEN by miR-22 had been reported (Fan et al., 2016; Tan et al., 2012), a Renilla reporter gene containing the 3’UTR region of *PTEN* (i.e. *PTEN-miR22b*) was here constructed, together with a mutant of miR-22-binding site (i.e. *PTEN-miR22b-mut*, Figure 3D, *upper panel*). If miR-22 would bind the 3’-UTR region of *PTEN* transcripts, the *PTEN-miR22b*-driven Renilla activity was significantly reduced by miR22 (Figure 3D, *lower panel*), and also partially decreased by ectopic expression of Nrf1 or Nrf2 (Figure 3E). These negative effects were sufficiently abrogated by *PTEN-miR22b-mut*. Consistently, both mRNA and protein levels of PTEN were significantly reduced in *Nrf1α^−/−^* cells (with hyper-expression of Nrf2) but strikingly increased in *Nrf2^−/−ΔTA^* cells (Figure 3F). Notably, such opposing changes in PTEN levels are inversely correlated with the relevant values of miR22 measured in same cell lines (Figure 3A). Thus, transactivation of miR-22 by Nrf1α and Nrf2 leads to putative inhibition of PTEN.

To further determine whether such miR-22 pathway is involved in the regulation of COX1 by Nrf1 and Nrf2, the luciferase reporter gene was constructed by cloning the 3’-UTR sequence of *COX1* (i.e. *COX1-miR22b*), along with a mutant of the putative miR-22 binding site to yield a *COX1-miR22b*-mut reporter (Figure 6A, *upper panel*). However, the *COX1-miR22b*-driven Renilla activity was unaltered by miR-22, Nrf1 and Nrf2, when compared with that of *COX1- miR22b*-mut (Figure S6A1, S6A2). Another luciferase reporter gene (i.e. *P_COX1_-Luc*) was engineered by inserting the 1413-bp *COX1* gen*e* promoter, but the *P_COX1_-Luc* activity was also unaltered by forced expression of Nrf1 and Nrf2 (Figure 6B1). However, the responsiveness of the *P_COX1_-Luc* to TPA was induced (Figure S6B2), albeit it was relatively weak, when compared to the *P_COX2_-Luc* (Figure S2A). Intriguingly, the *P_COX1_-Luc* activity was modestly mediated by Jun, but almost unaffected by a canonical AP-1 dimer (Figure S6B3). This is consistent with the notion from (Smith et al., 1997), but further insights are required into the mechanisms underlying the regulation of COX1 by Nrf1α and Nrf2.

### Nrf1α and Nrf2 have mutual regulatory effects on downstream genes

Since an unusual increase in Nrf2 protein is accompanied by relative higher levels of ROS in *Nrf1α^−/−^* cells (Figures 1F, 2G), it is inferable that Nrf1α-deficient hepatoma cells are growing under severe redox stress conditions redefined at a new higher steady-state level leading Nrf2 to become hyperactive. As anticipated, mRNA expression levels of *HO-1, GCLC, GCLM, NQO1* and *xCT* (they are co-target genes mediated by Nrf1 and Nrf2 (Leung et al., 2003; Ohtsuji et al., 2008; Tsujita et al., 2014) were significantly increased *Nrf1α^−/−^* cells (Figure 4A). Meanwhile, a marked decrease in *LPIN1,* but no significant reduction in *PGC-1β* (both were identified as Nrf1-specific target genes by (Hirotsu et al., 2012)), was determined by comparison with equivalents of wild-type *Nrf1/2^+/+^* cells. Despite no obvious alterations in mRNA levels of Nrf2 (Figure 4A), Western blotting revealed significant increases in the abundance of Nrf2 protein and typical downstream gene products HO-1, GCLM, NQO1 and HIF1α in *Nrf1α^−/−^* cells, by contrast with *Nrf1/2^+/+^* cells (Figure 4B). All four protein levels of HO-1, GCLM, NQO1 and HIF1α were, however, markedly reduced in accordance with knockdown of Nrf2, by siRNA-targeting interference with *Nrf1α^−/−^* cells (Figure 3C). Silencing of Nrf2 also led to decreased mRNA expression levels of *HO-1, GCLM* and *xCT* (Figure S5F). Conversely, restoration of ectopic Nrf1 expression into *Nrf1α^−/−^* cells caused obvious decreases in abundances of Nrf2, HO-1, GCLM and NQO1 to different extents as detected (Figure 4D). Collectively, it is demonstrated that in *Nrf1α^−/−^* cells, hyper-active Nrf2 has a potent ability to mediate a subset of their co-target genes. Furthermore, the phosphorylated JNK, but not its total, proteins were markedly decreased, as Nrf2 protein was reduced by ectopic expression of Nrf1 after transfecting into *Nrf1α^−/−^* cells (Figure 4D). This finding, together with the evidence that Nrf2 is repressed by JNK inhibitor treatment of *Nrf1α^−/−^* cells (Figure 2H), implicates that Nrf2 may govern a not-yet-identified upstream kinase to phosphorylate JNK through a positive feedback loop.

**Figure 4.**
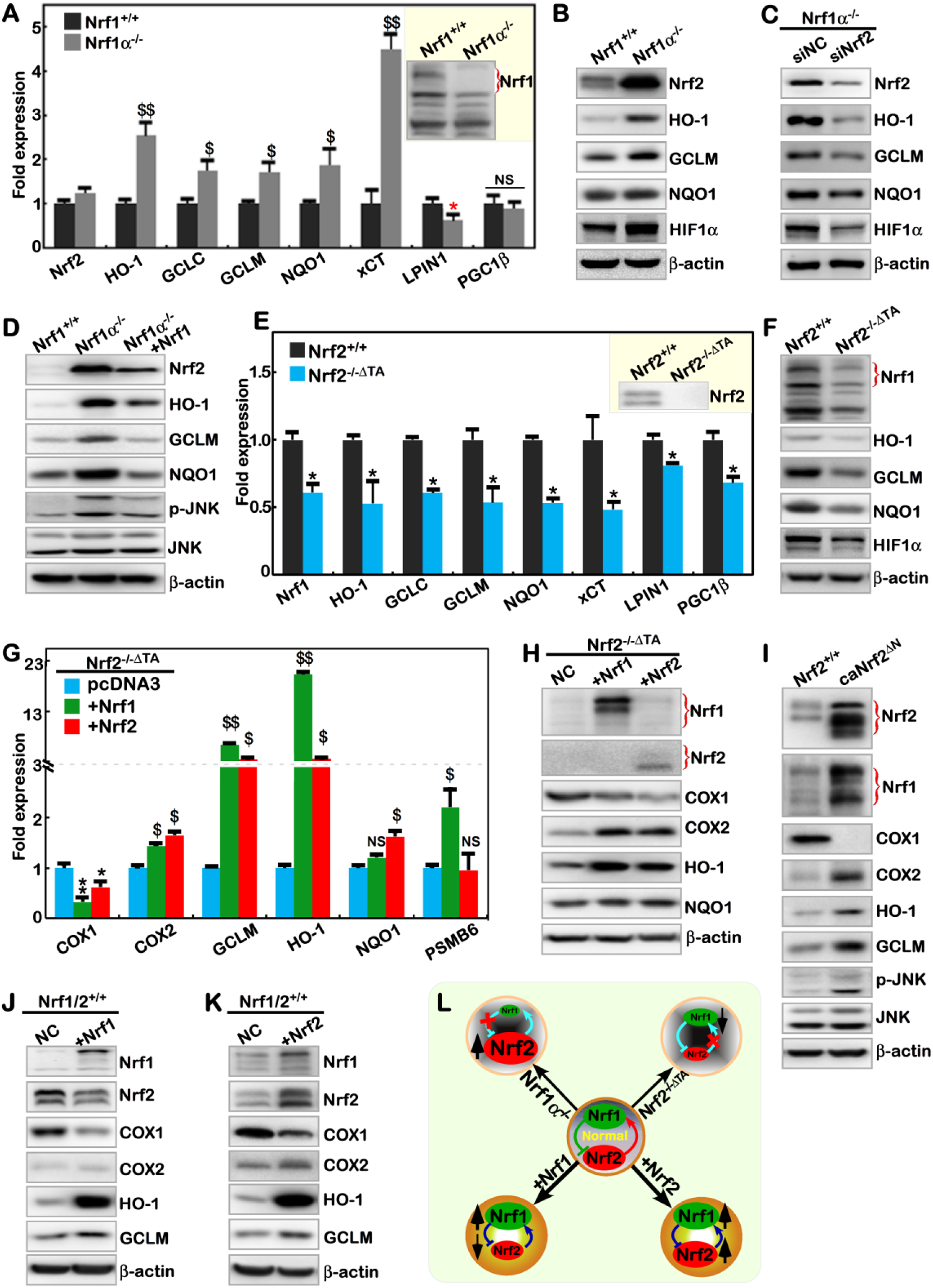
Opposing and unified cross-talks between Nrf1α and Nrf2. (A) Real-time qPCR determined the mRNA levels of *Nrf2, HO-1, GCLC, GCLM, NQO1, LPIN1,* and *PGC1β* expressed in *Nrf1/2^+/+^* and *Nrf1α^−/−^* cells. The data are shown as mean ± SEM (n = 3×3, \**p*<0.01; $, *p*<0.01; $$, *p*< 0.001. NS= no statistical difference). (B) The protein levels of Nrf1, HO-1, GCLM, NQO1 and HIF1α in *Nrf1/2^+/+^* and *Nrf1α^−/−^* cells were visualized by Western blotting. (C) *Nrf1α^−/−^* cells were interfered by siNrf2 (at 60 nM) to knock down Nrf2, and then allowed for 24-h recovery for 24 h, before abundances of HO-1, GCLM, NQO1 and HIF1a were examined by Western blotting. (D) After Nrf1 was allowed for restoration into *Nrf1α^−/−^* cells, changed protein levels of *Nrf2, HO-1, GCLM, NQO1, p-JNK and JNK* were determined in *Nrf1/2^+/+^, Nrf1α^−/−^* cells and *Nrf1α^−/−^* +Nrf1-restored cells (E) Expression of *Nrf1, HO-1, GCLC, GCLM, NQO1, LPIN1* and *PGC1β* genes in *Nrf1/2^+/+^* and *Nrf2^−/−ΔTA^* cells were analyzed by real-time qPCR. The data are shown as mean ± SEM (n = 3×3, \**p*< 0.01). (F) The protein levels of Nrf1, HO-1, GCLM, NQO1 and HIF1α in *Nrf1/2^+/+^* and *Nrf2^−/−ΔTA^* cells were seen by Western blotting. (G) *Nrf2^−/−ΔTA^* cells, that had been transfected with an expression construct for Nrf1 or Nrf2 or pcDNA3, were subjected to real-time qPCR analysis of *COX1, COX2, GCLM, HO-1, NQO1* and *PSMB6* expression. The data are shown as mean ± SEM (n = 3×3, \**p*< 0.01, \*\**p*< 0.001; $, *p*< 0.01; $$, *p*< 0.001. NS= no statistical difference). (H) Western blotting unraveled the changed abundances of Nrf1, Nrf2, COX1, COX2, GCLM, HO-1 and NQO1 proteins in *Nrf2^−/−^* cells as transfected with an expression construct for Nrf1 or Nrf2. NC = a negative control transfected with empty pcDNA3. (I) Alterations in protein levels of Nrf2, Nrf1, COX1, COX2, HO-1, GCLM, p-JNK and JNK in *Nrf1/2^+/+^* and *caNrf2^ΔN^*(containing the constitutive active Nrf2) cells were determined by Western blotting.

(J, K) *Nrf1/2^+/+^* cells were transfected with an expression construct for Nrf1 or Nrf2 or pcDNA3 and then allowed for 24-h recovery, before being examined by Western blotting to determine the changes in abundances of Nrf1, Nrf2, COX1, COX2, HO-1 and GCLM.

(L) A model is proposed to explain there exist an opposing and unifying inter-regulatory cross-talks between Nrf1 and Nrf2.

By contrast, inactivation of Nrf2 led to strikingly decreases in both mRNA and protein levels of Nrf1 in *Nrf2^−/−ΔTA^* cells (Figure 4E, 4F). This was accompanied by significantly diminishments in the expression of their co-regulated downstream genes *HO-1, GCLM, NQO1* and *HIF1α* in *Nrf2^−/−ΔTA^* cells (Figure. 4E, 4F), except with a modest reduction in *LPIN1* and *PGC-1β* mRNA levels. Thereby, such marked decreases in expression of *Nrf1, HO-1, GCLM, NQO1* and *HIF1α* resulting from loss of Nrf2 function demonstrate that *Nrf2^−/−ΔTA^* cell line could provide a favorite model to determine the changing downstream genes regulated by Nrf1, Nrf2 alone or both. Next, to address this, *Nrf2^−/−ΔTA^* cells were allowed for the ectopic expression of Nrf1 or Nrf2 in order to estimate specific downstream genes. As expected, it is validated that Nrf1-specific target gene *PSMB6* was increased by forced expression of Nrf1, but not Nrf2 allowed for restoration in *Nrf2^−/−ΔTA^* cells (Figure 4G). Conversely, expression of *NQO1* was induced by ectopic Nrf2, rather than Nrf1, after being transfected into *Nrf2^−/−ΔTA^* cells. This implies that *NQO1* is Nrf2-dependent, but insensitive to Nrf1 in *Nrf2^−/−ΔTA^* cells. In fact, Nrf1 and Nrf2 have overlapping roles in mediating transactivation of *HO-1* and *GCLM* (Figure 4G, 4H). Intriguingly, both CNC-ZIP factors also enhanced expression of *COX2*, but reduced *COX1* expression (Figure 4G, 4H). This seems consistent with additional examinations, revealing that silencing of Nrf2 in *Nrf1α^−/−^* cells gave rise to a relative higher expression of *COX1*, with an accompanying decrease in *COX2* (Figure S5F). Such co-inhibition of COX1 by two transcriptional activators Nrf1 and Nrf2 is puzzling, albeit it is known that transcriptional expression of downstream genes is mediated by their functional heterodimers with a partner of small MAF or other bZIP proteins through directly binding the *cis*-regulatory ARE sites in their promoters (Yamamoto et al., 2018; Zhang and Xiang, 2016). Together with the above data (Figures 2, 3), these collective findings indicate that Nrf1 and Nrf2 may also be two indirect players in the transcriptional regulation of COX1 by an unidentified pathway.

To determine which specific genes are constitutively activated by Nrf2, a dominant constitutive active mutant caNrf2^ΔN^-expressing cell line was established by the gene-editing to delete the N-terminal Keap1-binding portion of Nrf1 (Figure S5A). The resulting *caNrf2^ΔN^* cells indeed gave rise to a higher expression of Nrf2, as well as Nrf1, when compared to wild-type cells (Figures 4I, S5G). Interestingly, expression of COX1 almost disappeared as accompanied by significant increases of COX2 in *caNrf2^ΔN^* cells (Figures 4I, S5G). This further supports the above evidence obtained from inactivation of Nrf1α and Nrf2. Constitutive presence of caNrf2^ΔN^ also led to increases in abundances of both HO-1 and GCLM (Figure 4I), in addition to an enhanced expression of *xCT* and *Lpin1* (Figure S5G). Furthermore, phosphorylated JNK was significantly induced by caNrf2^ΔN^, with no changes in total JNK protein (Figure 4I), implying there may exist a putative upstream kinase monitored by Nrf2.

To further assess a mutual regulatory relationship between Nrf1 and Nrf2, we herein examined whether one of the endogenous proteins was influenced by another of both proteins that were allowed for ectopic over-expression in wild-type *Nrf1/2^+/+^* cells. As shown in Figure 4J, the endogenous Nrf2 protein was obviously decreased by ectopic Nrf1. Consequently, abundances of HO-1 and GCLM were markedly increased, whereas COX2 was weakly enhanced, but COX1 was significantly decreased following over-expression of Nrf1 (Figure 4J). By contrast, over-expression of ectopic Nrf2 caused an enhancement in endogenous Nrf1 (Figure 4K). This was accompanied by striking increases of COX2, HO-1 and GCLM, along with a remarkable decrease of COX1 (Figure 4K). Altogether, we assume there exists a mutual inter-regulatory relationship between Nrf1α and Nrf2, as summarized in Figure 4L. This may be an important strategy for a precision regulation of distinct downstream genes, to meet the needs for different cell processes.

### Nrf1α and Nrf2 transactivate the *Nrf1* promoter-driven reporter at different sites

To gain insights into the direct relationship between Nrf1 and Nrf2, we constructed their specific luciferase reporters by cloning the promoter regions of *Nrf1* and *Nrf2* genes and evaluated their activity by transfection into HepG2 cells (Figure 5A, 5B). As anticipated, the results showed that both *P_Nrf1_-luc* and *P_Nrf2_-luc* reporter genes were significantly induced by thapsigargin (TG, a classic ER stressor), or *tert*-Butylhydroquinone (tBHQ, a typical oxidative inducer), but not vitamin C (VC, a dual redox inducer) (Figure 5C). Thereby, *P_Nrf1_-luc* and *P_Nrf2_-luc* reporters are available to assess transcriptional expression of *Nrf1* and *Nrf2*. Subsequently, co-transfection of expression constructs for Nrf1 or Nrf2, together with *P_Nrf1_-luc* or *P_Nrf2_-luc* reporters, revealed that the transcription of *P_Nrf1_-luc*, but not *P_Nrf2_-luc*, genes was markedly induced by Nrf1 and Nrf2 (Figure 5D). Although no canonical ARE sequence (5′-TGACxxxGC-3′) exist in the 5025-bp *Nrf1* gene promoter, an attempt to identify which sites are located in the promoter enabling for specific transactivation by Nrf1 and Nrf2 was made, in order to yield a series of truncated mutants from *P_Nrf1_-luc* (Figures 5A). Fortunately, the resulting luciferase assays uncovered that several reporters containing the first exon region of *Nrf1* were activated by Nrf1 and Nrf2 possibly through different regulatory sites (Figure 5B). From various lengths of the *P_Nrf1_-luc* and mutants, it is inferable that the *Nrf1/Nfel1*-regulatory locus site-1 (i.e. Site-1) specific for Nrf2 is located in a 62-bp range between +572 bp and +634 bp, and the *Nrf1/Nfel1*-regulatory locus site-2 (i.e. Site-2) specific for Nrf1 *per se* is situated in another 100-bp range from +1031bp to +1131bp (Figure S7A).

**Figure 5.**
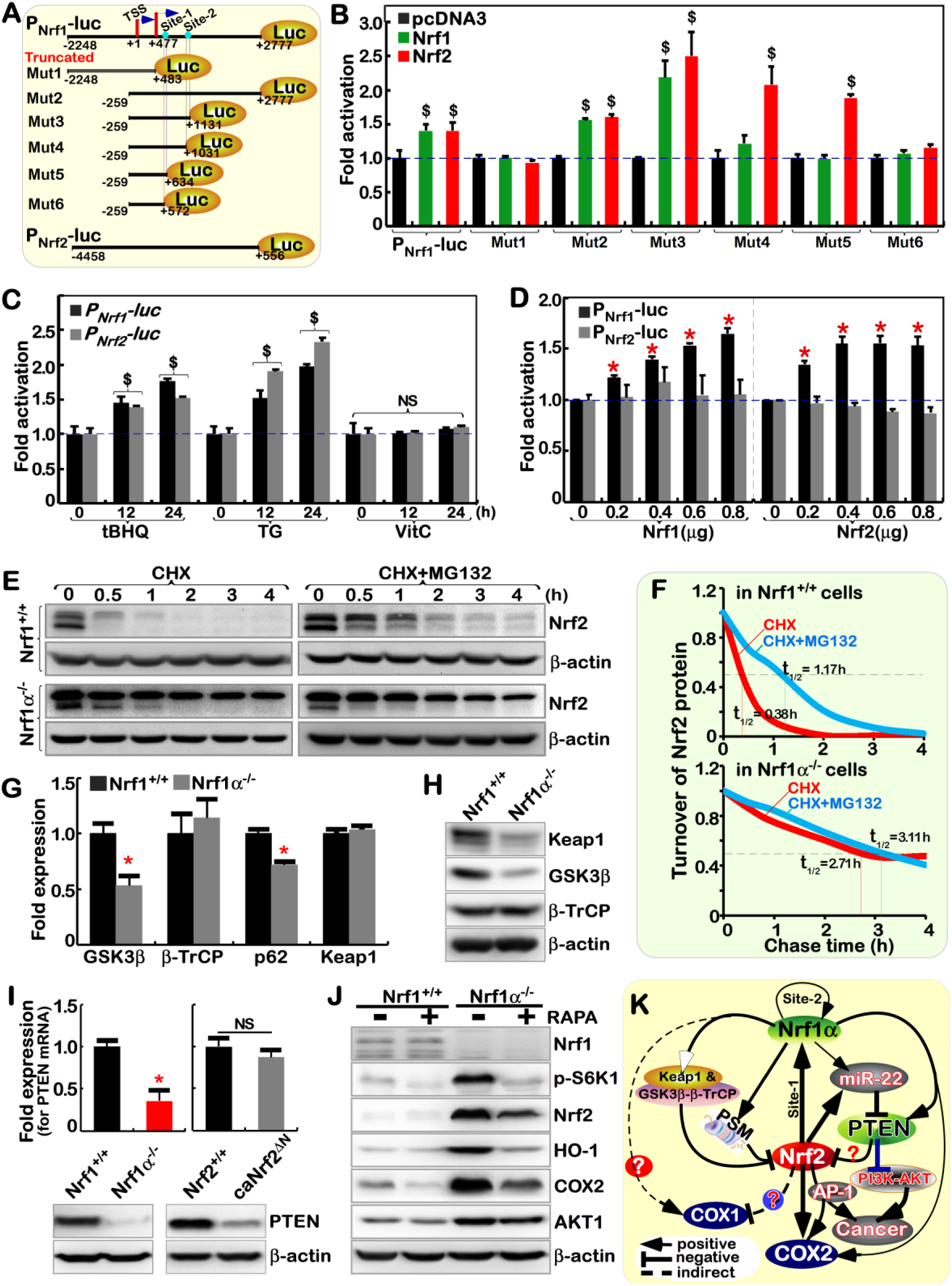
Nrf1α and Nrf2 have inter-regulatory cross-talks at distinct levels. (A) Schematic representation of both *P_Nrf1_-luc* and *P_Nrf2_-luc* reporters (driven by the human *Nrf1* and *Nrf2* gene promoters), along with various lengths of truncated *P_Nrf1_-luc* mutants as indicated, which were constructed in the PGL3-Basic vector. There exist two transcriptional starting sites (i.e. TSS1, TSS2) within the *Nrf1* gene promoter, which contains two *Nrf1/Nef2l1*-regulatory locus sites (i.e. Site-1, Site-2, also see Figure S6A). (B) Each of the *P_Nrf1_-luc* and indicated mutants, together with *pRL-TK*, plus an expression construct for Nrf1 or Nrf2 or pcDNA3, were co-transfected into *Nrf1/2^+/+^* cells and allowed for 24-h recovery, before the luciferase activity was measured. The data are shown as mean ± SEM (n = 3×3; $, *p*< 0.01 compared with the transfection with *P_Nrf1_-luc* and pcDNA3). (C) *Nrf1/2^+/+^* cells were co-transfected either *P_Nrf1_-luc* or *P_Nrf2_-luc,* together with *pRL-TK*, and allowed for 24-h recovery, before being treated with 50 μM tBHQ (*tert*-butylhydroquinone), 1 μM TG (thapsigargin) or 200 μM VC (vitamin C) for additional 24h, respectively. The data are shown as mean ± SEM (n = 3×3; $, *p*< 0.01. NS= no statistical difference). (D) Either *P_Nrf1_-luc* or *P_Nrf2_-luc,* plus *pRL-TK* and an expression construct for Nrf1 or Nrf2 at the concentrations as indicated were co-transfected into *Nrf1/2^+/+^* cells and then allowed for 24-h recovery before being determined. The data are shown as mean ± SEM (n = 3×3; \**p*< 0.01). (E) The pulse-chase analysis of Nrf2 in *Nrf1/2^+/+^* and *Nrf1α^−/−^* cells were carried out after treatment of these cells with 50 μg/mL of cycloheximide (CHX) alone or plus 5 μM of proteasomal inhibitor MG132 for various lengths of time as indicated. (F) The stability of Nrf2 was determined with its half-life in *Nrf1/2^+/+^* and *Nrf1α^−/−^* cells as treated above (*E*). (G) Expression of *GSK3β, β-TrCP, p62* and *Keap1* at mRNA levels in *Nrf1/2^+/+^* and *Nrf1α^−/−^* cells were examined. The qPCR data are shown as mean ± SEM (n = 3×3, \**p*< 0.01). (H) The protein abundances of Keap1, GSK3β and β-TrCP in *Nrf1/2^+/+^* and *Nrf1α^−/−^* cells were determined by Western blotting. (I) The mRNA (*upper column*) and protein (*lower panel*) levels of PTEN in *Nrf1/2^+/+^, Nrf1α^−/−^* and *caNrf2^ΔN^* cells were determined. The data are shown as mean ± SEM (n = 3×3, \**p*< 0.01, NS= no statistical difference). (J) *Nrf1/2^+/+^* and *Nrf1α^−/−^* cells had been treated with 100 nM rapamycin (RAPA) for 24 h, before being visualized by Western blotting to detect the changes of p-S6K1, AKT1, Nrf1, Nrf2, HO-1, and COX2 proteins. (K) An inter-regulatory model is proposed to explain opposing and unifying cross-talks between Nrf1 and Nrf2 at distinct levels.

### *Nrf1α^−/−^*-leading accumulation of Nrf2 results from decreased Keap1

The putative inter-regulation between Nrf1α and Nrf2 was further investigated to interpret the rationale underlying an abnormal accumulation of Nrf2 protein with no changes in its mRNA expression in *Nrf1α^−/−^* cells (Figure 2G,4A). Based on this finding, combined with the nation that Nrf1, but not Nrf2, acts as a primary transcriptional regulator of 26S proteasomal subunits (Radhakrishnan et al., 2010; Steffen et al., 2010), thereby it is hypothesized that aberrant accumulation of Nrf2 results from loss of Nrf1α’s function leading to an imbalance between Nrf2 protein synthesis and degradation processing. As shown in Figure S7(B, C), total protein ubiquitination was significantly accumulated in *Nrf1α^−/−^* cells, but not in *Nrf2^−/−ΔTA^* cells, when compared with wild-type cells. Further analysis of mRNA expression levels revealed that 20 of 36 genes encoding all 26S proteasomal subunits and relevant proteins were significantly reduced by knockout of Nrf1α (with only an exception of *PSMB10* enhanced) (Figure S7D). By contrast, no marked changes in transcriptional expression of 31 of the above 36 genes (except that *PSMB3, PSMB10, PSMC5,* and *PSMD3* were reduced and *PSME1* increased) were determined in *Nrf2^−/−ΔTA^* cells (Figure S7E). Therefore, such proteasomal dysfunction by loss of Nrf1α may result in an accumulation of Nrf2 by impaired proteasomal degradation pathway.

To address this, Nrf2 protein turnover was further determined by pulse-chase analysis of its half-life in *Nrf1α^−/−^* cells (Figure 5E). Surprisingly, it was found that *Nrf1α^−/−^* cells gave rise to relatively stable protein of Nrf2 with a prolonged half-life to 2.71 h (=163 min) after treatment with cycloheximide (CHX, an inhibitor of newly-synthesized polypeptides), and also such longevity of Nrf2 was largely unaffected by the proteasome inhibitor MG132 (Figure 5F, *lower panel*). As controls, *Nrf1^+/+^* cells displayed rapid turnover of Nrf2 with a short half-life of 0.38 h (=23 min) after CHX treatment, and this lifetime was extended to 1.17 h (= 70 min) by MG132 (Figure 5F, *upper panel*). Overall, the aberrant accumulation of Nrf2 in *Nrf1α^−/−^* cells results from impaired proteasome-mediated degradation.

Next, several upstream regulators of Nrf2 were examined to determine which pathways are impaired towards its protein turnover in *Nrf1α^−/−^* cells. Intriguingly, an abundance of Keap1 protein was significantly decreased (Figure 5H), even though its mRNA expression levels were unaltered, along with its turnover regulator *p62* was strikingly reduced in *Nrf1α^−/−^* cells (Figure 5G). Thereby, turnover of Keap1 may also occur through a p62-independent pathway. As such, impairment of Keap1-mediated proteasomal degradation of Nrf2, in particular, oxidative stress (Kobayashi et al., 2006), is likely to contribute to an aberrant accumulation of Nrf2 by the loss of Nrf1α. However, accumulation of Nrf2 is attributable to another impairment of GSK3β-phosphorylated β-TrCP-mediated proteasomal degradation of the CNC-bZIP protein. This is due to a marked decrease of GSK3β at mRNA and protein levels in *Nrf1α^−/−^* cells (Figure 5G,5H).

### Nrf1α and Nrf2 exert opposing and unifying roles in the regulation of PTEN signaling

More importantly, we found that both protein and mRNA levels of PTEN (acts as a key master versatile regulator of Nrf2, Keap1, PI3-kinase, AKT and GSK3β (Pitha-Rowe et al., 2009; Rojo et al., 2014; Sakamoto et al., 2009; Taguchi et al., 2014), were significantly diminished or even abolished in *Nrf1α^−/−^* cells (retaining high expression of Nrf2) (Figure 5I, *left panel*). In contrast, inactivation of Nrf2 caused a striking increase in PTEN mRNA, but not its protein levels, in *Nrf2^−/−ΔTA^* cells (yielding low expression of Nrf1) (Figure 3F, 4F). On the contrary, in *caNrf2^ΔN^* cells (giving rise to a high expression of Nrf2 and Nrf1, Figure 4I), a significant decrease in PTEN protein abundance, but not its mRNA levels, was determined (Figure 5I). Collectively, together with the above-described data (Figure 3), both Nrf1α and Nrf2 are much likely to exert opposing and unifying roles in the precision regulation of PTEN by both miR-22-dependent and -independent pathways, in which Nrf2 is preferably dominant-negative, whereas Nrf1α has a limited positive role.

Further analysis of the *PTEN* gene unraveled that there exist two typical ARE sites within its promoter region (Figure 3G). The luciferase assays demonstrated that transcription activity of the *PTEN* promoter-driven luciferase reporter *P_PTEN_-luc* was significantly induced by Nrf1, but not by Nrf2 (Figure 3H). Mutagenesis analysis uncovered that the second ARE2 site made a primary contribution to transactivation activity of the *P_PTEN_-luc* reporter by Nrf1, while the first ARE1 site also gained a minor contribution to Nrf1-mediated transactivation of *P_PTEN_-luc* (Figure 3I).

Based on the fact that loss of PTEN function leads to constitutive activation of the PI3K-AKT signaling pathway to augment the nuclear accumulation of Nrf2 and its resulting activation (Mitsuishi et al., 2012; Sakamoto et al., 2009), we determined whether the PI3K-AKT signaling is activated by abolishment of PTEN in *Nrf1α^−/−^* cells (in which Nrf2 is aberrantly accumulated). The results demonstrated *Nrf1α^−/−^-*leading increased abundances of Nrf2, AKT1, COX2 and HO-1, but such increases were significantly suppressed by the classic mTOR inhibitor, rapamycin (RAPA) (Figure 5J). This implies that the mTOR may also be activated in *Nrf1α^−/−^*cells. Accordingly, increased abundances of both AKT and phospho-S6K1 in *Nrf1α^−/−^* cells were markedly blocked by mTOR inhibitor RAPA. This is inversely correlative with the consequence that over-expression of Nrf1 suppresses AKT induction (Hirotsu et al., 2014). Together, this study indicates opposing and unifying cross-talks between Nrf1α and Nrf2 to regulate the PTEN-mTOR-AKT signaling to the Nrf2-COX2 pathway. Based on these findings, we summarized an endogenous inter-regulatory network (Figure 5K).

### Blockage of *Nrf1α ^+/+^*-bearing or *Nrf1α^−/−^*-derived tumor growth by Nrf2 deficiency

Our previous work revealed that malignant growth of *Nrf1α^−/−^*-derived hepatoma is accompanied by metastasis to the liver in xenograft mice (Ren et al., 2016). Here, to elucidate what effects have been elicited by Nrf1α and Nrf2 on tumor repression or promotion, we further investigate distinct genotypic tumors derived from *Nrf1/2^+/+^, Nrf1α^−/−^, Nrf1α^−/−^+siNrf2, Nrf2^−/−ΔTA^* and *caNrf2^ΔN^* cells in xenograft mice. Their tumorigenicity was evaluated by measuring tumor volume and weight. As illustrated in Figure 6(A to C), *Nrf2^−/−ΔTA^* cells were inoculated in nude mice, but did not form more than one solid tumor. This implies that tumorigenicity of *Nrf1/2^+/+^* cells, as controls, is almost completely abolished by inactivation of Nrf2. Conversely, constitutive activation of Nrf2 did not obviously influence the resulting *caNrf2^ΔN^*-driven tumorigenicity, by comparison to that of *Nrf1/2^+/+^*. This indicates that Nrf2-prone cancer promotion is dominantly confined by the presence of Nrf1α. Upon loss of Nrf1*α* function in *Nrf1α^−/−^* cells (in which hyper-active Nrf2 is accumulated), the resultant tumorigenicity was hence significantly higher than that of *Nrf1/2^+/+^*-tumor, but rather was much strikingly suppressed by silencing of Nrf2 (in the *Nrf1α^−/−^+siNrf2*-derived tumors) to the much less extent than that of *Nrf1/2^+/+^* cells (Figure 6, A to C). Collectively, these findings demonstrate that Nrf2 acts as a tumor promoter, but it is efficiently confined by Nrf1α serving as a dominant tumor repressor, implying both are a pair of mutual antagonizing twin factors. Overall, malignant transformation of *Nrf1α^−/−^*-derived cells is attributable to hyperactivation of Nrf2.

**Figure 6.**
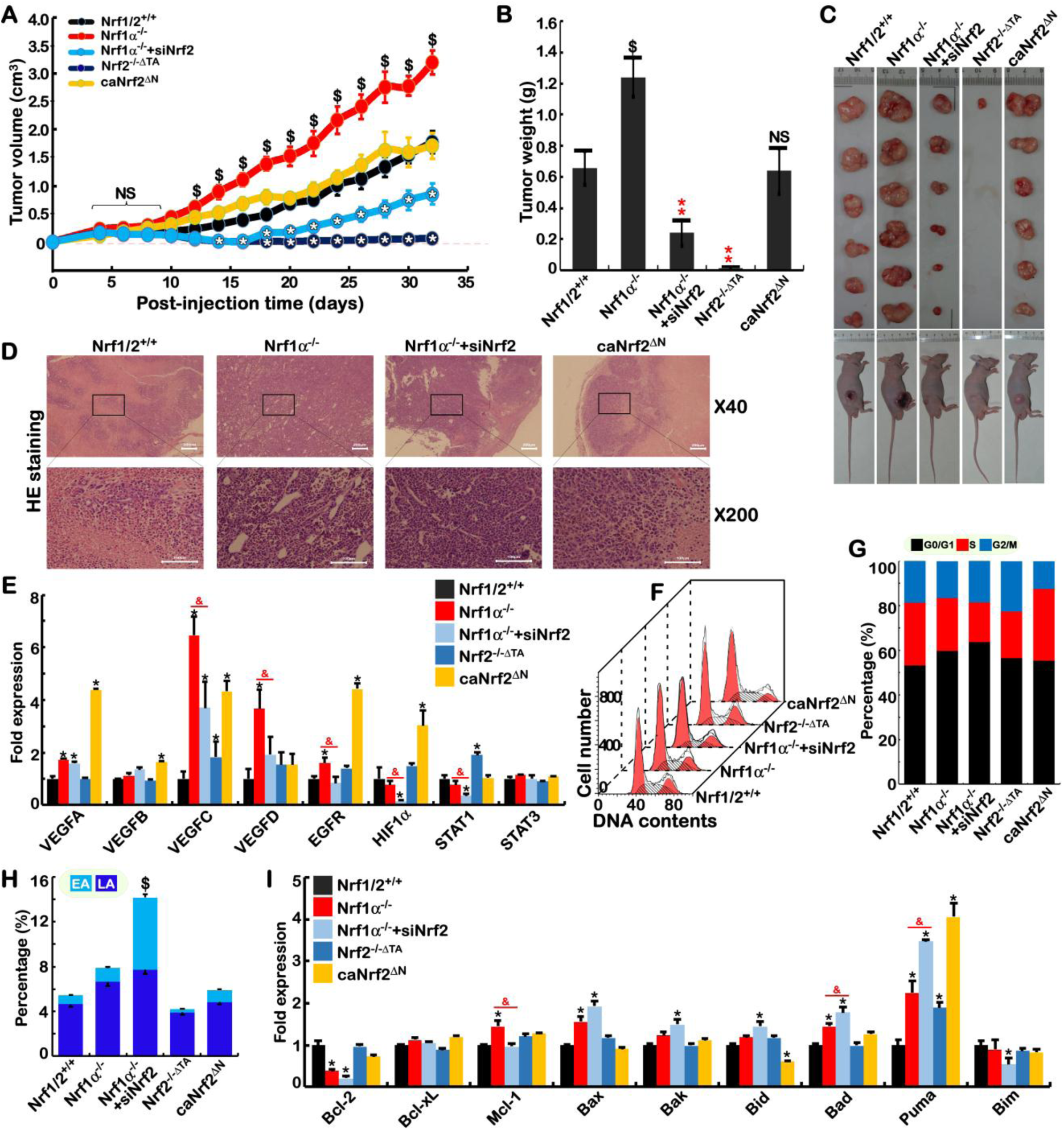
Distinct animal tumor phenotypes derived from *Nrf1α^−/−^, Nrf1α^−/−^*+siNrf2, *Nrf2^−/−ΔTA^, caNrf2^ΔN^* from *Nrf1/2^+/+^* cells. (A) Distinct mouse subcutaneous xenograft tumors derived from *Nrf1/2^+/+^, Nrf1α^−/−^ Nrf1α^−/−^*+siNrf2, *Nrf2^−/−ΔTA^* and *caNrf2^ΔN^* cells were measured in size every two days, before being sacrificed on the 32^nd^ day. The data are shown as mean ± SEM (n = 6 per group, \**p*< 0.01; $, *p*< 0.01, NS= no statistical difference at the early incubation phase). (B) The data final tumor weight of tumors were calculated m and are shown as mean ± SEM (n = 6, \*\**p*< 0.01; $, p< 0.01, NS= no statistical difference, when compared to the wild-type). (C) Distinct animal xenograft tumors derived from *Nrf1/2^+/+^, Nrf1α^−/−^ Nrf1α^−/−^*+siNrf2, *Nrf2^−/−ΔTA^* and *caNrf2^ΔN^* cells. (D) The histological photographs of indicated tumors were achieved by HE (hematoxylin & eosin) staining. The scale bar = 200 μm in ×40 pictures, or = 100 μm in ×200 pictures. (E) The qPCR analysis of some angiogenesis-related genes in distinct cells as indicated was validated by transcriptome. The data are shown as mean ± SEM (n= 3×3, \**p*< 0.01, \*\**p*< 0.001; $, *p*< 0.01; $$, *p*< 0.001). (F and G) The flow cytometry analysis of distinct cell cycle was indicated. The data (n = 3) are shown in two different fashions (H) The early apoptosis (EA) and late apoptosis (LA) of five distinct cell lines was examined by flow cytometry. The data are shown as mean ± SEM (n = 9; $, *p*< 0.01). (I) Expression of some apoptosis-related genes in different indicated cells was transcriptomically analyzed. The data are shown as mean ±

Histological examination showed that a considerable number of blood vessels were formed in *Nrf1α^−/−^* tumors but was reduced by Nrf2 knockdown in *Nrf1α^−/−^+siNrf2*-derived tumors (Figure 6D). However, no marked differences in vascularity between *caNrf2^ΔN^*-and *Nrf1/2^+/+^*-bearing tumors were observed. Further insights into angiogenesis-related genes revealed that mRNA expression levels of *VEGFA, VEGF*C, *VEGFD, EGFR,* but not *HIF1α* or *STAT1* were strikingly elevated by knockout of Nrf1α, but the increased expression of *VEGF*C, *VEGFD and EGFR* was significantly reduced by silencing Nrf2 (Figure 6E). Notably, knockdown of Nrf2 almost completely abolished expression of *HIF1α* and *STAT1* in *Nrf1α^−/−^+siNrf2* cells, but no changes in these two factors were observed in *Nrf1α^−/−^* cells, as compared to those obtained from *Nrf1/2^+/+^* cells. Rather, all the other angiogenesis genes except *VEGFD* were up-regulated in *caNrf2^ΔN^* cells (giving high expression of Nrf1 and Nrf2), while only *STAT1* but not other genes were up-regulated by inactive *Nrf2^−/−ΔTA^* (Figure 6E). Altogether, both Nrf1α and Nrf2 are diversely involved in regulating the expression of angiogenesis genes except for *STAT3* as examined above.

Intriguingly, the vascularity of *Nrf1α^−/−^+siNrf2*-derived tumors seemed to be higher than that *Nrf1/2^+/+^*-bearing tumors (Figure 6D), but, such angiogenetic changes cannot explain the observation that the *Nrf1α^−/−^+siNrf2*-tumor volumes and weights were significantly less than those obtained from the *Nrf1/2^+/+^*-tumors. This implicates other rationales beyond angiogenesis. Thus, we employed flow cytometry to determine changes in the cell cycle and apoptosis in five distinct cell lines. As shown in Figure 6(F, G), the S-phase of *Nrf1α^−/−^+siNrf2* cells was significantly shortened. Such a severe S-phase arrest of cell cycle is supported by quantitative gene expression analysis revealing that significant up-regulation of *p16, p19, p21 p53* and *CDK4* was simultaneously accompanied by down-regulation of *RB1, CDK1, CyclinA2, CyclinB2, E2F3, E2F5,* and *E2F6* in *Nrf1α^−/−^+siNrf2* cells, when compared with its progenitor *Nrf1/2^+/+^* or *Nrf1α^−/−^* cells (Figure S8A). In addition to the S-phase arrest, the G0/G1-phase was relatively extended in *Nrf1α^−/−^+siNrf2* cell cycle (Figure 6G). Consistently, apoptosis of *Nrf1α^−/−^+siNrf2* cells was also significantly enhanced, when compared to other cell lines (Figure 6H, S8D). This is also supported by further analysis of apoptosis-related genes unraveling that *Bax, Bak, Bid, Bad*, and *Puma* were significantly up-regulated, while anti-apoptotic *BCL-2* gene was down-regulated, with no changes in *BCL-xL* and *Mcl-1* in *Nrf1α^−/−^+siNrf2* cells (Figure 6I).

Although no significant differences in growth and vascularity of between *caNrf2^ΔN^*-and *Nrf1/2^+/+^*-bearing tumors, the G2/M-phase of the *caNrf2^ΔN^* cell cycle was shortened, along with the S-phase extended (Figure 6G). This implies that a G2/M-phase arrest is likely caused by the constitutive activation of Nrf2, in agreement with the supportive evidence that inactivation of Nrf2 markedly prolonged the G2/M-phase of *Nrf2^−/−ΔTA^* cells (Figure 6G). Further gene expression analysis revealed that *p15, p21* and *Puma* were significantly up-regulated, but *p18, CDK1, E2F2* and *Bid* were down-regulated by *caNrf2^ΔN^* (Figure 6I, S8A). Conversely, inactive *Nrf2^−/−ΔTA^* still up-regulated *RB1, CDK1, E2F3*, and *Cyclin D1* (Figure S8A), but strikingly down-regulated *FTH1* and *FTL* (both encode ferritin heavy and light chains involved in iron-dependent lipid peroxidation and ferroptosis, in Figure S8G). Overall, these demonstrate that Nrf1α and Nrf2 coordinately regulate certain key genes involved in cell cycle and apoptosis.

SEM (n=3, \**p*< 0.01, ** *p*< 0.001; $, *p*< 0.01; $$, *p*< 0.001).

### Different subsets of genes are finely regulated by Nrf1α, Nrf2 alone or both

Nrf1 and Nrf2 are two important CNC-bZIP transcription factors that are widely expressed in various tissues and also regulate seemingly similar expression patterns of ARE-driven downstream genes that have been identified (Bugno et al., 2015; Tebay et al., 2015). In fact, the ever-accumulating evidence demonstrates that Nrf1 and Nrf2 exert many different and even opposing functions and in particular, unique indispensable functions of Nrf1 are not substituted by Nrf2 (Zhang and Xiang, 2016). Accordingly, the above-described data also unraveled that both CNC-bZIP factors have elicited mutual synergistic and antagonistic roles in regulating precision expression of cognate genes in distinct cell processes, aiming to maintain normal cellular homeostasis. Here, to further evaluate the functional similarities and differences between Nrf1α and Nrf2, the genome-wide expression of genes in *Nrf1/2^+/+^, Nrf1α^−/−^, Nrf1α^−/−^+ siNrf2, Nrf2^−/−ΔTA^* and *caNrf2^ΔN^* cells was determined by transcriptome sequencing. Those detectable genes with a fold change ≥ 2 and diverge probability ≥ 0.8 were defined as differentially expressed genes (DEGs), by comparison with equivalents measured from *Nrf1/2^+/+^* cells (Figure 7A). Consequently, *Nrf1α^−/−^* cells gave rise to 1,213 of DEGs (i.e. 697 up-plus 850 down-regulated), but the number of DEGs in *Nrf1α^−/−^+siNrf2* cells was significantly increased to 3,097 genes, 2247 of which were down-regulated by siNrf2. Intriguingly, only 545 of DEGs were detected in *Nrf2^−/−ΔTA^* cells, implying that many genes are silenced or prevented by inactive Nrf2 mutant (distinctive from simple knockout of this factor). These data suggest that, in this regulatory system by the cooperation of Nrf1 and Nrf2, a single change of both has only limited effects on overall gene expression, and thus both changes will have a greater impact. For instance, when compared to those of *Nrf1α^−/−^* cells, silencing of Nrf2 caused 124 genes to be up-regulated, and still led 1,338 genes to be down-regulated in *Nrf1α^−/−^+siNrf2* cells (Figure 7A, *last column*), so that malignant growth of *Nrf1α^−/−^*-derived tumor was repressed by Nrf2 knockdown. Conversely, reinforced expression of Nrf2 (and Nrf1) in *caNrf2^ΔN^* cells up-regulated 1655 genes and also down-regulated 423 genes. These findings indicate that Nrf2 is a dominant activator to regulate many genes, while Nrf1α appears to exert dominant negative effects on some genes.

**Figure 7.**
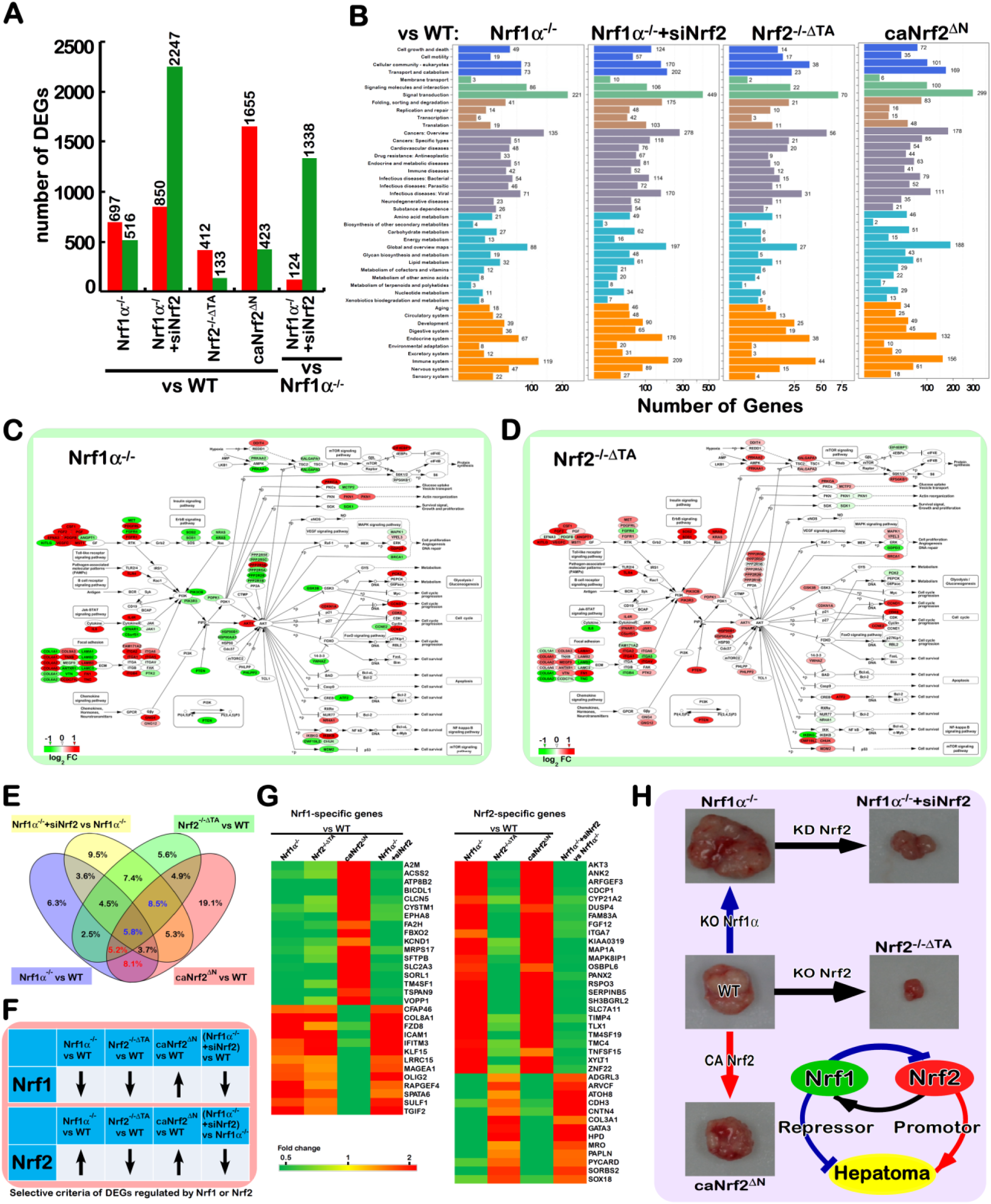
An axiomatic rationale underlying distinct animal xenograft tumor phenotypes. (A) Differentially expressed genes (DEGs) in distinct cell lines were analyzed by transcriptome sequencing. The differences in the number of DEGs are shown after being compared with wild-type or indicated cells. Those increased or decreased DEGs were represented by red or green columns, respectively. The DEGs were selected according to the following criteria: fold change ≥ 2 or ≤ 0.5 and diverge probability ≥ 0.8 (as compared to the control group). (B) KEGG classification of DEGs for each pairwise. The X-axis shows the number of DEGs, while the Y-axis represents distinct second-grading KEGG pathways. The top pathways are shown in different colors, such as cellular processes (*blue)*, metabolism (l*ight blue*), environmental information processing (*green*), genetic information processing (*brown*), human disease (*purple*), and organism system (*orange*). (C, D) Significant differences in the DEGs enriched responsible for the PI3K-AKT signaling pathway in *Nrf1α^−/−^* and *Nrf2^−/−ΔTA^* cells. (E) The Venn diagram shows the DEGs in four single variable group. To expand the screening range, the DEG is redefined as a fold change greater than 1.5 or less than 0.66. Either Nrf1α-specific or Nrf2-specific downstream genes were indicated by red and green numbers. (F) Distinct changes in abundances of Nrf1α and Nrf2 were illustrated after both protein levels present in each indicated cell was compared with the equivalent values from wild-type cells. (G) The Heat maps of particularly Nrf1α-and Nrf2-specific downstream genes, which were screened from the transcriptome data in this experimental setting. (H) An explicit model is proposed to decipher the axiomatic rationale underlying distinct animal xenograft tumor phenotypes, demonstrating significant differences in the cancer pathobiology of between Nrf1α and Nrf2.

Enrichment analysis revealed that DEGs of *Nrf1α^−/−^* cells were subject to 16 pathways (*p* < 0.001), of which 9 are responsible for human disease and 4 are involved in the environmental information processing (Table S1). By contrast, most of the cellular processes were significantly changed in *Nrf2^−/−ΔTA^* and *Nrf1α^−/−^+siNrf*2 cells. Thus, loss of Nrf1α relevant to the disease suggests that its function is essential for maintaining cellular homeostasis, while Nrf2 exerts its greater roles in regulating most of cellular physiological processes. For example, the above-described alterations in the cells cycle of *Nrf1α^−/−^+siNrf2* were also validated by transcriptome (Table S1). Further calculation of the DEGs distribution unraveled that signal transduction, cancer-relevant, immune system and metabolism were most abundant secondary KEGG pathways in Nrf1-or Nrf2-deficient cells (Figure 7B). An insight into the cellular signaling transduction uncovered that the most DEGs are involved in the PI3K-AKT pathway (Table S1). In this pathway, a key tumor suppressor PTEN was significantly and oppositely altered in both *Nrf1α^−/−^* and *Nrf2^−/−ΔTA^* cell lines (Figures 3F, 7C, 7D). Based on these specific findings, much-focused DEGs in *Nrf1α^−/−^* and *Nrf2^−/−ΔTA^* cells were mapped according to the KEGG pathway. The results illustrated that both cell lines displayed such significant opposing changes in the PI3K-AKT pathways (Figures 7C, 7D, S9A). As interested, knockout of *Nrf1α* (with accumulated Nrf2) caused a general reduction in transcription of most AKT-signaling molecules, but they were hence generally increased by inactivation of *Nrf2*. Such striking disparity is dictated by the distinction of Nrf2 proteins in between the two cell lines (Figure 2).

Notably, albeit seemingly similar downstream genes are regulated by Nrf1 and Nrf2, *de facto* activation of Nrf2 by knockout of *Nrf1α* can inevitably cause their opposite effects on some genes against theoretic expectations. This is further evidenced by results from *Nrf1α^−/−^+siNrf2* cells, revealing that many of those accumulated Nrf2’s effects on downstream genes by *Nrf1α^−/−^* were strikingly reduced by knockdown of Nrf2. Therefore, by comparison of the DEGs between *Nrf1α^−/−^* and *Nrf1α^−/−^+siNrf2* cell lines, an opposite expression profiling of 87 genes was uncovered by Nrf2 knockdown (Figure S9B-D). About 24 % of these genes are responsible for metabolism-related enzymes. This implies that the function of Nrf2 is closely related to cellular metabolism, particularly in the absence of Nrf1α. This is further approved by another opposite expression profiling of other 83 DEGs in between *Nrf2^−/−ΔTA^* and *caNrf2^ΔN^* cell lines (Figure S10). Still 16% of the differential expression genes are related to metabolism process, but 24% of these genes are involved in signaling transduction. This observation indicates that in the presence of Nrf1α, Nrf2 is a major player in cellular signaling cascades, but its role in metabolism appears to be restricted possibly by Nrf1α.

The Venn diagrams illustrated that distinct subsets of DEGs were regulated by Nrf1, Nrf2 alone or both (Figure 7E). The common genes regulated by Nrf1α and Nrf2 were seen by comparison of DEGs in *Nrf1α^−/−^* or *caNrf2^ΔN^* with wild-type. In the intersection of *Nrf1α^−/−^* and *caNrf2^ΔN^*, the remaining portions after excluding *Nrf1α^−/−^+siNrf2* with *Nrf1α^−/−^* were composed of the (*red numbered*) genes closely correlated to regulation by Nrf1α. The genes regulated by Nrf2 were also found by comparison of DEGs in *Nrf1α^−/−^+siNrf2* with *Nrf1α^−/−^*, as well as *Nrf2^−/−ΔTA^* or *caNrf2^ΔN^* with wild-type, so the intersection of these three sets comprised the (*blue numbered*) genes preferably regulated by Nrf2. Further, based on the changes in Nrf1 and Nrf2 proteins detected in distinct cell lines (Figure 7F), we screened which portions of high-relevant downstream genes were consistent with or opposite to the changing trends of Nrf1 or Nrf2, respectively. Consequently, 30 Nrf1α-specific downstream genes were shown (in Figure 7G, *left panel*), of which 17 genes were up-regulated and 13 genes were down-regulated. Meanwhile, 38 Nrf2-specific downstream genes were also found herein, of which 25 were up-regulated and 13 were down-regulated (*right panel*). Collectively, our findings provide an axiomatic rationale for differential expression of different subsets of genes to dictate distinct phenotypes of animal xenograft tumors (Figure 7H). Significantly, the malfunction of Nrf2 is as a potent tumor promoter, but it can be efficiently confined or suppressed by Nrf1α acting as a dominant tumor repressor.

## DISCUSSION

Accumulating evidence has demonstrated that Nrf1 is a key player in the pathogenesis of NASH and HCC, as well as other relevant cardiovascular diseases and type 2 diabetes (Bugno et al., 2015; Zhang and Xiang, 2016). However, it should be noted that these experimental mouse genomes were manipulated to delete all Nrf1 isoforms from the single *Nrf1/Nef2l1* gene. In this study, human *Nrf1α*-specific knockout was achieved by the gene-editing to create the frameshift mutation. The phenotypes of NASH and malignancies were reconstructed by using monoclonal *Nrf1α^−/−^* cell lines. Thereby, this provides an available model for a follow-up study to elucidate the relevance of Nrf1α with NASH and its malignance into HCC. In the *Nrf1α^−/−^*-leading model, the inflammation marker COX2 is constitutively increased, which entails a non-resolving feature. By contrast, the development-related COX1 was almost completely abolished in *Nrf1α^−/−^* cells. The resultant metabolites of arachidonic acid by the rate-limiting enzyme COX2, which also serves as a direct target of Nrf2 (Itoh et al., 2004; Sherratt et al., 2003), are much likely to play a crucial role in the development and progression of inflammation, particularly NASH and hepatoma caused by knockout of *Nrf1α*.

Further examinations revealed that the *Nrf1α^−/−^*-caused increase of COX2 occurred by accumulated Nrf2 protein, but both were effectively diminished by inhibitors of JNK (i.e. SP600125) and mTOR (i.e. rapamycin). Hence, the Nrf2-COX2 pathway is inferable to be regulated by both JNK and mTOR signaling, albeit the detailed mechanisms remain unclear. Here, we found that inhibition of the Nrf2-COX2 pathway is accompanied by decreases in AKT, S6K1 and GSK3β. This is consistent with the claim that Nrf2 is regulated by the mTOR-AKT-GSK3β pathway (Yang et al., 2018). Our findings also unravel that Nrf2 may be monitored by JNK signaling towards AP-1 pathway, but in turn, some AP-1 components (i.e. Jun, Fra-1) are mediated by Nrf2 insofar as to form a feedback loop. Contrary to *Nrf1α^−/−^*, MEFs of *Nrf1^−/−(ΔDBD)^*, in which almost all DBD (DNA-binding domain)-containing Nrf1 isoforms are disrupted (Hirotsu et al., 2012; Ohtsuji et al., 2008; Tsujita et al., 2014; Xu et al., 2005), exhibited marked decreases in total COX2 and most of Nrf2 to much lower levels roughly similar to those determined in *Nrf2^−/−(ΔDBD)^* MEFs. (Figure S5C). This difference between human *Nrf1α^−/−^* and mouse *Nrf1^−/−(ΔDBD)^* demonstrates Nrf1 isoform-dependent regulation of the Nrf2-COX2 pathway in distinct species as experimented.

Notably, an accumulation of free radicals in *Nrf1^−/−(ΔDBD)^* MEFs results from decreased expression of ARE-driven genes involved in glutathione synthesis, antioxidant and detoxification (Kwong et al., 1999). Similar but different stress caused by liver-specific knockout of *Nrf1^−/−(ΔDBD)^* activates a subset of Nrf2-dependent ARE-battery genes in mice, but Nrf2 cannot still compensate for the loss of Nrf1’s function leading to NASH and HCC (Ohtsuji et al., 2008; Xu et al., 2005). The inducible liver-specific knockout of *Nrf1^−/−(ΔDBD)^* in mice increased glutathione levels; this results from up-regulation of *xCT* (a component of the cystine/glutamate antiporter system *X_C_*^-^), but with no changes in glutathione biosynthesis enzymes (Tsujita et al., 2014). In this work, human *Nrf1α^−/−^* increases ROS and lipid levels, along with high expression of *xCT* and other ARE-driven genes (e.g. *HO-1, GCLC, GCLM, NQO1*). These genes are Nrf2-dependent because their expression is reduced by inactive *Nrf2^−/−ΔTA^* mutant and also repressed by silencing of Nrf2 (in *Nrf1α^−/−^ +siNrf2* cells). In addition to COX1 and COX2, both Alox5 and FLAP (also involved in arachidonic acid metabolism) are significantly up-regulated in *Nrf1α^−/−^* cells, and also modestly increased in *Nrf2^−/−ΔTA^* cells. However, liver-specific *Nrf1^−/−(ΔDBD)^* mice display no changes in *COX1, COX2* and *Alox5* expression (Tsujita et al., 2014). Overall, these discrepancies are likely attributed to the variations of which Nrf1 isoforms have two-sided effects on Nrf2 and diverse downstream genes, depending on different cell types in distinct species. Hence, it is crucially important to determine the *bona fide* effects of Nrf1 and Nrf2 alone or in combination on distinct cognate genes within regulatory networks (Figure 5K).

Albeit Nrf1 and Nrf2 are recruited for directly binding the ARE sites in the *COX1* and *COX2* promoter regions (Itoh et al., 2004; Sherratt et al., 2003), our evidence unravels that both CNC-bZIP factors have different or opposing roles in bi-directional regulation of *COX1* and *COX2* by distinct interrelated positive and negative pathways. In particular, Nrf1α has a two-handed potency to execute as an activator or repressor, depending on distinct cognate genes (e.g. *COX1* and *COX2*), through different regulatory pathways. For example, regulation of *COX2* is contributed positively by Nrf2 and negatively by Nrf1α, albeit its promoter-driven *P_COX2_-Luc* reporter is also transactivated by Nrf1 and Nrf2. The direct activation by Nrf1 (as well as Nrf2) may be neutralized or counteracted by its dominant-negative effects triggered by indirect mechanisms (as shown in Figure 5K).

Just contrary to *COX2*, expression of *COX1* is regulated positively by Nrf1α but negatively by Nrf2, albeit no direct activation of its promoter-driven *P_COX1_-Luc* reporter by ectopic Nrf1 and Nrf2 was detected herein. In an attempt to explore into the mechanisms by which COX1 is indirectly regulated by Nrf1/2, we found that both CNC-bZIP factors can directly activate the expression of miR-22 driven by its ARE site. The miR-22, along with Nrf1 and Nrf2, all inhibit the *PTEN-miR22b-Renilla* reporter activity, implying that these two factors have an intrinsic ability to suppress the tumor suppressor PTEN through activating miR-22, as consistent with the previous reports (Tan et al., 2012; Xu et al., 2012). However, *de facto* endogenous expression of PTEN at mRNA and protein levels is almost completely abolished in *Nrf1α^−/−^* cells (retaining hyper-active Nrf2), but also dramatically increased in *Nrf2^−/−ΔTA^* cells (with decreased Nrf1). Collectively, these findings demonstrate that Nrf2 is a dominant negative to inhibit PTEN; this is further evidenced by a significant reduction of PTEN by *a priori* constitutive activation of Nrf2 in *caNrf2^ΔN^*cells (also with enhanced Nrf1). By contrast, Nrf1α has Ying-Yang two-sided effects on PTEN. On one side, Nrf1α acts as a major positive regulator of PTEN, while on the other side of Nrf1α, it is enabled to exert a minor negative role in PTEN, but this negation could be concealed by dominant negative Nrf2 or counteracted by the major positive action of Nrf1α *per se*.

Since PTEN is known to act as the most critical inhibitor of the PI3K-AKT pathway (Tan et al., 2012; Xu et al., 2012), thereby, inactivation of PTEN by ROS provokes activation of its downstream PI3K-AKT signaling cascades to promote cell survival (Chetram et al., 2011; Kitagishi and Matsuda, 2013). Notably, the ever-increasing evidence demonstrates that PTEN can direct inhibition of expression of ARE-driven gene by inhibiting Nrf2 (Rojo et al., 2014; Sakamoto et al., 2009). Taken together with our results from this study, it is demonstrated that the cellular ROS levels are monitored by Nrf1α-and/or Nrf2-mediating ARE-battery genes, but in turn, expression of Nrf1/2-target genes is also negatively regulated by ROS-activated mir22-PTEN signaling to form a feedback regulatory circuit. Of note, activation of Nrf1/2 by ROS can promote miR-22 expression, which may serve as an important approach to regulate the PTEN-PI3K-AKT pathway. Thereby, the quantitative regulations of cellular ROS levels are achieved by close cooperation of Nrf1α and Nrf2 coordinately through direct and indirect mechanisms, so as to maintain normal redox homeostasis. Conversely, dysfunction of Nrf1α and Nrf2 (particularly its malfunction) leads to severe redox stress and cancer development possibly by the aberrant PTEN-PI3K-AKT signaling pathway.

It is inferable that almost abolishment of PTEN in malignantly growing *Nrf1α^−/−^*-derived tumor cells results from an aberrant accumulation of Nrf2 protein, because rescue of PTEN expression occurs after Nrf2 is silenced, such that the existing *Nrf1α^−/−^+siNrf2*-derived tumor growth is dramatically repressed by knockdown of Nrf2. In turn, the aberrant accumulation of Nrf2 in *Nrf1α^−/−^*cells is caused by impaired PTEN expression. This is fully consistent with the pathology of *PTEN^−/−^*-leading cancer, in which the abnormal nuclear accumulation of Nrf2 is caused by impairment of GSK3β-directed β-TrCP-based proteasome-mediated degradation, as described by (Best et al., 2018; Rojo et al., 2014; Taguchi et al., 2014). In addition to impairment of the GSK3β-directed β-TrCP pathway, aberrant accumulation of Nrf2 is augmented by inhibition of Keap1-based proteasome-mediated degradation in *Nrf1α^−/−^*-derived tumor cells. Noticeably, the Keap1 protein, rather than mRNA, levels are significantly reduced in *Nrf1α^−/−^* cells, albeit its binding partner p62, acting as a major regulator of Keap1 to the autophagic degradation (Kageyama et al., 2018), is strikingly down-regulated in *Nrf1α^−/−^* cells. Thus, we postulate that a p62-independent mechanism may account for the Keap1 protein degradation and also is reinforced in *Nrf1α^−/−^* cells. Yet, it cannot be ruled out that biosynthesis of Keap1 polypeptides may also be retarded during these conditions.

Several lines of evidence presented here demonstrate that Nrf2 is predominantly negatively regulated by Nrf1α because *Nrf1α^−/−^* enables Nrf2 to be released from the confinements by the PTEN-GSK3β-directed β-TrCP-based and Keap1-based proteasomal pathways (Figure 5K). Consequently, accumulation of Nrf2 leads to aberrant activation of ARE-driven cytoprotective genes (e.g. *HO-1, GCLM, NQO1*) to shelter or promote *Nrf1α^−/−^*-driven tumor cells. In fact, These ARE-battery genes can be directly activated by Nrf1α, but some of these downstream genes could also be inhibited through braking control of the Nrf2 activity. Overall, distinct levels of Nrf1 alone or in cooperation with Nrf2 finely tune and also quantitatively regulate expression of diverse downstream genes to meet different cellular needs (Figures 4L, 5K). Thus, these resulting collective effects determine distinct phenotypes of animal xenograft tumor models as deciphered in this study (Figure 7H). Consistently, the malignant growth of *Nrf1α^−/−^*-derived tumor is substantially suppressed by knockdown of Nrf2, by comparison with *Nrf1α^−/−^+siNrf2*-derived tumor. Conversely, almost no solid tumor is formed in nude mice that have been inoculated by injecting inactive *Nrf2^−/−ΔTA^*-derived cells, albeit Nrf1 is slightly decreased with loss of Nrf2’s function. These demonstrate that Nrf1α acts as a dominant tumor suppressor principally by confining the oncogenicity of Nrf2. In turn, Nrf2 exerts a dominant tumor-promoting role in tumorigenesis and malignant growth, but it can also directly mediate the *Nrf1* gene transcription to form a feedback regulatory loop. This is validated by further evidence revealing that, upon the presence of Nrf1 in *caNrf2^ΔN^*-derived tumor cells, its growth is almost unaffected by constitutive activation of Nrf2, as well as antioxidant and detoxifying genes, when compared with wild-type *Nrf1/2^+/+^*-bearing tumor.

In an attempt to clarify those seemingly contradictory results obtained from loss of Nrf1α and its functional gain (i.e. ectopic over-expression), we have surprisingly found that there exists a mutual regulatory relationship between Nrf1α and Nrf2, thereby enabling both factors to elicit opposing and unifying roles in regulating distinct downstream genes (particularly ARE-driven cognate genes). Importantly, we have also discovered that that forced expression of Nrf1 enables the Nrf2 protein to be reduced, whereas loss of Nrf1α led to a significant increase in Nrf2 protein, but not its mRNA levels (Figure 4L). By contrast, both mRNA and protein levels of Nrf1 are increased by over-expression of Nrf2 or its constitutive activation, but also repressed by inactivation of Nrf2. Further experiments have unraveled no activation of the human *Nrf2* promoter-driven *P_Nrf2_-Luc* reporter by Nrf1 or Nrf2, albeit mouse *Nrf2* contains ARE sites as described (Kwak et al., 2002). However, the human *Nrf1* promoter-driven *P_Nrf1_-luc* reporter is *trans*-activated by Nrf1 (at the locus Site-2) and Nrf2 (at the locus Site-1) (Figures 5, S6A). These findings demonstrate there are two (i.e. transcript and protein abundance) levels at which Nrf1α and Nrf2 have cross-talks with each other to influence the expression of ARE-driven genes. Thereby, synergistic or antagonistic effects of Nrf1α and Nrf2 depend on mutual competition or somehow coordination with spatiotemporally binding to the same or different ARE enhancers within downstream genes. Overall, such inter-regulatory cross-talks between Nrf1α and Nrf2 could be a vitally important strategy for the precision regulation of distinct downstream genes. This rationale provides a better explanation of those complicated physio-pathological functions with distinct disease phenotypes exhibited in different models (as reviewed by Menegon et al., 2016; Zhang and Xiang, 2016).

Importantly, a hot controversy surrounds the roles of Nrf2 in the pro-or anti-cancer contexts, termed ‘the Nrf2 paradox’ (Menegon et al., 2016; Rojo de la Vega et al., 2018). This study has defined that function of Nrf2 is dictated by activation or inactivation of Nrf1α. This is because deterioration of *Nrf1α^−/−^*-tumor results from hyper-active Nrf2, along with decreased PTEN and activation of AKT signaling, but *Nrf1/2^+/+^*-tumor growth is unaffected by constitutive activation of Nrf2 when compared with *caNrf2^ΔN^*-tumor. Consistently, it has been recently showed that Nrf2 acts as a tumor-promoting player, depending upon aberrant activation of the PI3K-AKT signaling pathway, whereas it serves as a tumor-preventing player through activating ARE-driven cytoprotective genes under normal activation conditions (Best et al., 2018). However, a similar subset of ARE-driven genes is also highly expressed in *Nrf1α^−/−^* and *caNrf2^ΔN^* cell lines. Our findings demonstrate that the tumor-promoting role of Nrf2 is determined by loss of Nrf1α function, independent of those cytoprotective gene expressions. Even as a braking control of Nrf2 activity, Nrf1α may play a role for ‘decision-maker’ or ‘executor’ in the cell senescence and cancer progression, since a secretory phenotype of senescent cells occurs by a Nrf2-independent mechanism (Wang et al., 2017), albeit the relevance to Nrf1 needs to be verified.

In conclusion, this study provides a panoramic view of mutual inter-regulatory cross-talks existing between Nrf1α and Nrf2 to determine quantitative expression of distinct downstream genes involved in different patho-physiological processes. Significantly, the axiomatic rationale underlying distinct animal xenograft tumor phenotypes has been also unraveled by transcriptome analysis of the genome-wide gene expression in *Nrf1α^−/−^, Nrf1α^−/−^+siNrf2, Nrf2^−/−ΔTA^* and *caNrf2^ΔN^* cell lines, by comparison with wild-type *Nrf1/2^+/+^*. Notably, an overwhelming majority of the PTEN-directed PI3K-AKT signaling cascades are strikingly activated in *Nrf1α^−/−^*, but rather repressed in *Nrf2^−/−ΔTA^* cells. Silencing Nrf2 leads to opposing expression of 87 genes in between *Nrf1α^−/−^* and *Nrf1α^−/−^+siNrf2* cell lines. Although most cognate genes are, to different extents, co-regulated by Nrf1α and Nrf2, this study has highlighted about 30 of Nrf1α-specific downstream genes, and 38 of Nrf2-specific downstream genes. Among Nrf1α-regulated genes, those encoding A2M, EPHA8, FBXO2, KCND1, SLC2A3, SORL1, OLIG2, and RAPGEF4 may be responsible for the nervous system, albeit it is unclear whether they are relevant to those phenotypes of *Nrf1α^−/−^*-leading neurodegenerative diseases as reported by (Kobayashi et al., 2011; Lee et al., 2011). Only *ACSS2, FA2H,* and *KLF15* genes are associated with lipid metabolism, but it is required to determine their roles in relevant phenotypes, as described by (Bartelt et al., 2018; Hirotsu et al., 2014; Hou et al., 2018; Xu et al., 2005). By contrast, a portion of Nrf2-specific genes are critical for the development of various tissues and organs, neurons and cardiomyocytes, but none of the specific physio-pathological phenotypes in the *Nrf2^−/−ΔDBD^* mice are observed, implying their functions can be compensated. As such, the other Nrf2-specific genes may be involved in the development, movement and adhesion of epithelial cells, but it is unknown whether these gene functions enable Nrf2 to be endowed with its potent tumor-promoting roles in cancer progression and metastasis. All together, the malfunction of Nrf2, as a tumor promoter, is predominantly suppressed by Nrf1α that acts as a dominant tumor repressor. This pathophysiological process is tightly governed by endogenous regulatory networks. On the inside, there exist mutual opposing and unifying cross-talks between Nrf1α and Nrf2 at distinct levels. Concordantly, Nrf2 directly mediates transcription of the *Nrf1* gene to form a coupled positive and negative feedback circuit, in order to monitor Nrf1 and Nrf2 functioning towards precision expression of distinct downstream genes.

## STAR* METHODS

Detailed methods are provided in the online version of this paper and include the following:

- **KEY RESOURCES TABLE**
- **CONTACT FOR REAGENT AND RESOURCE SHARING**
- **EXPERIMENTAL MODEL AND SUBJECT DETAILS**
  - **Animal care and use**
  - **mouse embryonic fibroblasts (MEFs)**
  - **Human liver cell lines with distinct genotypes**
- **METHODS DETAILS**
  - **Cell culture and transfection**
  - **Expression constructs and other oligos used for siRNA and MirRNA**
  - **Subcutaneous tumor xenografts in nude mice**
  - **Histology**
  - **Lipid staining**
  - **Cellular ROS staining**
  - **Luciferase reporter assay**
  - **Real-time quantitative PCR**
  - **Western blotting**
  - **Flow cytometry analysis of cell cycle and apoptosis**
  - **The genome-wide transcriptiomic analysis**
- **QUANTIFICATION AND STATISTICAL ANALYSIS**
  - **Study approval**

## SUPPLEMENTAL INFORMATION

Supplemental Information includes ten figures and one table.

## AUTHOR CONTRIBUTIONS

L.Q. designed and performed most of the experiments except indicated elsewhere, made all figures and wrote the manuscript draft. M.W. performed the statistical analysis of transcriptome data. Y.R. participated in the preparation of gene knockout cell lines. X.R. and S.H. participated in animal experiments. S.W. provided critical suggestion and invaluable materials to improve the work. Y.Z. designed and supervised this study, interpreted all data, generated project resources, and wrote the manuscript. All authors reviewed and commented on the manuscript. Meanwhile, these authors declare no competing financial and other interests.

## ACKNOWLEDGMENTS

We are grateful to Dr. Akira Kobayashi (at Doshisha University, Japan) for providing both *Nrf1^−/−(ΔDBD)^* and *Nrf1^+/+^* MEF lines as a gift. The work was supported by the National Natural Science Foundation of China (81872336, 91129703, 91429305, and 31270879) awarded to Prof. Yiguo Zhang (University of Chongqing, China), and also funded in part by Chongqing University postgraduates′ innovation project (No. CYB15024) funded to Mr. Lu Qiu.

## STAR+METHODS

### KEY RESOURCES TABLE

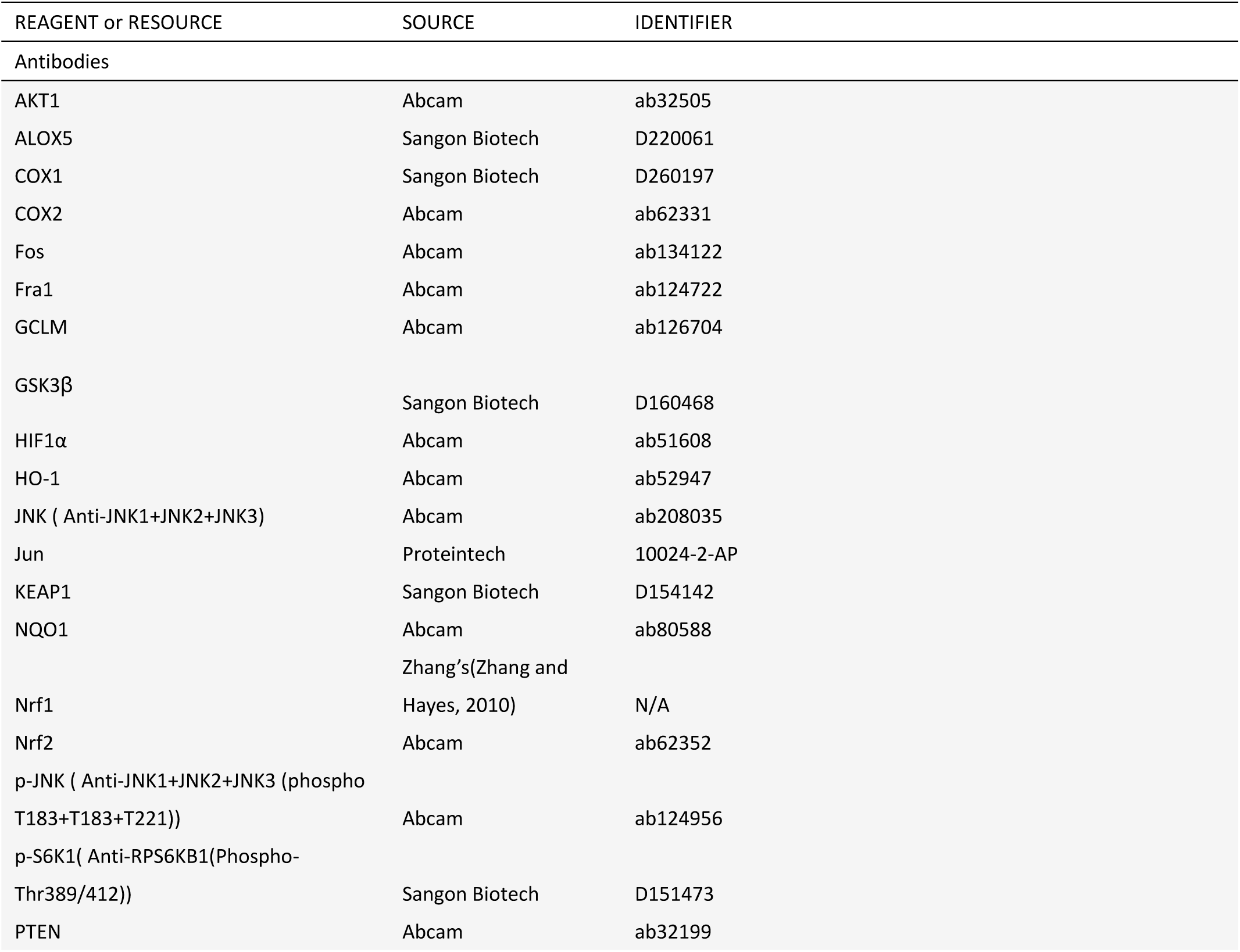
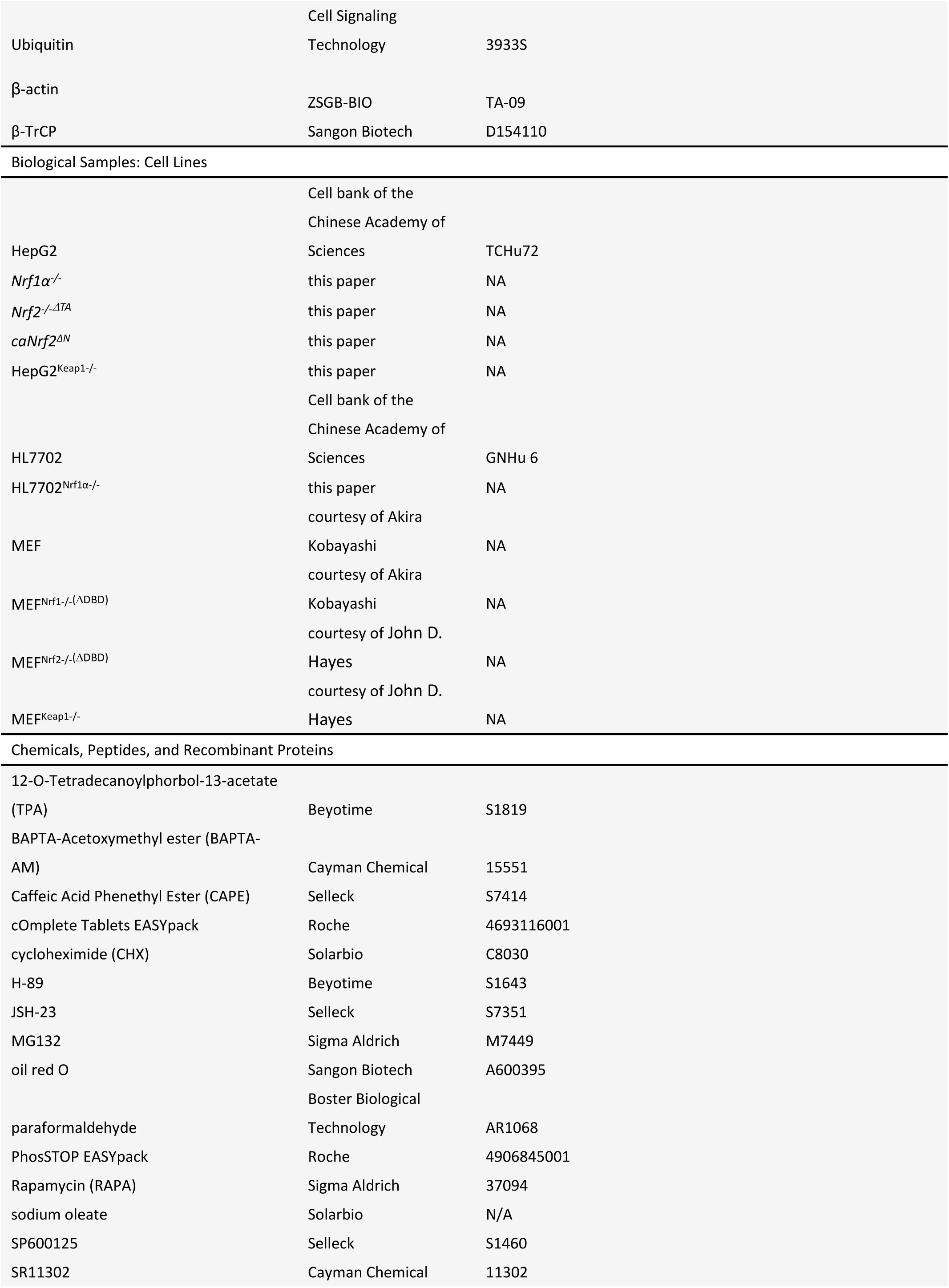
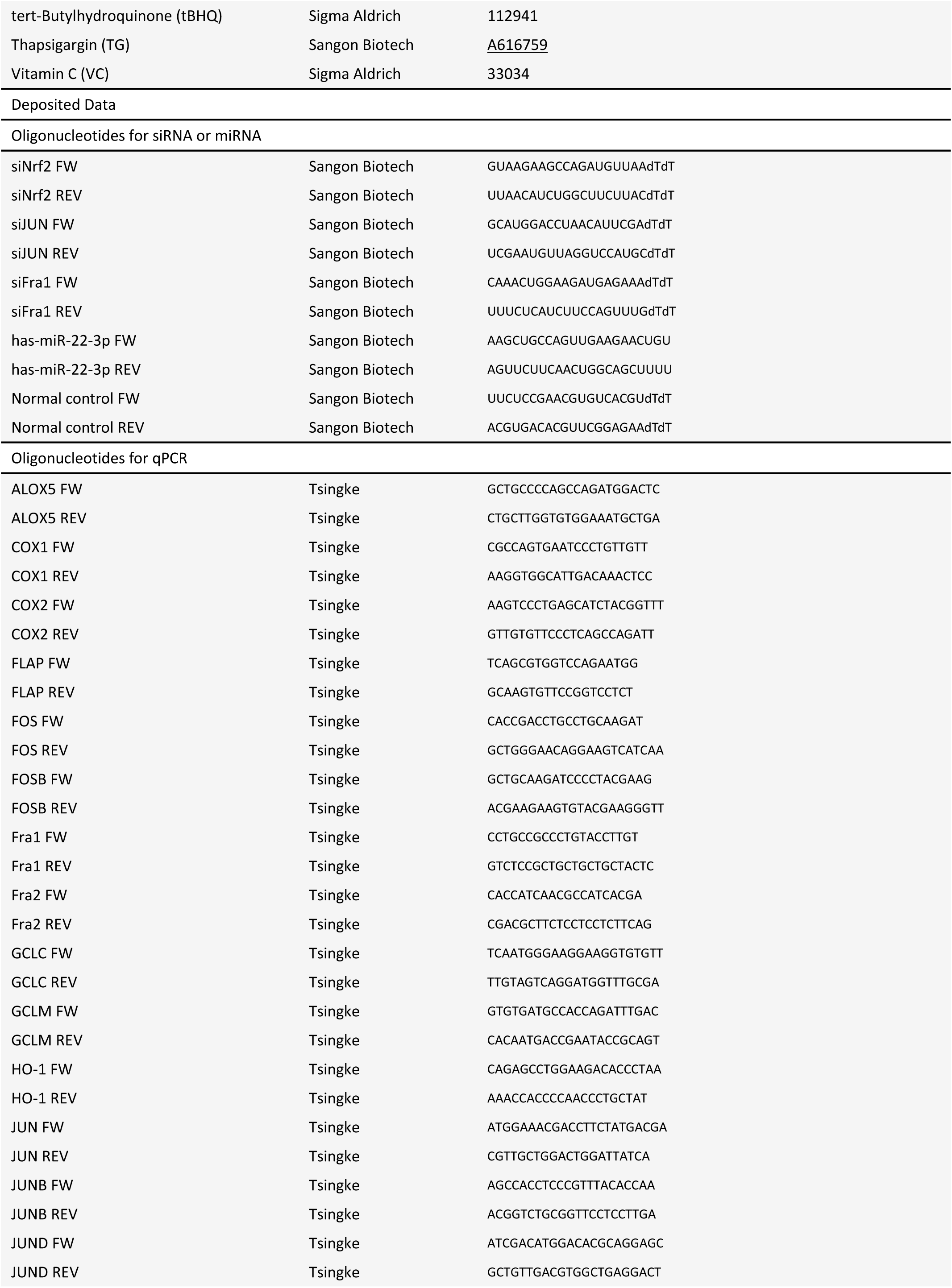
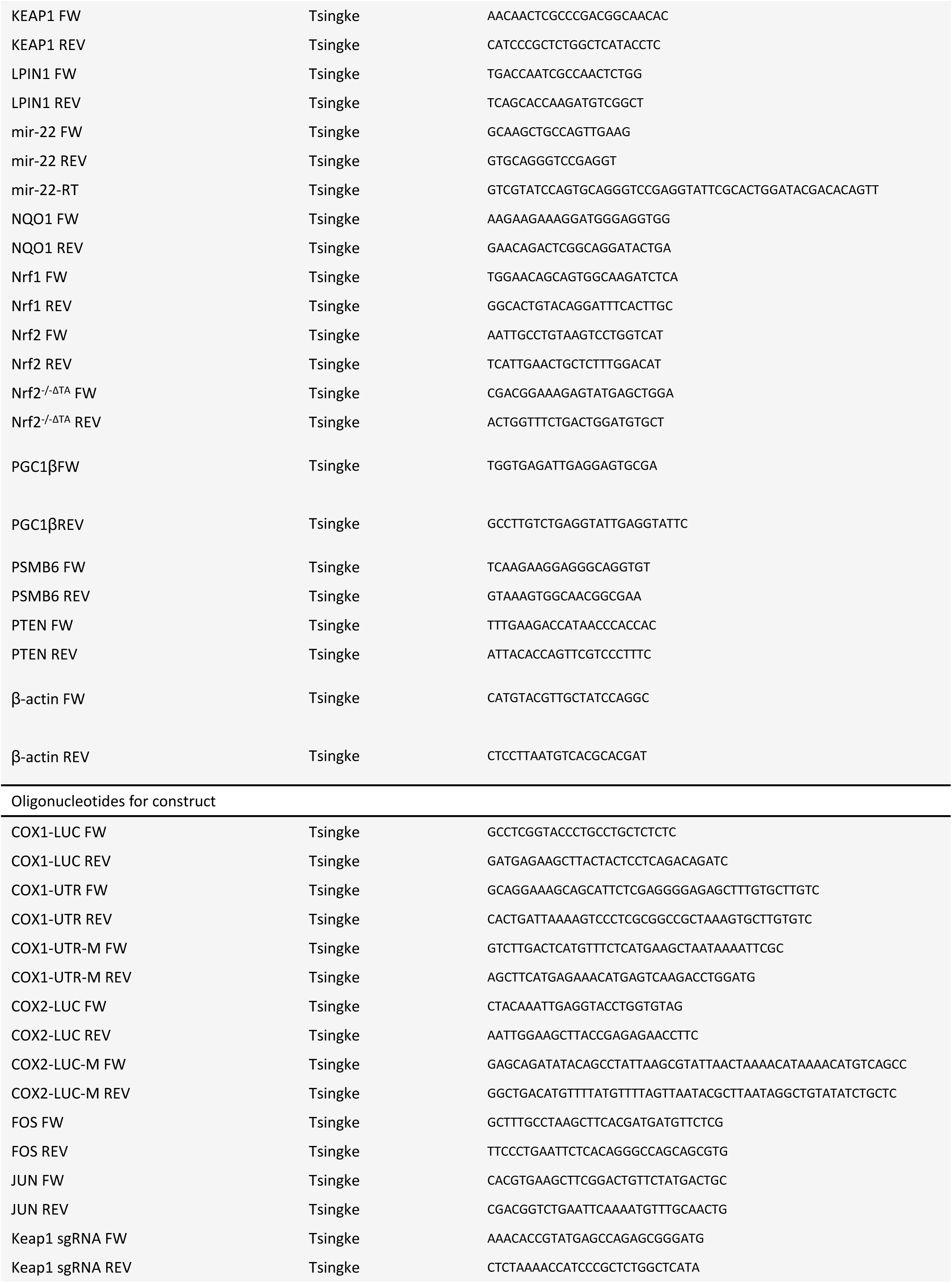
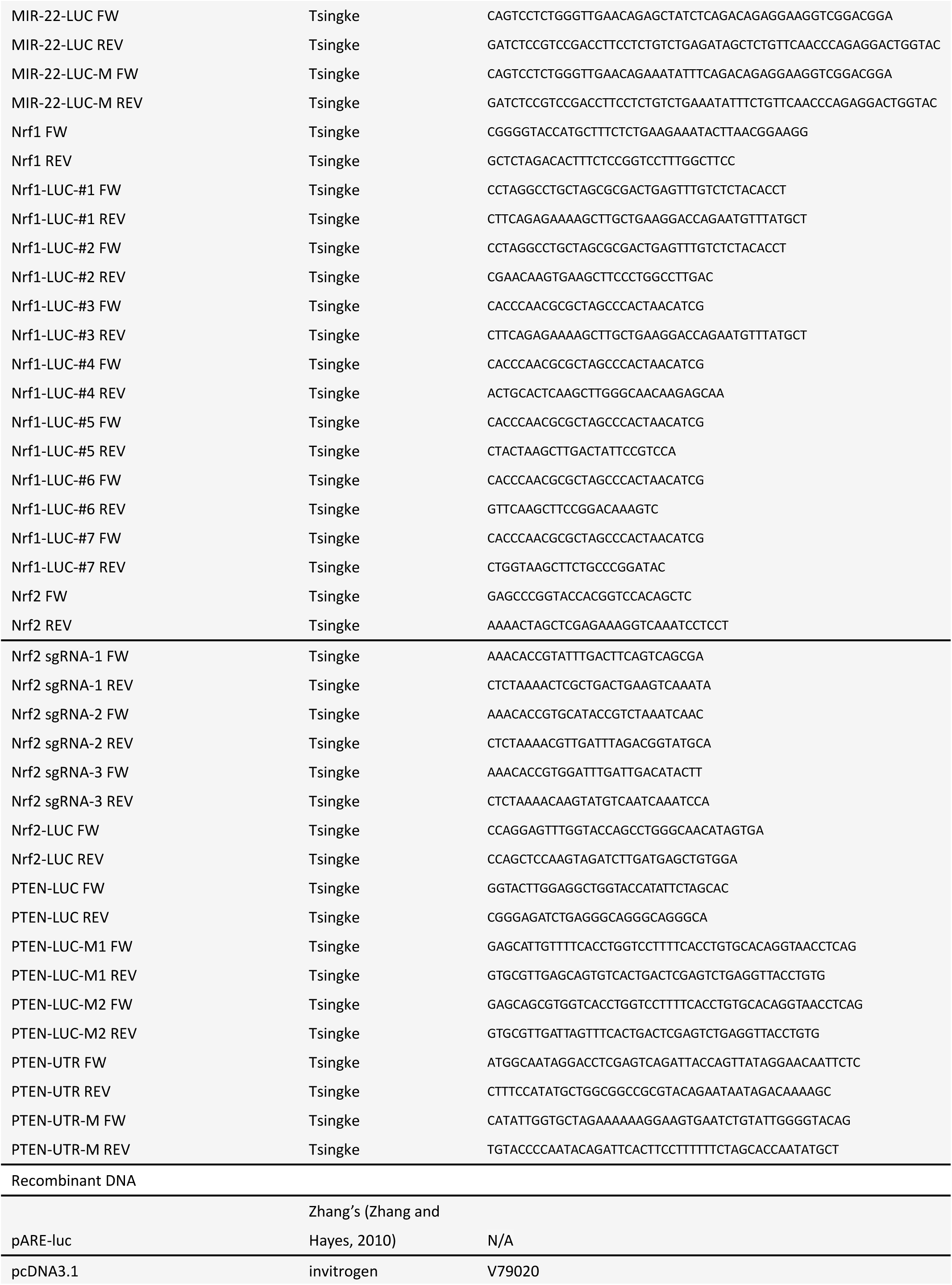
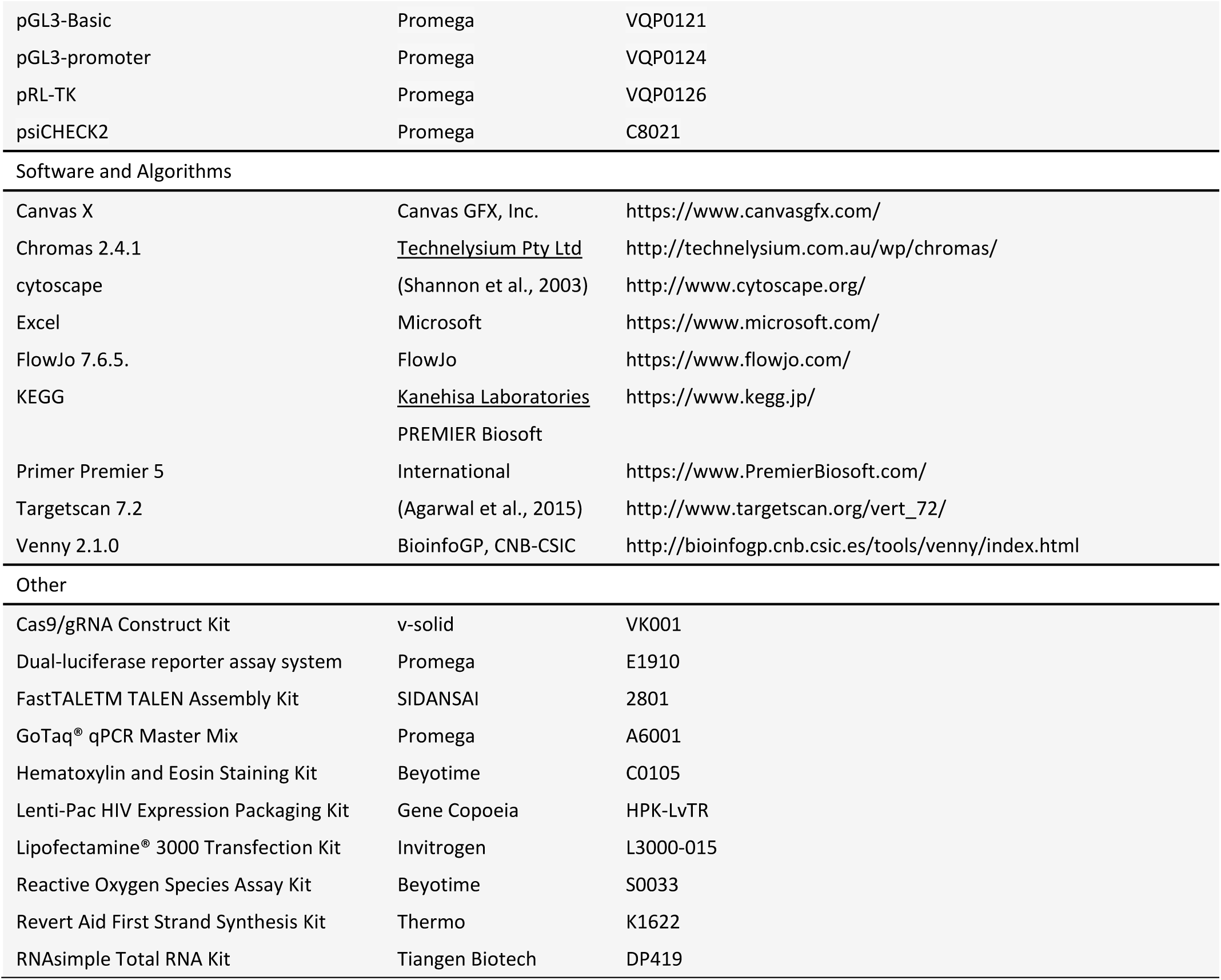

## CONTACT FOR REAGENT AND RESOURCE SHARING

Further information and requests for resources and reagents should be directed to and will be fulfilled by the Lead Contact, Yiguo Zhang

## EXPERIMENTAL MODEL AND SUBJECT DETAILS

### Animal care and use

In this study, the relevant animal experiments were indeed conducted according to the valid ethical regulations that have been approved. All mice were maintained under standard animal housing conditions with a 12-h dark cycle and allowed access *ad libitum* to sterilized water and diet. All relevant studies were carried out on 8-week-old male mice (with the license No. PIL60/13167) in accordance with the United Kingdom Animal (Scientific Procedures) Act (1986) and the guidelines of the Animal Care and Use Committees of Chongqing University and the Third Military Medical University, both of which were subjected to the local ethical review (in China). All relevant experimental protocols were approved by the University Laboratory Animal Welfare and Ethics Committee (with two institutional licenses SCXK-PLA-20120011 and SYXK-PLA-20120031).

### Mouse embryonic fibroblasts (MEFs)

The wild-type MEF and MEF^Nrf1−/−(ΔDBD)^ cell lines (in which ΔDBD represents deletion of mouse Nrf1 DNA-binding domain (DBD)-adjoining codon sequence) were provided as a gift by Dr. Akira Kobayashi (Doshisha University, Japan). Additional wild-type MEF and MEF^Nrf2−/−(ΔDBD)^, and MEF^Keap1−/−^ were from Prof. John D. Hayes (University of Dundee, UK). In fact, it should also be credited to Prof. Masayuki Yamamoto (Tohoku University, Japan) because all relevant mouse lines were originally made in his laboratory.

### Human liver cell lines with distinct genotypes

All four cell lines *Nrf1α^−/−^, Nrf1α^−/−^*+siNrf2, *Nrf2^−/−ΔTA^* and *caNrf2^ΔN^* were created in this study. Their progenitor cells are the human hepatocellular carcinoma (HepG2) and/or another non-cancerous human liver (HL7702) cell lines. The latter two lines HepG2 and HL7702 are wild-type (*Nrf1/2^+/+^, Keap1^+/+^*) cells, because not any mutants in the *Nrf1, Nrf2* and *Keap1* genes are therein confirmed by sequencing. However, it is important to be noted that Nrf1 and its long TCF11 isoform are co-expressed at a ratio of 1:1 in HL7702 cells. By contrast, significantly decreased expression of Nrf1 is observed in HepG2 cells (Ren et al., 2016), while almost none of its longer TCF11 transcripts were detected. For relevant identification of these cell lines, see Figures 1 and S1.

## METHODS DETAILS

### Cell culture and transfection

Cells were grown in DMEM supplemented with 5 mM glutamine, 10% (v/v) foetal bovine serum (FBS), 100 units/ml of either of penicillin and streptomycin, in the 37 °C incubator with 5% CO_2_. The experimental cells were transfected with indicated plasmids alone or in combination for 8 h using Lipofectamine^®^ 3000 Transfection Kit (Invitrogen, USA), and then allowed for recovery from transfection in the fresh medium for 24 h before being subjected to indicated experiments.

### Expression constructs and other oligos used for siRNA and miRNA

Expression constructs for human Nrf1, Nrf2, JUN and FOS were made by cloning each of their full-length cDNA sequences into a pcDNA3 vector, respectively. Notably, the other plasmids specifically for the genome-editing of Nrf1 or Nrf2 by Talens or CRISPR/Cas9 were created (in Figure S1) and identified (in Figure 1). Furtherly, we were made four specific luciferase reporters, which were driven by distinct gene promoter regions from the human *Nrf1, Nrf2, COX1* and *COX2*. Different lengths of these gene promoter regions were amplified by PCR from their genomic loci and inserted into the PGL3-basic vector. In addition to these intact reporter genes *P_Nrf1_-Luc, P_Nrf2_-Luc, P_COX1_-Luc, P_COX2_-Luc* and *miR22-ARE-Luc*, all these relevant ARE-specific mutant reporters were engineered. Moreover, double fluorescent reporters (i.e. *PTEN-miR22b* and *COX1-miR22b*) were created by cloning the 3’ UTR region sequences of *COX1* and *PTEN*, that were amplified from reverse transcription PCR products and ligated into the psiCHECK2 vector. All primers and other oligos used for siRNAs and Mir-RNAs were synthesized by Sangon Biotech (Shanghai, China). The fidelity of all constructs used in this study was confirmed to be true by sequencing.

### Subcutaneous tumor xenografts in nude mice

Mouse xenograft models were made by subcutaneous heterotransplantation of the human hepatoma HepG2 (i.e. *Nrf1/2^+/+^* or each derivate of *Nrf1α^−/−^, Nrf1α^−/−^*+siNrf2, *Nrf2^−/−ΔTA^* and *caNrf2^ΔN^* cell lines into nude mice, as described (Morton and Houghton, 2007). Experimental human hepatoma cells (1 × 10^7^, allowed for growth in the exponential phase) were suspended in 0.2 ml of serum-free DMEM and were inoculated subcutaneously into the right upper back region of male nude mice (BALB/C*nu/nu*, 6 weeks, 18 g, from HFK Bioscience, Beijing) at a single site. The procedure of injection into all mice was completed within 30 min, and the formation of the subcutaneous tumour exnografts was observed. Once the tumor exnografts emerged, their sizes were successively measured once every two days, until the 32^nd^ day when the mice were sacrificed before the transplanted tumors were excised. The sizes of growing tumors were calculated by a standard formula (i.e. V = ab^2^/2) and then are shown graphically (n = 6 per group). Subsequently, the tumor tissues were also subjected to the histopathological examination by the routine hematoxylin-eosin staining. Notably, all the relevant animal experiments in this study were indeed conducted according to the valid ethical regulations that have been approved.

### Histology

Xenograft tumor tissues were immersed in 4% paraformaldehyde for overnight and then transferred to 70% ethanol. Individual tumor tissues were placed in processing cassettes, dehydrated through a serial of alcohol gradient, and embedded in paraffin wax blocks. Before staining, the tissue sections were de-waxed in xylene, rehydrated through decreasing concentrations of ethanol, and washed in PBS. Lastly, they were stained with hematoxylin and eosin (H&E), and visualized by microscopy.

### Lipid staining

Experimental cells were seeded in 6-well plates and cultured with 200 µM sodium oleate (Solarbio, Beijing, China) medium. The cells were fixed for 30 min with 4% paraformaldehyde (AR1068, Boster Biological Technology, Wuhan, China) and then stained for 30 min with a solution of 3 g/L oil red O (A600395, Sangon Biotech, Shanghai, China). The stained cells were rinsed 3 times with 60 % of isopropyl alcohol (Kelong, Chengdu, China) before the red lipid droplets were visualized by microscopy.

### Cellular ROS staining

Experimental cells were allowed for growth to an appropriate confluence in 6-well plates and then incubated in serum-free medium containing 10 µM of 2’,7’-Dichlorodihydrofluorescein diacetate (DCFH-DA) (S0033, Beyotime, Shanghai, China) at 37°C for 20 min. Thereafter, the cells were washed three times with serum-free medium, before the green fluorescent images were achieved by microscopy.

### Luciferase reporter assay

Equal numbers (1.0×10^5^) of experimental cells were seeded into each well of the 12-well plates. After reaching 80% confluence, the cells were transfected with a Lipofectamine 3000 mixture with luciferase plasmids with or without other expression plasmids. In the pGL3 plasmid system, the Renilla-luciferase, which expression by pRL-TK plasmid, as an internal control for transfection efficiency. And in the psi-CHECK2 plasmid system, the Pyralis-luciferase activity is the internal control, while the Renilla-luciferase activity is the experimental test object. The luciferase activity was measured by using the dual-luciferase reporter assay system (E1910, Promega). The resultant data were normalized as a fold change (mean ± S.D) relative to the activity of the control group (at a given value of 1.0). The data presented here represent at least 3 independent experiments undertaken on separate occasions that were each performed in triplicate. Significant differences in the transcriptional activity were subjected to statistical analysis.

### Real-time quantitative PCR

Experimental cells were subjected to isolation of total RNAs by using the RNAsimple Kit (Tiangen Biotech CO., Beijing). Subsequently, 500 ng of total RNAs were added in a reverse-transcriptase reaction to generate the first strand of cDNA (with Revert Aid First Strand Synthesis Kit from Thermo). The synthesized cDNA was served as the template for qPCR, in the GoTaq^®^ qPCR Master Mix (from Promega), before being deactivated at 95°C for 10 min, and amplified by 40 reaction cycles of the annealing at 95°C for 15 s and then extending at 60°C for 30 s. The final melting curve was validated to examine the amplification quality, whereas the mRNA expression level of β-actin served as an optimal internal standard control.

### Western blotting

Experimental cells were harvested in a lysis buffer (0.5% SDS, 0.04 mol/L DTT, pH 7.5), which was supplemented with the protease inhibitor cOmplete Tablets EASYpack or phosphatase inhibitor PhosSTOP EASYpack (either 1 tablet per 10 mL, Roche, Germany). The lysates were denatured immediately at 100°C for 10 min, sonicated sufficiently, and diluted in 3× loading buffer (187.5 mmol/L Tris-HCl, pH 6.8, 6% SDS, 30% Glycerol, 150 mmol/L DTT, 0.3% Bromphenol Blue) at 100°C for 5 min. Subsequently, equal amounts of protein extracts were subjected to separation of proteins by SDS-PAGE containing 4-15% polyacrylamide, and then visualization by Western blotting with distinct antibodies as indicated. On some occasions, the blotted membranes were stripped for 30 min and then re-probed with additional primary antibodies. β-actin served as an internal control to verify equal loading of proteins in each of electrophoretic wells.

### Flow cytometry analysis of cell cycle and apoptosis

Experimental cells (5 × 10^5^) were allowed for growth in 60-mm cell culture plate for 48 h and synchronization by 12-h starvation in a serum-free medium, before being treated with 10 μmol/L BrdU for 12 h. The cells were fixed for 15 min with 100 μl of BD Cytofix/Cytoperm buffer (containing a mixture of the fixative paraformaldehyde and the detergent saponin) at room temperature and permeabilized for 10 min with 100 μl of BD Cytoperm permeabilization buffer plus (containing fetal bovine serum as a staining enhancer) on ice. Thereafter, the cells were re-fixed and treated with 100 μl of DNase (at a dose of 300 μg/ml in PBS) for 1 h at 37 °C, in order to expose the incorporated BrdU, followed by staining with FITC conjugated anti-BrdU antibody for 60 min at room temperature. Subsequently, the cells were suspended in 20 μl of 7-amino-actinomycin D solution 20 min for the DNA staining and re-suspended in 0.5 ml of a staining buffer (i.e. 1 × DPBS containing 0.09% sodium azide and 3% heat-inactivated FBS), prior to the cell cycle analysis by flow cytometry. Furthermore, additional fractions of cells (5 × 10^5^) were allowed for 48 h growth in 60-mm cell culture plate before being used for apoptosis analysis. The cells were pelleted by centrifuging at 1000 × *g* for 5 min and washed by PBS for three times, before being incubated for 15 min with 5 μl of Annexin V-FITC and 10 μl of propidium iodide (PI) in 195 μl of the binding buffer, prior to flow cytometry analysis of cell apoptosis. The results being analyzed by the FlowJo 7.6.1 software.

### The genome-wide transcriptomic analysis

Total RNAs were subjected to the sequencing by the Beijing Genomics Institute (BGI, www.genomics.org.cn) on the platform of BGISEQ-500 (contract No. is F17FTSCCWLJ1161). After removing the ‘dirty’ raw reads with data filtering, the clean reads were generated and mapped to the reference by using both HISAT (Kim et al., 2015) and Bowtie2 (Langmead et al., 2009) tools. Of note, gene expression levels were calculated by using the FPKM (Fragments Per Kilobase of exon model per Million mapped fragments) method with RSEM (Li and Dewey, 2011). Then, differentially expressed genes (DEGs) were identified with the criteria Fold-change ≥2 and diverge probability ≥0.8 by using the NOISeq (Tarazona et al., 2011). For the functional annotation, all DEGs were mapped to gene ontology (GO) terms in the database (http://www.geneontology.org/) and the pathway enrichment analysis of DEGs was performed by using KEGG (Kanehisa et al., 2008).

## QUANTIFICATION AND STATISTICAL ANALYSIS

Significant differences were statistically determined using the Student’s *t*-test and Multiple Analysis of Variations (MANOVA). The data are here shown as a fold change (mean ± S.D.), each of which represents at least 3 independent experiments that were each performed in triplicate.

### Study approval

All experimental procedures were approved by the Animal Care and Use Committees of Chongqing University and the Third Military Medical University, both of which were subjected to the local ethical review (in China). Randomly assigned nude mice were injected subcutaneously with distinct cell lines as experimented.

---

**Figure S1.**
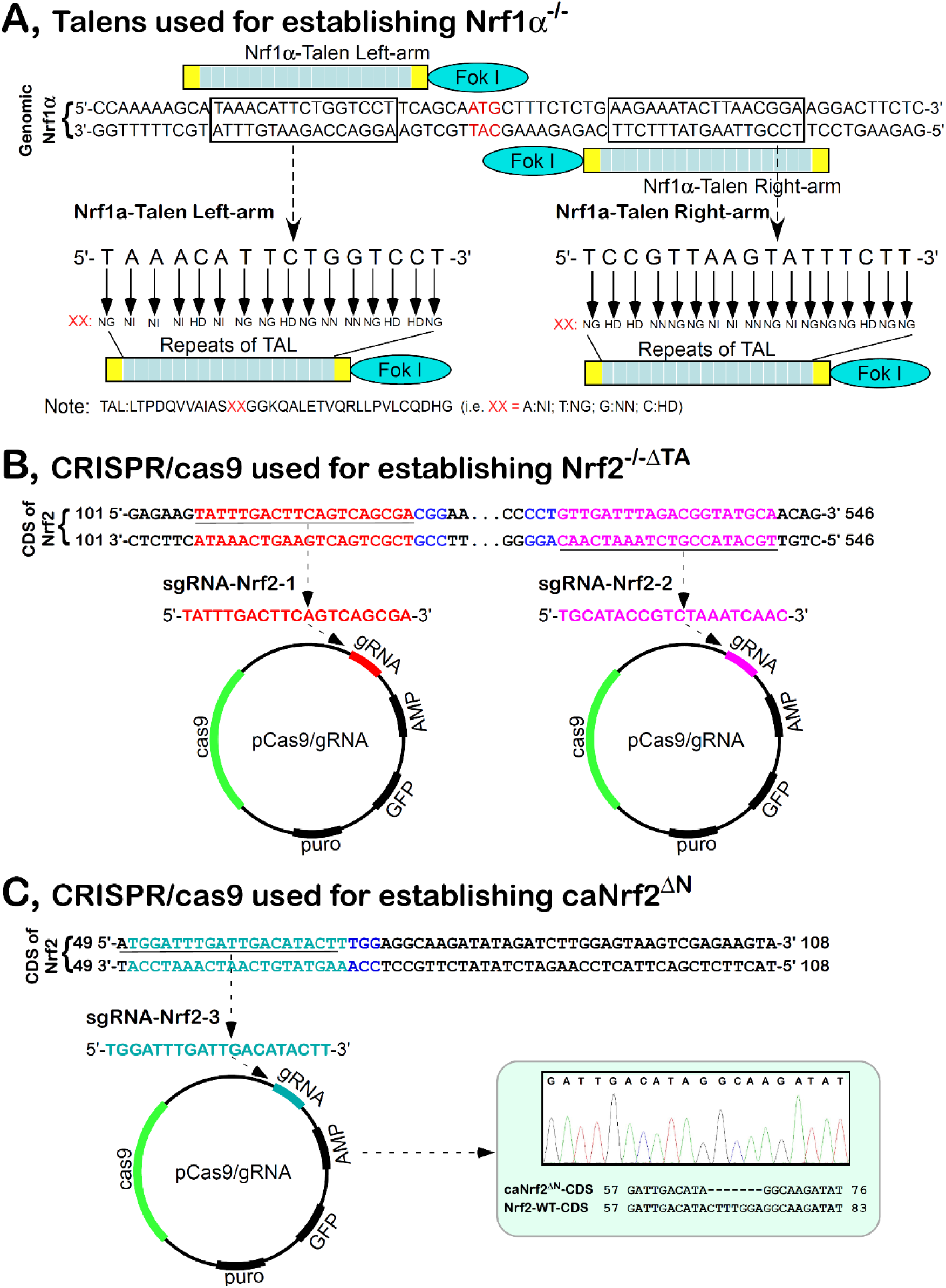
The human *Nrf1-* and *Nrf2-specific* gene-editing constructs. (A) Nrf1α-specific targeting constructs for TALEN-mediated gene editing. Both its left-and right-arms were designed for deletion of the first translation initiation codons of the *Nrf1* gene (i.e. *Nrf1α^−/−^*). (B) Nrf2-specific constructs for CRISPR/CAS9-directed gene editing. They were designed for deleting a fragment of the *Nrf2* gene encoding most of both Neh4 and Neh5 domains (to yield an inactive *Nrf2^−/−ΔTA^*). (C) Another Nrf2-specific editing construct by CRISPR/CAS9. It was designed for the dominant-active mutant of Nrf2, so as to delete the sequence encoding the N-terminal Keap1-binding domain. The resulting mutant (i.e. *caNrf2^ΔN^*) was aligned with wild-type nucleotide sequence of *Nrf2*.

**Figure S2.**
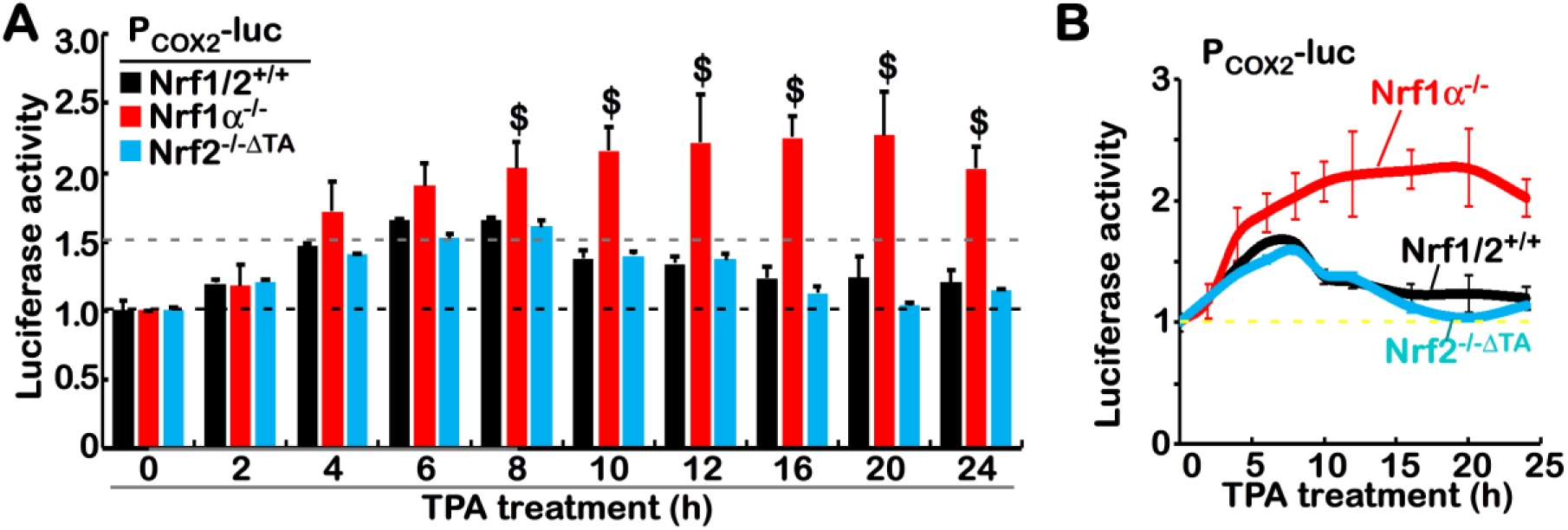
Distinct cellular responses of the *P_COX2_-luc* reporter gene to TPA. (A) *Nrf1/2^+/+^, Nrf1α^−/−^* and *Nrf2^−/−ΔTA^* cells were transfected with the *P_COX2_-luc* and *pRL-TK* reporters for 12 h, and then treated with100 nM of TPA for indicated lengths of time, before being measured for the luciferase activity. The data are shown as mean ± SEM (n = 3×3; $, *p*< 0.01 compared with the untreated control values). (B) The above data are shown graphically.

**Figure S3.**
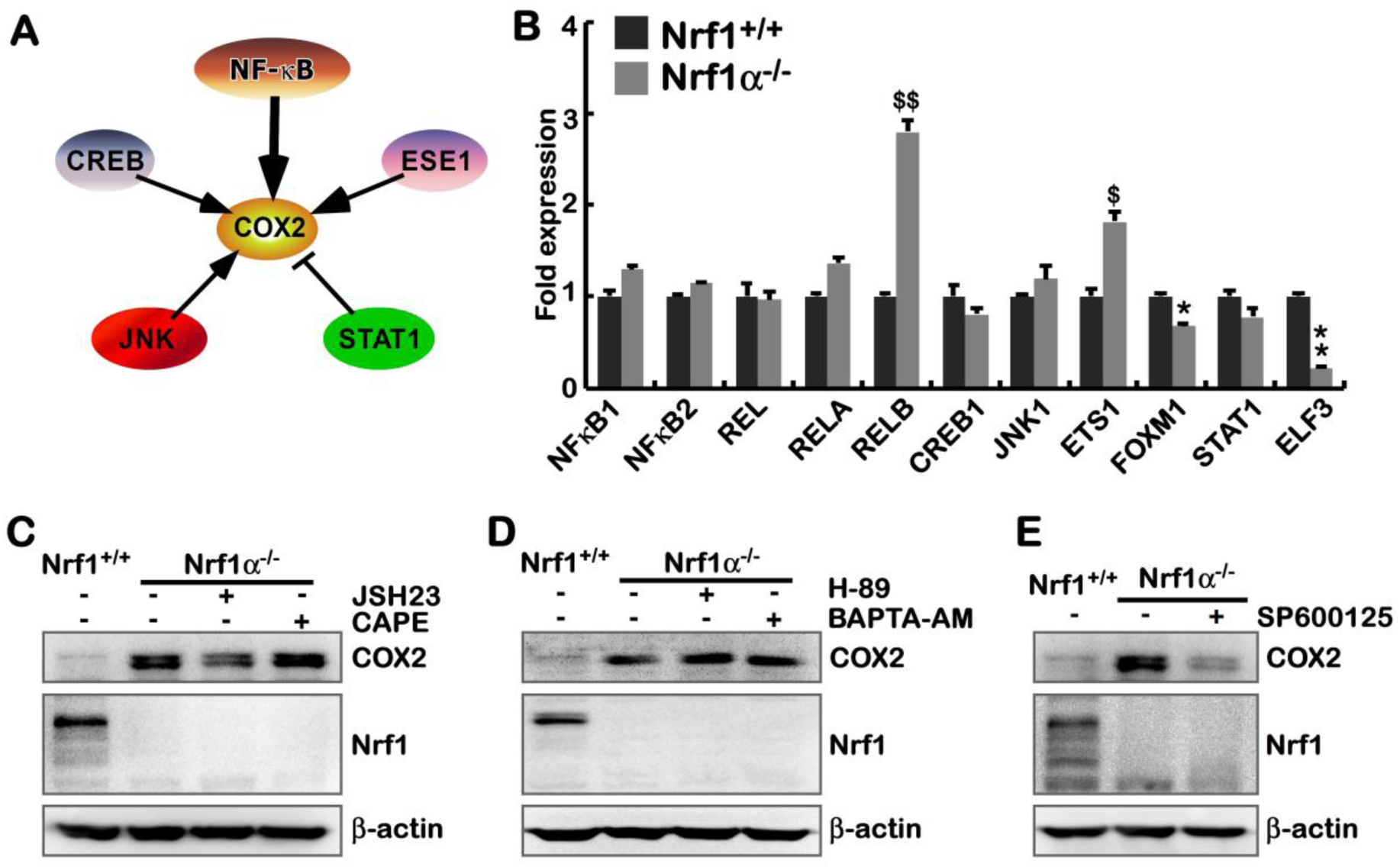
The JNK inhibitor blocks the *Nrf1α^−/−^*-leading increase of COX2. (A) Schematic representation of potential upstream signaling to regulate COX2. (B) Alterations in the indicated gene expression in *Nrf1α^−/−^*, compared with *Nrf1/2^+/+^*, cells. The data were obtained from transcriptome and are shown as mean ± SEM (n=3; \**p*< 0.01; \*\**p*< 0.001; $, *p*< 0.01; $$, *p*< 0.001).

(C to E) *Nrf1α^−/−^* cells were treated for 24 h with (*C*) 20 μM of JSH23, 25μM of CAP, (*D*) 10 μM of H-89, 1 μM of BAPTA-AM, or (*E*) 20 μM of SP600125, before COX2 was examined by Western blotting.

**Figure S4.**
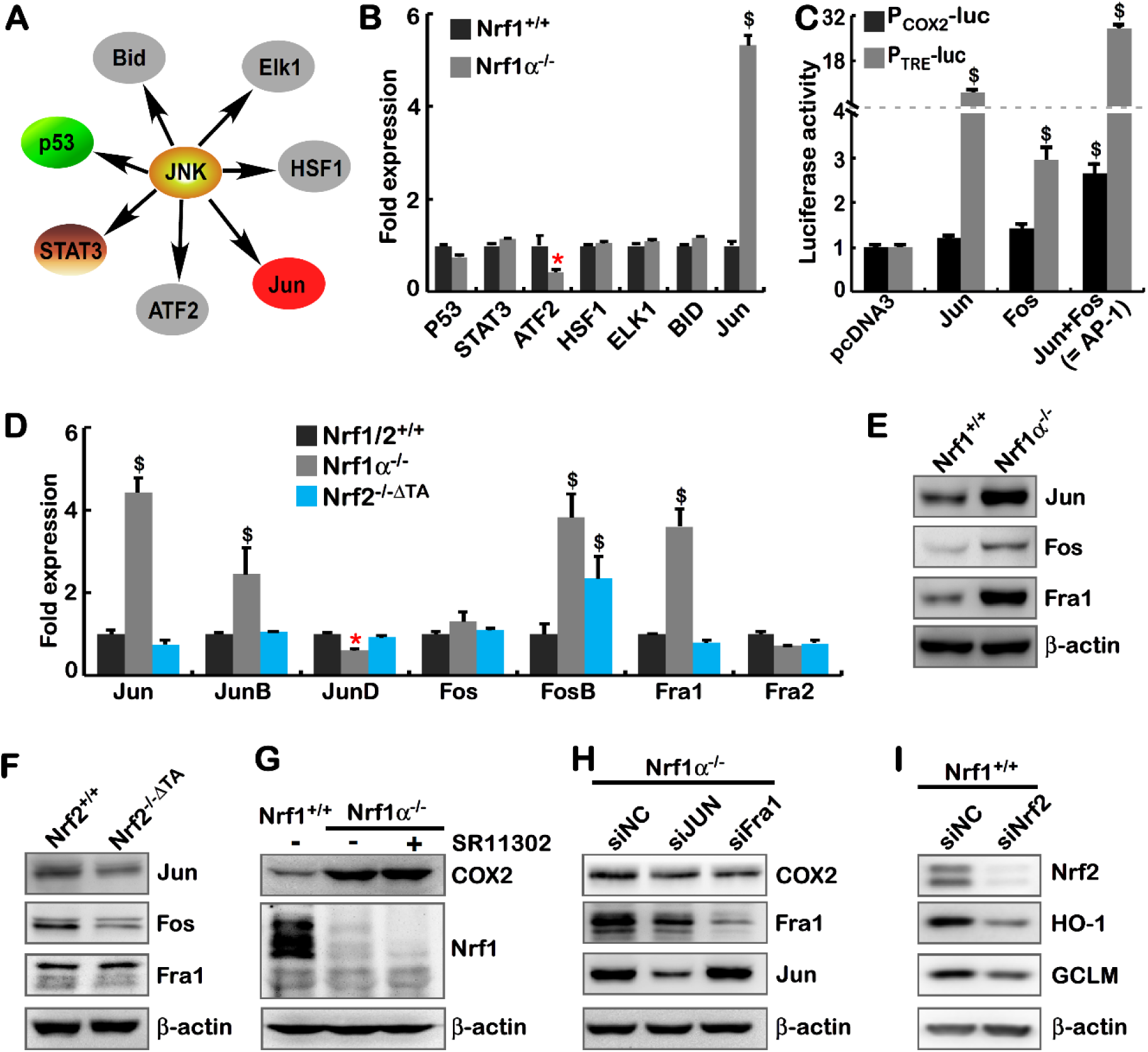
Activation of some AP-1 components in *Nrf1α^−/−^* cells. (A) The cartoon shows possible JNK signaling to downstream targets. (B) The transcriptome analysis of major downstream genes regulated by JNK signaling. The data are shown as mean ± SEM (n=3, \**p*< 0.01; $$, *p*< 0.001 compared with wild-type values) (C) Either *P_COX2_-luc* or P_TRE_-luc together with *pRL-TK* were co-transfected with each of indicated expression constructs or empty pcDNA3 vector and allowed for 24-h recovery, before being determined. The data are shown as mean ± SEM (n = 3×3; $, *p*< 0.01; $$, *p*< 0.001). (D) The real-time qPCR analysis of distinct AP-1 subunits at their mRNA levels in *Nrf1/2^+/+^, Nrf1α^−/−^* and *Nrf2^−/−ΔTA^* cells. The data are shown as mean ± SEM (n= 3×3, \**p*< 0.01, $ *p*< 0.01; $$ *p*< 0.001). (E) Western blotting of JUN, FOS, and Fra1 abundances in *Nrf1α^−/−^* and *Nrf1/2^+/+^*, cells. (F) Abundances of JUN, FOS, and Fra1 was visualized Western blotting of *Nrf2^−/−ΔTA^* and *Nrf1/2^+/+^* cells. (G) *Nrf1α^−/−^* cells were treated with 4 μM of SR11302 for 24 h before COX2 were examined by Western blotting. (H) *Nrf1α^−/−^* cells were allowed for knockdown by siJUN (60 nM) and siFOSL1 (60 nM) for 24 h, respectively, before COX2, Fra1 and JUN were determined by Western blotting. (I) *Nrf1/2^+/+^* cells were subjected to silencing of siNrf2 (60 nM) and allowed for 24-h recovery, before Nrf2, HO1 and GCLM were visualized by immunoblotting.

**Figure S5.**
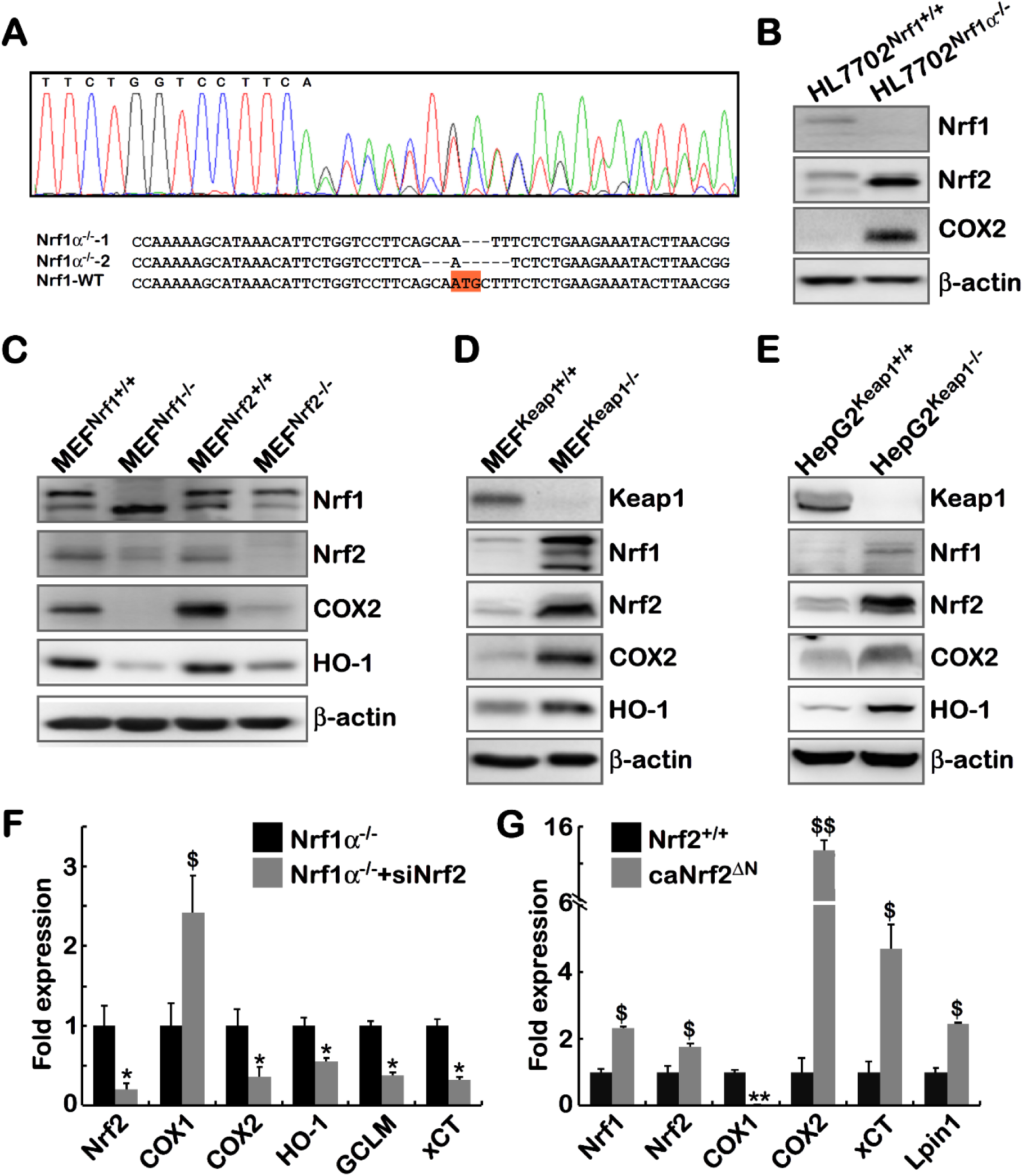
Cross-talks between Nrf1 and Nrf2 to regulate COX2. (A) Identification of HL7702^Nrf1α*−/−*^ by the genomic site-specific sequencing. The resulting mutant of *Nrf1α* was aligned with the wild-type nucleotide sequence. (B) Distinctions of Nrf1, Nrf2 and COX2 in between HL7702^Nrf1α*−/−*^ and HL7702^Nrf1^*+/+* cells was observed by Western blotting. (C) Subtle nuances in the abundances of Nrf1, Nrf2, COX2 and HO-1 in between MEF ^Nrf1 +/+^, MEF ^Nrf1−/−^, MEF ^Nrf2+/+^ and MEF ^Nrf2−/−^ were determined by Western blotting. (D) Alterations in the expression of Keap1, Nrf1, Nrf2, COX2 and HO-1 in between MEF ^Keap1+/+^ and MEF^Keap1−/−^ were detected by Western blotting. (E) Differences of Keap1, Nrf1, Nrf2, COX2, HO-1 abundances in between HepG2 ^Keap1+/+^ and HepG2^Keap1^*^−/−^* were visualized by Western blotting. (F) Differential expression of *Nrf2, COX1, COX2, HO-1, GCLM* and *xCT* at mRNA levels in *Nrf1α^−/−^* and *Nrf1α^−/−^*+siNrf2 cells were determined by the transcriptome. The data are shown as mean ± SEM (n=3, \**p*< 0.01; $ *p*< 0.01). (G) Both *Nrf2^−/−ΔTA^* and *caNrf2^ΔN^* cell lines differentially expressed mRNA levels of *Nrf1, Nrf2, COX1, COX2, xCT* and *Lpin1.* The transcriptome FPKM data are shown as mean ± SEM (n=3, \**p*< 0.01; $ *p*< 0.01)

**Figure S6.**
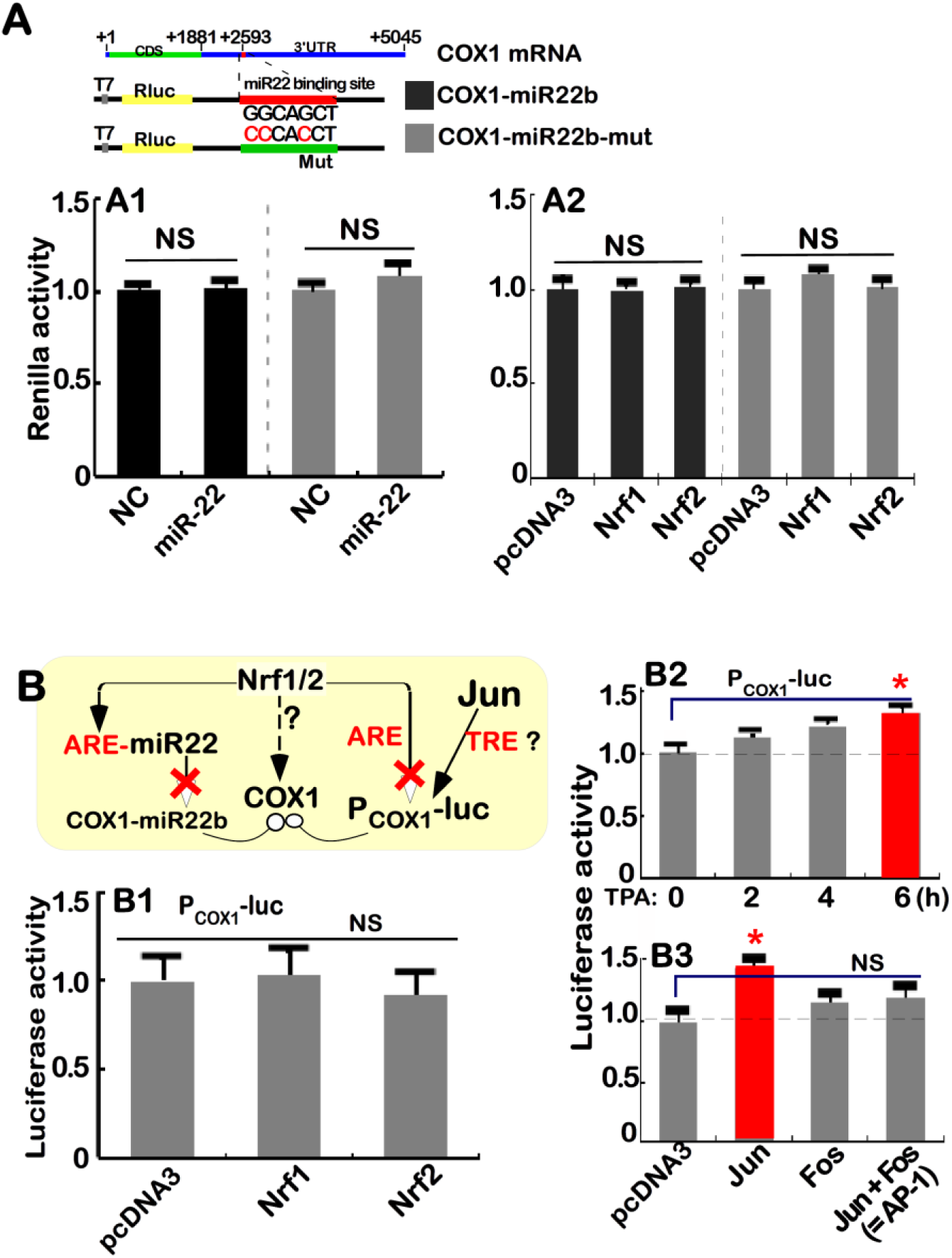
Genetic analysis of COX1 regulation. (A) The *COX1-miR22b* was constructed as above, which contains miR-22 binding site which in the COX1’s 3’UTR region(*upper*). *Nrf1/2^+/+^* cells were co-transfected with *COX1-miR22b* or *COX1-miR22b-mut*, together with miR-22 or NC plasmids (A1), or pcDNA3, an expression construct for Nrf1 or Nrf2 (A2), and then allowed for 24-h recovery before being determined. The data are shown as mean ± SEM (n = 3×3, NS= no statistical difference). (B) *Nrf1/2^+/+^* cells were co-transfected with the *P_COX1_-luc* and *pRL-TK* (***B1 to B3***), plus pcDNA3 or indicated expression constructs for Nrf1, Nrf2 (***B1***), Jun, Fos or Jun+pFos (***B3***), and allowed for 24-h recovery, before being treated (***B2***), or were not treated (***B1, B3***), with 100 nM of TPA for 2-6 h, prior to being measured for the luciferase activity. The data are shown as mean ± SEM (n = 3×3, \**p*< 0.01, NS= no statistical difference).

**Figure S7.**
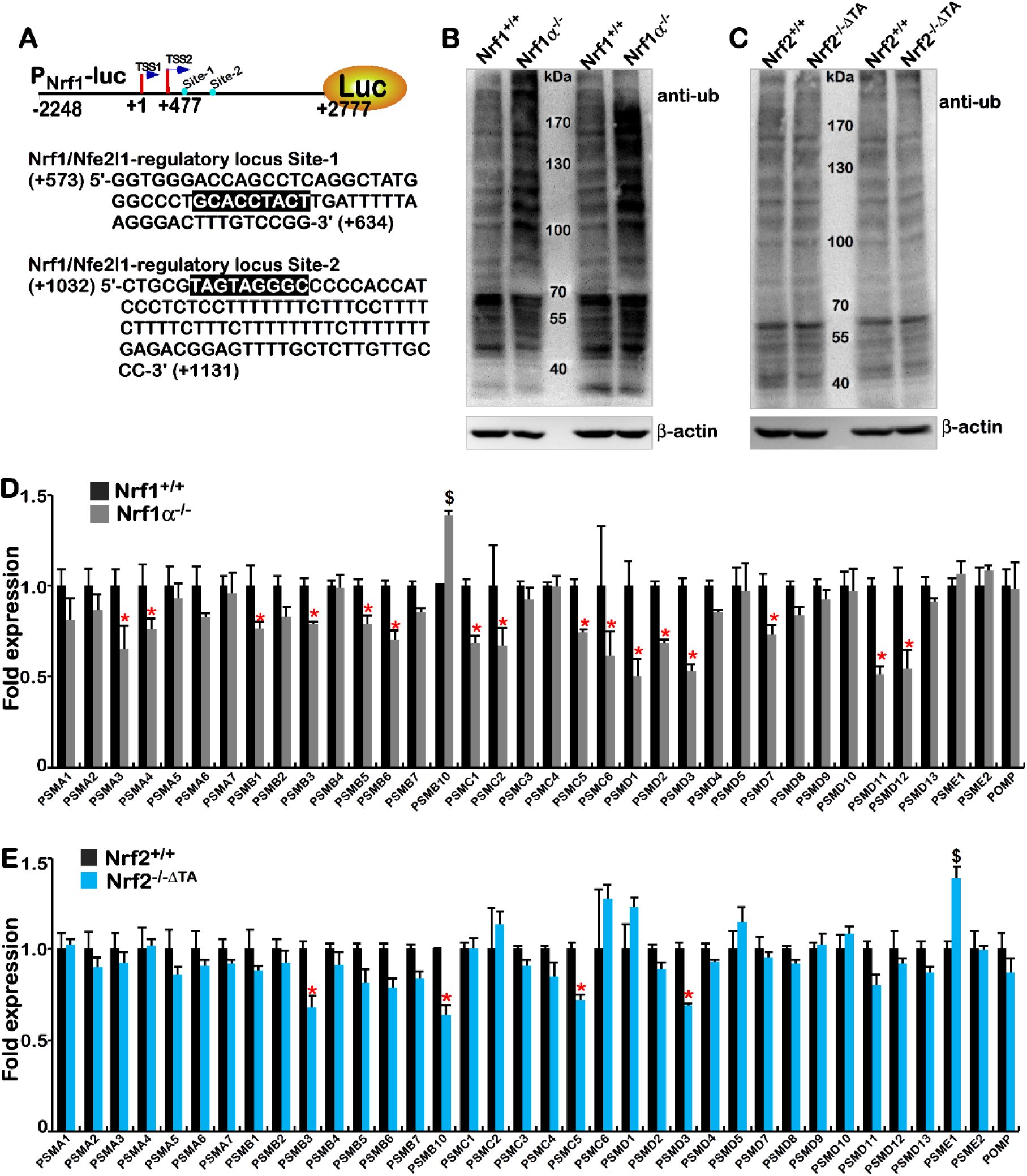
Differences in transcriptional expression of proteasomal subunits regulated by Nrf1 and Nrf2. (A) Two *cis*-*Nrf1/Nef2l1*-regulatory locus sites (i.e. Site-1 and Site-2) exist in this gene promoter, as located (*upper*). The nucleotide sequence of both Site-1 and Site-2 are shown. (B) Immunoblotting with antibodies against ubiquitinated proteins (i.e. anti-ub) in *Nrf1/2^+/+^* and *Nrf1α^−/−^* cells. (C) Almost no or less anti-ub cross-reactivity with ubiquitinated proteins in *Nrf1/2^+/+^* and *Nrf2^−/−ΔTA^* cells was observed. (D) Significant decreases in the expression of most of the 26S proteasomal subunits and related proteins were detected in *Nrf1α^−/−^* cells when compared with those in *Nrf1/2^+/+^*. The transcriptome data are shown as mean ± SEM (n=3, \**p*< 0.01; $ *p*< 0.01). (E) Almost no changes in transcriptional expression of most proteasomal and related genes in *Nrf1/2^+/+^* and *Nrf2^−/−ΔTA^*cells. The transcriptome data are shown as mean ± SEM (n=3, \**p*< 0.01; $ *p*< 0.01).

**Figure S8.**
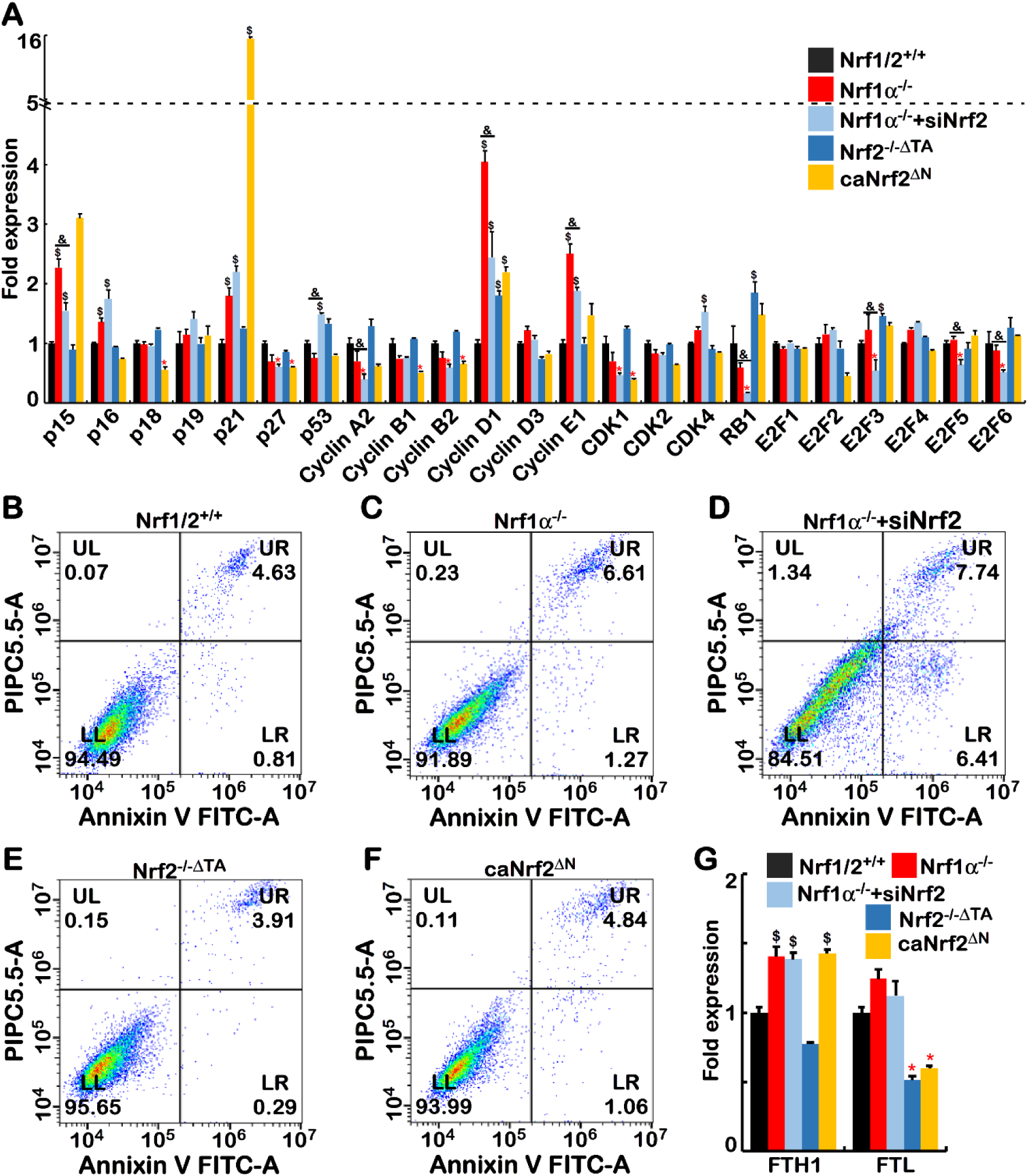
Subtle nuances in distinct cell cycles and apoptosis processes. (A) Changes in expression of cell cycle-related genes in five distinct cell lines as indicated. The transcriptome data are shown as mean ± SEM (n=3, \**p*< 0.01; $ *p*< 0.01). (B to F) Flow cytometry analysis of apoptosis in five distinct cell lines as indicated. Abbreviations: UL, necrotic cells; UR, early apoptotic cells; LL, normal cells; LR, late apoptotic cells. (G) The expression of *FTH1* and *FTL* genes were detected by transcriptome sequencing. The data are shown as mean ± SEM (n=3, \**p*< 0.01; $ *p*< 0.01).

**Figure S9.**
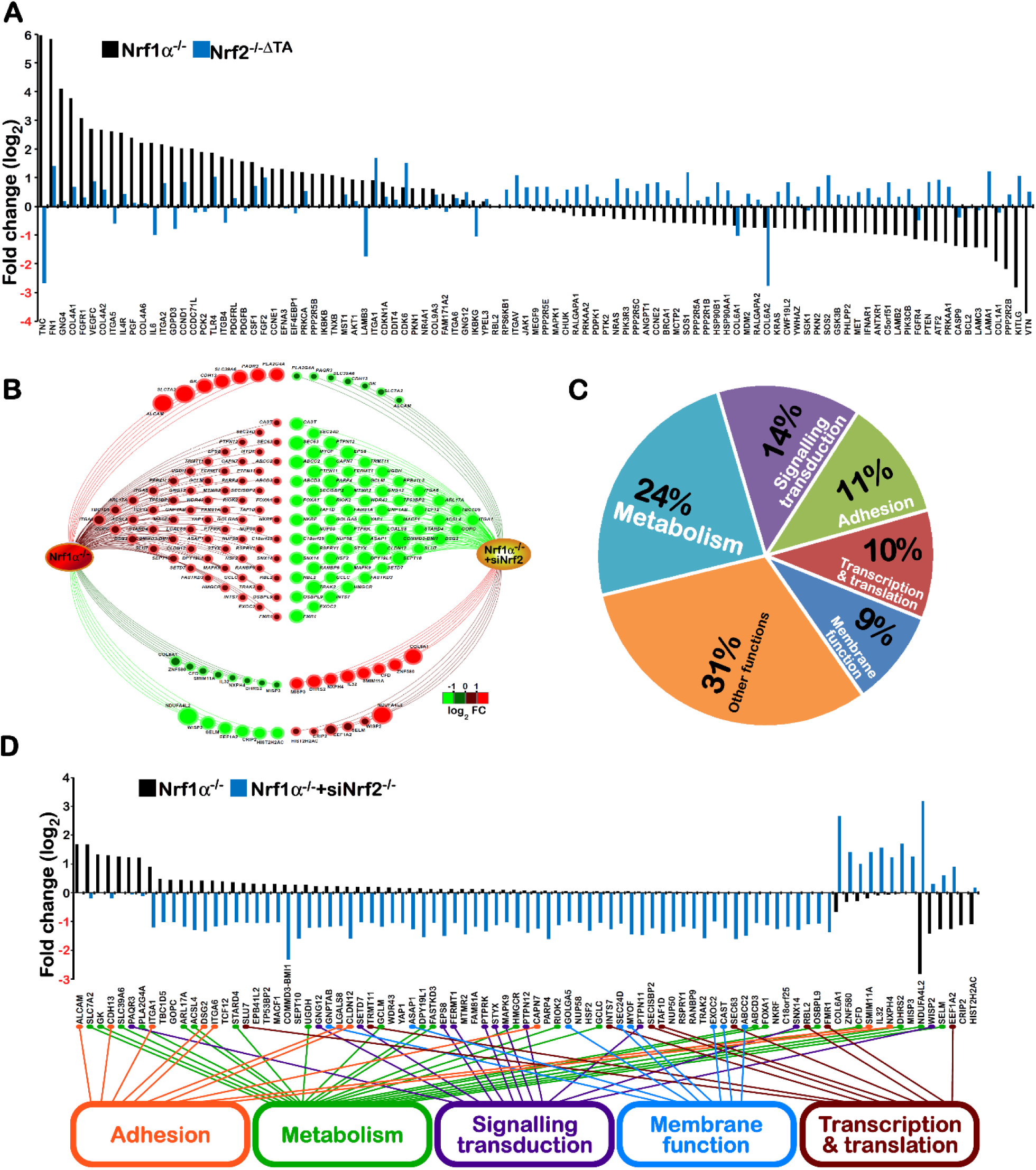
Opposite changes in DEGs measured from transcriptome in distinct cell lines. (A) Significant differences in the indicated DEGs responsible for PTEN-directed PI3K-AKT signaling pathways (also shown in Figure 7B & 7C) in between *Nrf1α^−/−^* and *Nrf2^−/−ΔTA^* cell lines are shown graphically, after normalization to relevant values measured from *Nrf1/2^+/+^* cells by transcriptome sequencing (n=3). (B to D) Opposite alterations in DEGs in between *Nrf1α^−/−^* and *Nrf1α^−/−^*+siNrf2 cell lines after being normalized to those in *Nrf1/2^+/^* cells are shown in different ways. The major functions of these genes are also classified.

**Figure S10.**
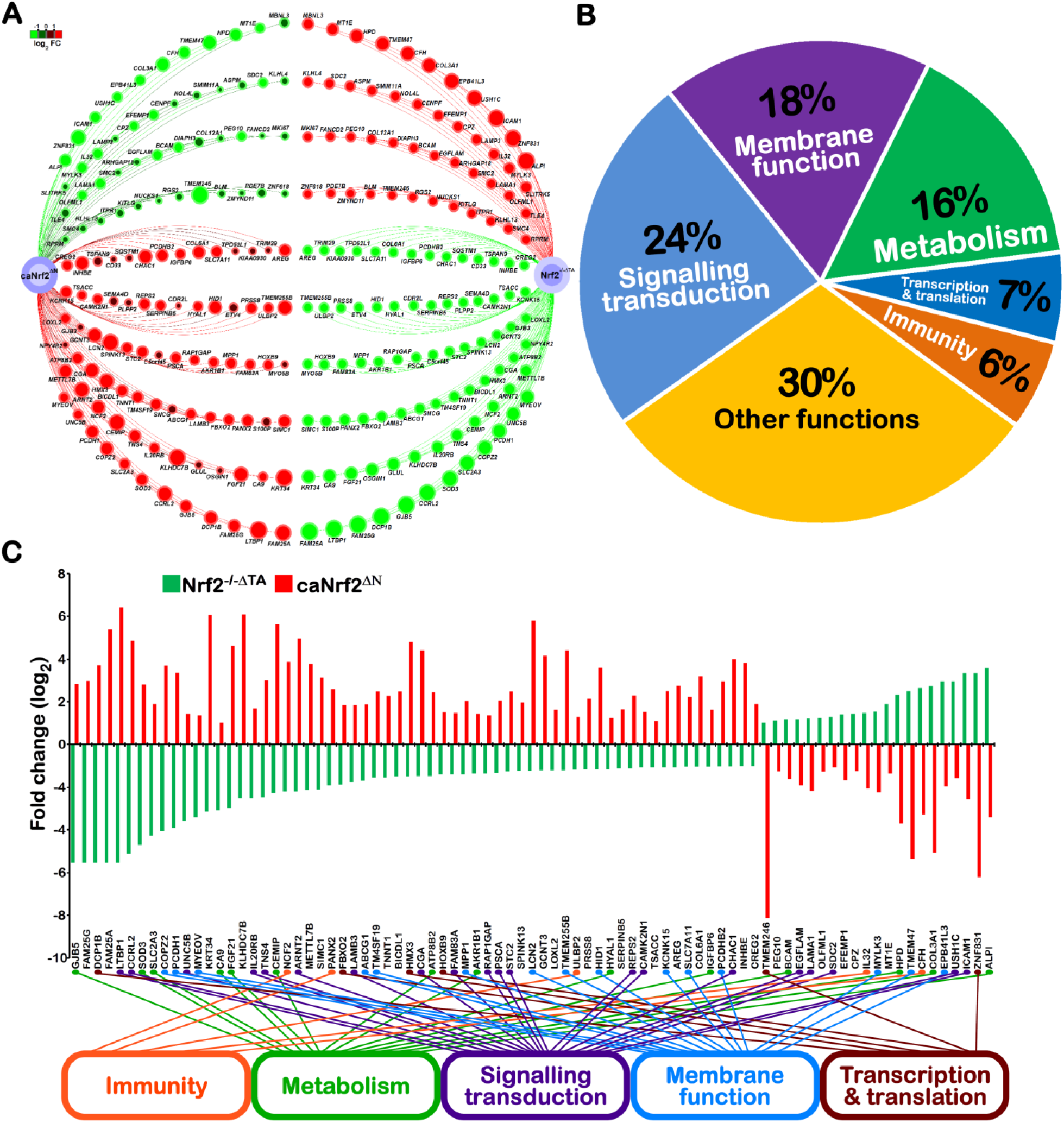
Opposite alterations in DEGs measured from transcriptome in *Nrf2^−/−ΔTA^* and *caNrf2^ΔN^* cells. These genes display opposite trends in their expression levels in between *Nrf2^−/−ΔTA^* and *caNrf2^ΔN^*, after normalization to relevant values measured from *Nrf1/2^+/+^* cells by transcriptome sequencing (n=3). The major functions of these genes are also classified.

**Table S1.**
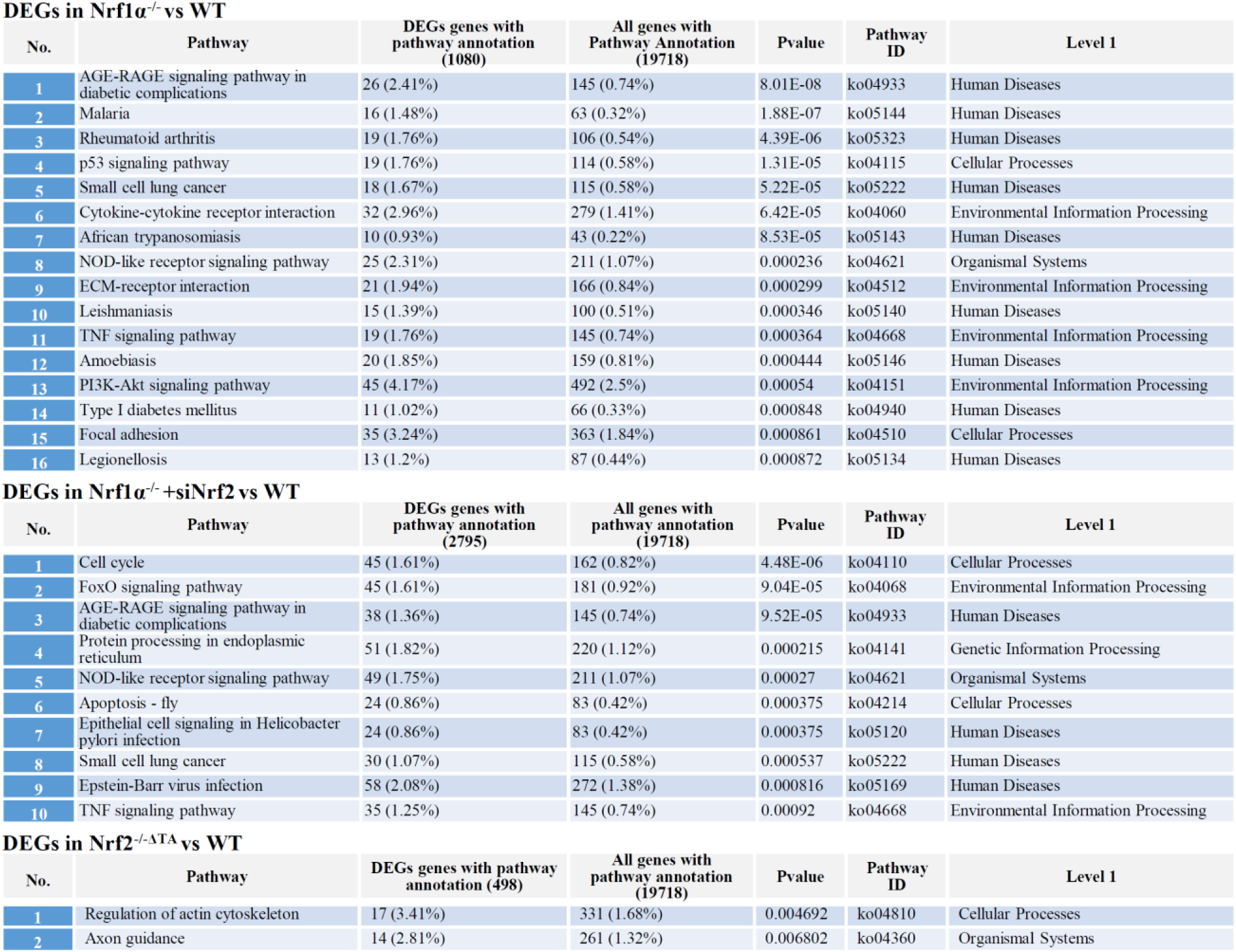
KEGG pathway enrichment analysis of DEGs

DEGs in *Nrf1* $\alpha^{-/-}$ vs WT
| No. | Pathway | DEGs genes with pathway annotation (1080) | All genes with Pathway Annotation (19718) | Pvalue | Pathway ID | Level 1 |
| --- | --- | --- | --- | --- | --- | --- |
| 1 | AGE-RAGE signaling pathway in diabetic complications | 26 (2.41%) | 145 (0.74%) | 8.01E-08 | ko04933 | Human Diseases |
| 2 | Malaria | 16 (1.48%) | 63 (0.32%) | 1.88E-07 | ko05144 | Human Diseases |
| 3 | Rheumatoid arthritis | 19 (1.76%) | 106 (0.54%) | 4.39E-06 | ko05323 | Human Diseases |
| 4 | p53 signaling pathway | 19 (1.76%) | 114 (0.58%) | 1.31E-05 | ko04115 | Cellular Processes |
| 5 | Small cell lung cancer | 18 (1.67%) | 115 (0.58%) | 5.22E-05 | ko05222 | Human Diseases |
| 6 | Cytokine-cytokine receptor interaction | 32 (2.96%) | 279 (1.41%) | 6.42E-05 | ko04060 | Environmental Information Processing |
| 7 | African trypanosomiasis | 10 (0.93%) | 43 (0.22%) | 8.53E-05 | ko05143 | Human Diseases |
| 8 | NOD-like receptor signaling pathway | 25 (2.31%) | 211 (1.07%) | 0.000236 | ko04621 | Organismal Systems |
| 9 | ECM-receptor interaction | 21 (1.94%) | 166 (0.84%) | 0.000299 | ko04512 | Environmental Information Processing |
| 10 | Leishmaniasis | 15 (1.39%) | 100 (0.51%) | 0.000346 | ko05140 | Human Diseases |
| 11 | TNF signaling pathway | 19 (1.76%) | 145 (0.74%) | 0.000364 | ko04668 | Environmental Information Processing |
| 12 | Amoebiasis | 20 (1.85%) | 159 (0.81%) | 0.000444 | ko05146 | Human Diseases |
| 13 | PI3K-Akt signaling pathway | 45 (4.17%) | 492 (2.5%) | 0.00054 | ko04151 | Environmental Information Processing |
| 14 | Type I diabetes mellitus | 11 (1.02%) | 66 (0.33%) | 0.000848 | ko04940 | Human Diseases |
| 15 | Focal adhesion | 35 (3.24%) | 363 (1.84%) | 0.000861 | ko04510 | Cellular Processes |
| 16 | Legionellosis | 13 (1.2%) | 87 (0.44%) | 0.000872 | ko05134 | Human Diseases |

DEGs in *Nrf1* $\alpha^{-/-}$ +siNrf2 vs WT
| No. | Pathway | DEGs genes with pathway annotation (2795) | All genes with pathway annotation (19718) | Pvalue | Pathway ID | Level 1 |
| --- | --- | --- | --- | --- | --- | --- |
| 1 | Cell cycle | 45 (1.61%) | 162 (0.82%) | 4.48E-06 | ko04110 | Cellular Processes |
| 2 | FoxO signaling pathway | 45 (1.61%) | 181 (0.92%) | 9.04E-05 | ko04068 | Environmental Information Processing |
| 3 | AGE-RAGE signaling pathway in diabetic complications | 38 (1.36%) | 145 (0.74%) | 9.52E-05 | ko04933 | Human Diseases |
| 4 | Protein processing in endoplasmic reticulum | 51 (1.82%) | 220 (1.12%) | 0.000215 | ko04141 | Genetic Information Processing |
| 5 | NOD-like receptor signaling pathway | 49 (1.75%) | 211 (1.07%) | 0.00027 | ko04621 | Organismal Systems |
| 6 | Apoptosis - fly | 24 (0.86%) | 83 (0.42%) | 0.000375 | ko04214 | Cellular Processes |
| 7 | Epithelial cell signaling in Helicobacter pylori infection | 24 (0.86%) | 83 (0.42%) | 0.000375 | ko05120 | Human Diseases |
| 8 | Small cell lung cancer | 30 (1.07%) | 115 (0.58%) | 0.000537 | ko05222 | Human Diseases |
| 9 | Epstein-Barr virus infection | 58 (2.08%) | 272 (1.38%) | 0.000816 | ko05169 | Human Diseases |
| 10 | TNF signaling pathway | 35 (1.25%) | 145 (0.74%) | 0.00092 | ko04668 | Environmental Information Processing |

Table S1. KEGG pathway enrichment analysis of DEGs
| No. | Pathway | DEGs genes with pathway annotation (498) | All genes with pathway annotation (19718) | Pvalue | Pathway ID | Level 1 |
| --- | --- | --- | --- | --- | --- | --- |
| 1 | Regulation of actin cytoskeleton | 17 (3.41%) | 331 (1.68%) | 0.004692 | ko04810 | Cellular Processes |
| 2 | Axon guidance | 14 (2.81%) | 261 (1.32%) | 0.006802 | ko04360 | Organismal Systems |

